# Light-induced assembly and repeatable actuation in Ca^2+^-driven chemomechanical protein networks

**DOI:** 10.1101/2025.03.03.641304

**Authors:** Xiangting Lei, Carlos Floyd, Laura Casas-Ferrer, Tuhin Chakrabortty, Nithesh Chandrasekharan, Aaron R. Dinner, Scott Coyle, Jerry Honts, Saad Bhamla

## Abstract

Programming rapid, repeatable motions in soft materials has remained a challenge in active matter and biomimetic design. Here, we present a light-controlled chemomechanical network based on *Tetrahymena thermophila* calcium-binding protein 2 (Tcb2), a Ca^2+^-sensitive contractile protein. These networks—driven by Ca^2+^-triggered structural rearrangements—exhibit dynamic self-assembly, spatiotemporal growth, and contraction rates comparable to actomyosin systems. By coupling light-sensitive chelators for optically triggered Ca^2+^ release, we achieve precise growth and repeatable mechanical contractility of Tcb2 networks, revealing emergent phenomena such as boundary-localized active regions and density gradient-driven reversals in motion. A coupled reaction-diffusion and elastic model explains these dynamics, highlighting the interplay between chemical network assembly and mechanical response. We further demonstrate active transport of particles via network-mediated forces *in vitro* and implement reinforcement learning to program seconds-scale spatiotemporal actuation *in silico*. These results establish a platform for designing responsive active materials with rapid chemomechanical dynamics and tunable optical control, with applications in synthetic cells, sub-cellular force generation, and programmable biomaterials.

## I. INTRODUCTION

In recent years there has been significant research focus on chemomechanical materials in which chemical or optical cues drive mechanical and structural changes [1]. Both purely synthetic and bio-inspired materials have been developed with design goals such as reversible self-assembly, on-demand force generation, and spatial programmability. For example, synthetic approaches have coupled pattern-forming chemical reactions with hydrogels to induce periodic swelling and force generation [2– 6]; however, such motions are often slow and difficult to direct, as they rely on intrinsic chemical oscillations. Light-driven strategies, including the covalent assembly of polymer networks [7, 8], offer greater external control but tend to form long-lived structures unsuitable for dynamic force production. Reversible self-assembly has also been demonstrated in polymeric and liquid crystalline materials [9–12], though these systems typically respond to uniform chemical triggers and lack fine spatial control.

As an alternative to purely synthetic systems, bioderived chemomechanical materials offer evolved platforms for non-equilibrium force generation. These include cellular and bacterial suspensions [13, 14], cilia carpets [15, 16], and, notably, assemblies of cytoskeletal filaments and motor proteins like actomyosin or microtubule-kinesin mixtures [17–27]. Recent bioengineering of light-sensitive molecular motors enables dynamical control over force generation using spatiotemporal light patterning in cytoskeletal materials [22, 23, 28– 36]. However, light-controllable cytoskeletally-derived materials face limitations such as flows constrained to filament orientations and reliance on complex molecular bioengineering. To our knowledge no synthetic or bioderived material has yet demonstrated the simultaneous combination of self-assembly triggered by spatial light patterning, on-demand repeatable and reorientable force generation, fast (seconds) assembly rates, a bioorthogonal energy source, and minimal fabrication complexity. A material with these properties would be an excellent candidate for application in synthetic cell biology, in which light-activated rapid force-generation could be used to guide intracellular dynamics.

While most biomechanical processes are powered by a direct coupling between hydrolysis of adenosine triphosphate (ATP) and molecular motion, nature has also developed alternative paradigms for generating biological motion [37– In various heterotrich ciliates, contractile motion arises from specialized Ca^2+^-binding protein assemblies which in different organisms have been termed myonemes [37, 40] and spasmonemes [39, 41]. Our comparative genomics analyses reveal sequence similarities among these various contractile proteins (SI I A). Unlike traditional cytoskeletal systems, these ciliate contractile systems generate motion through direct, Ca^2+^-induced structural rearrangements of the polymers [42], rather than through relative sliding of filament pairs. These systems generate repeatable forces for ultrafast *in vivo* contractions—the fastest known motions in biology [37]—and are roughly ten or more times faster than actomyosin-based networks (see Discussion). As a result, these Ca^2+^-driven protein systems hold promise for deriving new force-generating biomaterials with faster dynamics than traditional cytoskeletal components.

To begin theoretically understanding their dynamics, in Ref. 43 we modeled ciliates’ myoneme-based contraction *in vivo*, which is triggered by a propagating wave of Ca^2+^ in the cytosol. However, despite this preliminary modeling and related studies [44–49], our quantitative understanding of these naturally evolved protein networks remains limited compared to cytoskeletal systems. Furthermore, experimental methods akin to those used for reconstituting and controlling cytoskeletal materials *in vitro* using spatiotemporal light patterning are currently unavailable for Ca^2+^-triggered proteins. Finally, while theoretical and computational advances have facilitated the design of control strategies for light-triggered cytoskeletal systems [50–57], such methods are currently lacking from our toolkit to study Ca^2+^-triggered protein dynamics.

Here, we purify and reconstitute polymeric networks of *Tetrahymena* calcium-binding protein 2 (Tcb2), a Ca^2+^-sensitive contractile protein from the ciliate *Tetrahymena thermophila* [58, 59]. Although its physiological function is not fully known, it is thought to play a role in Ca^2+^-dependent processes such as ciliary movement, pronuclear exchange, and signaling in the cortex. [58, 60, 61]. This protein is homologous to the Ca^2+^-binding proteins that underlie ultrafast contraction in other ciliates, and we focus on it here because there are available molecular tools for its expression and it does not require cosynthesis with a scaffold protein (SI I A).

We develop an experimental methodology to reconstitute a contractile Tcb2 protein network *in vitro* and spatiotemporally control it using light-triggered Ca^2+^ release by diffusing chelators. We show that this system exhibits rich dynamical behaviors, including a Ca^2+^-driven active region (CAR) near the boundary and a reversal of contractility direction, from radially inward to outward. We explain these dynamics by introducing a continuum model of an overdamped elastic solid with inhomogeneous density and an elastic rest length dependent on the local Ca^2+^ concentration, which we couple with a reaction-diffusion-based description of the chemical components. Additionally, we demonstrate preliminary control over the transport of external particles by the Tcb2 networks *in vitro*, as well as fine-grained control over network contraction *in silico* using reinforcement learning. We conclude by discussing its potential applications in synthetic biology. This system represents a new biologically derived soft material for producing repeatable and rapidly responsive forces at subcellular scales. We conclude by discussing its potential applications in synthetic biology.

## II. RESULTS

### A. Spatiotemporal light patterning controls Tcb2 protein network dynamics

We first aim to develop an experimental platform for studying Ca^2+^-driven contractile proteins *in vitro*. The ultrafast contractile motion of ciliates involves assemblies such as myonemes, and it is driven by rapid waves of Ca^2+^ released from internal stores such as the endoplasmic reticulum (ER) [37, 43, 49]. After the organism contracts, ATP is hydrolyzed to pump Ca^2+^ back into the ER to reset the process. Several ciliate species in the “atlas” of ultrafast cells exhibit high-speed contraction [62], but they are not model biological systems and, hence, we currently lack sufficient experimental protocols to study them using protein reconstitution. However, homologs of centrin (one of the main components of myonemes) which also interact with Ca^2+^, though not observed to produce whole-cell ultrafast cellular contractions, have been identified in other model ciliates (SI I A). Here, we focus on the homologous protein Tcb2 from *Tetrahymena thermophila*, which we show exhibits a marked structural response upon binding to Ca^2+^ [42], making it a valuable model for studying Ca^2+^-driven contractile proteins. Using available molecular tools we express this protein in ample quantities, allowing for straightforward reconstitution *in vitro*.

Tcb2 is a 25 kDa Ca^2+^-binding protein found in the cortical layer of *Tetrahymena* [60]. Tcb2 has four EF-hand domains with putative Ca^2+^-binding sites [42, 58, 63]. In the absence of a solved crystal structure, we use AlphaFold3 [64] to predict the structure of Tcb2 monomers and their conformational changes after binding Ca^2+^. The predicted conformational changes are consistent with previous NMR data [42] (SI I A), and these conformational changes underscore the potential of using this protein in a chemomechanical material with contractile dynamics.

We clone DNA encoding a synthetic Tcb2 gene that was codon-optimized for bacterial expression into an expression plasmid. We then express and purify the protein from *E. coli* as in Ref. 65 (Methods and SI I C). We observe that purified Tcb2 alone, without additional scaffolding proteins, readily forms fibrous protein networks in response to Ca^2+^ ions and subsequently contracts along the network boundary (SI Video Part 1, Section I). Electron microscopy reveals fine, irregular filaments around 250-500 nm at low Ca^2+^ concentrations (Fig. 1a). At higher Ca^2+^ concentrations we observe denser and largescale networks of filamentous protein as shown by micrographs with fluorescent mCherry-tagged Tcb2 at a larger spatial scale (Fig. 1b). This propensity to self-assemble into a cortex-like [28] network distinguishes Tcb2 from other Ca^2+^-binding proteins like calmodulin [66].

**FIG. 1.**
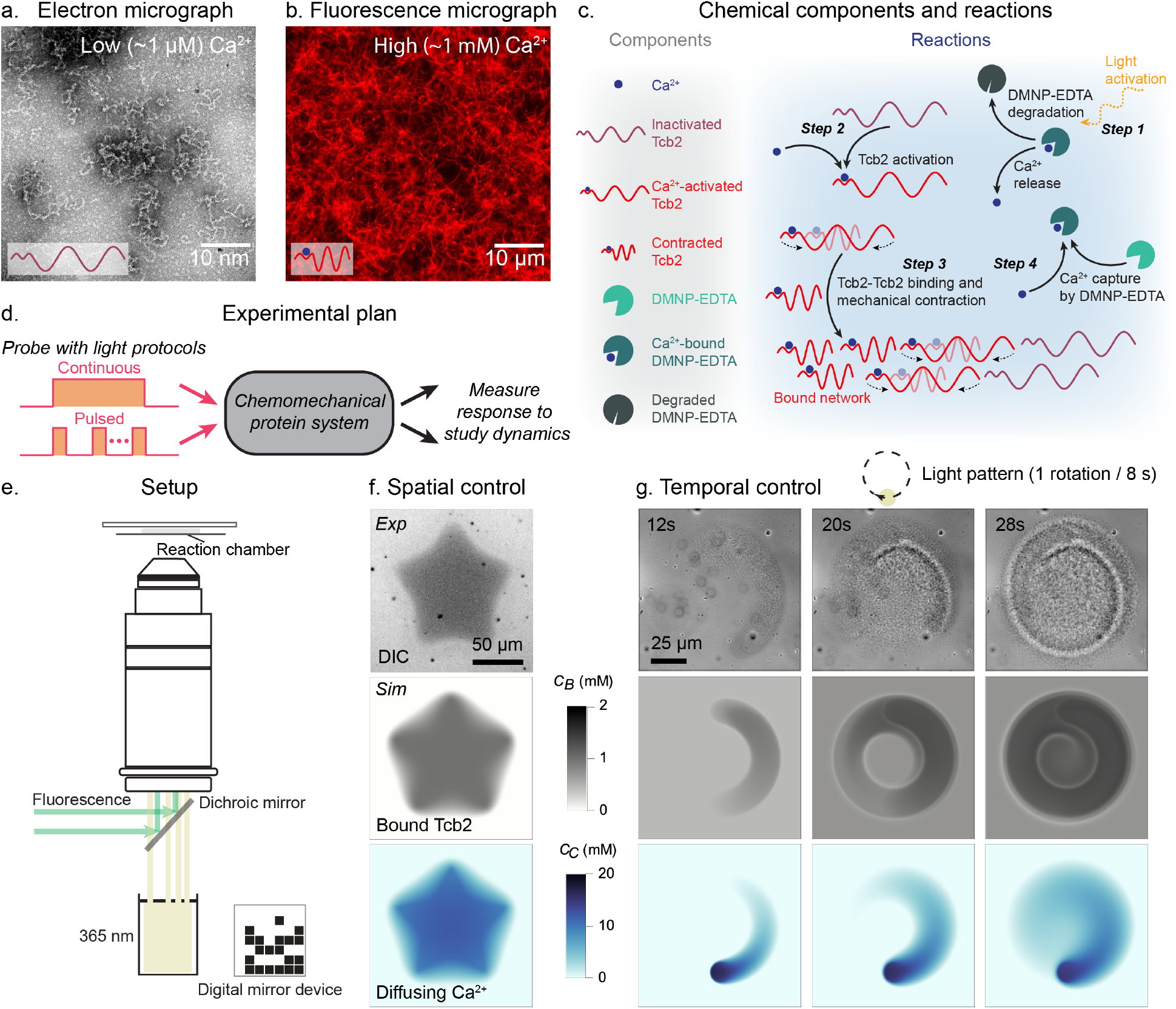
Experimental control of Tcb2 network assembly. (a) Tcb2 filaments observed at ∼ 1 *µ*M Ca^2+^ concentrations using electron microscopy, showing typical filament lengths of approximately 250-500 nm. (b) Tcb2/Tcb2-mCherry filaments in the presence of ∼ 1 mM Ca^2+^ observed via fluorescence microscopy. (c) Chemical components and reactions involving Tcb2 and Ca^2+^, including light activation, DMNP-EDTA binding/unbinding, and Tcb2 binding/unbinding. (d) Schematic illustration of the experimental plan in this paper. (e) Schematic illustration of the experimental setup. (f) Tcb2 network formation, imaged using DIC, for a star-shaped DMD pattern. Simulated bound Tcb2 (*C*_*B*_) and diffusing Ca^2+^ (*C*_*C*_) concentrations are shown below. (g) Dynamic Tcb2 network formation using a cyclic light pattern, with accompanying simulations of bound Tcb2 and diffusing Ca^2+^ concentrations. The light moves in a circle at 0.125 Hz, shown at different cycles.

We employ an experimental methodology that releases Ca^2+^ in a spatiotemporally controlled manner using photolyzable Ca^2+^ chelators. The DMNP-EDTA Ca^2+^ chelator has a photolabile bond, which, when severed upon illumination by 365 nm light, releases Ca^2+^ in milliseconds [67] (SI I D). This complex has been used extensively in optogenetics [68–70], and we adapt it here to achieve precise optical control of Ca^2+^ release in Tcb2 solutions. We note that in contrast to the situation *in vivo*, in which energy is provided to the system via ATP-driven pumping of Ca^2+^ ions and their subsequent rapid release from intracellular stores, here energy is provided by photons which cleave Ca^2+^ chelators. These are different implementations of the same underlying logic: Ca^2+^ storage through intracellular stores or DMNP-EDTA binding, and release through calcium channel opening or photocleavage of chelating groups. In Fig. 1c we schematically show the chemical components and reactions of the system, including chelators, Ca^2+^ ions, Tcb2 proteins, and externally applied light. Our goal in this work is to use canonical light protocols, such as step functions and periodic pulses, to probe the unknown dynamical properties of this material (Fig. 1d) [71].

To spatiotemporally control DMNP-EDTA photolysis we use a custom optical setup, integrating a digital micromirror device (DMD) pattern illuminator into a microscope (SI I D). This setup uses dual optical pathways via a multi-port illuminator, enabling both fluorescence and 365 nm illumination (Fig. 1e). This system provides sub-micrometer spatial resolution and millisecond temporal precision [72]. To illustrate dynamical control using this system, we project specific spatial and temporal patterns of illumination onto the Tcb2 solution. We illuminate a sharp star shape using the DMD and, within one second, observe a corresponding pattern in the differential interference contrast (DIC) images of the Tcb2 solution, indicating the formation of a Ca^2+^-bound Tcb2 network at the corresponding location (Fig. 1f). These patterns can also be programmed into time series, enabling more complex manipulation of the network. We demonstrate this using a circular illumination pattern traveling in a circular trajectory at 0.125 Hz (Fig. 1g and SI Video Part 1 Section II). We note that the timescale of the observed assembly of Tcb2 networks is seconds. This is faster than those observed in actomyosin or microtubule-kinesin systems, which typically require minutes to hours to assemble and achieve similar dynamics [31, 73].

We augment this experimental setup with a continuum reaction-diffusion model of the chemical system illustrated in Fig. 1c (Methods and the SI I E). Under reasonable estimates for the unknown model parameters we observe qualitative agreement between the predictions of the model and key experimentally observed features of Tcb2 network dynamics (Figs. 1f,g). We elaborate on details of this model as they become relevant below.

### B. A Ca^2+^-driven active region results from the contraction dynamics of diffusion-limited growth

Having established the experimental and computational setup, we next seek to characterize the chemomechanical dynamics of the Tcb2 solution systematically using standard light activation protocols such as step functions and pulses (Fig. 1d). We first characterize the growth dynamics of the Tcb2 protein network by projecting a sustained circular light pattern on the solution (SI Video Part 1 Section II). Unless otherwise indicated, all results in this paper use a 75 *µ*m diameter for the illuminated circle, but in SI II A we show results using different illumination diameters. During 100 s of illumination we observe that first, upon illumination, Tcb2 quickly forms a connected network within the illuminated region (Figs. 2a,b), where the Ca^2+^ concentration increases suddenly due to chelator photolysis. Over time the network grows radially outward (Fig. 2a at 100 s and Fig. 2b). In SI Fig. S11 we plot the detected area of the network as a function of time, finding that it scales approximately linearly, which indicates diffusion-limited growth dynamics. The growth rate also increases with the diameter of the illumination region. We additionally observe outward diffusion of Ca^2+^ using rhodamine-2, a Ca^2+^ sensitive dye (SI Fig. S20). In Ref. 74 we further develop calibrated rhodamine assays to quantify growth dynamics of Tcb2 networks.

**FIG. 2.**
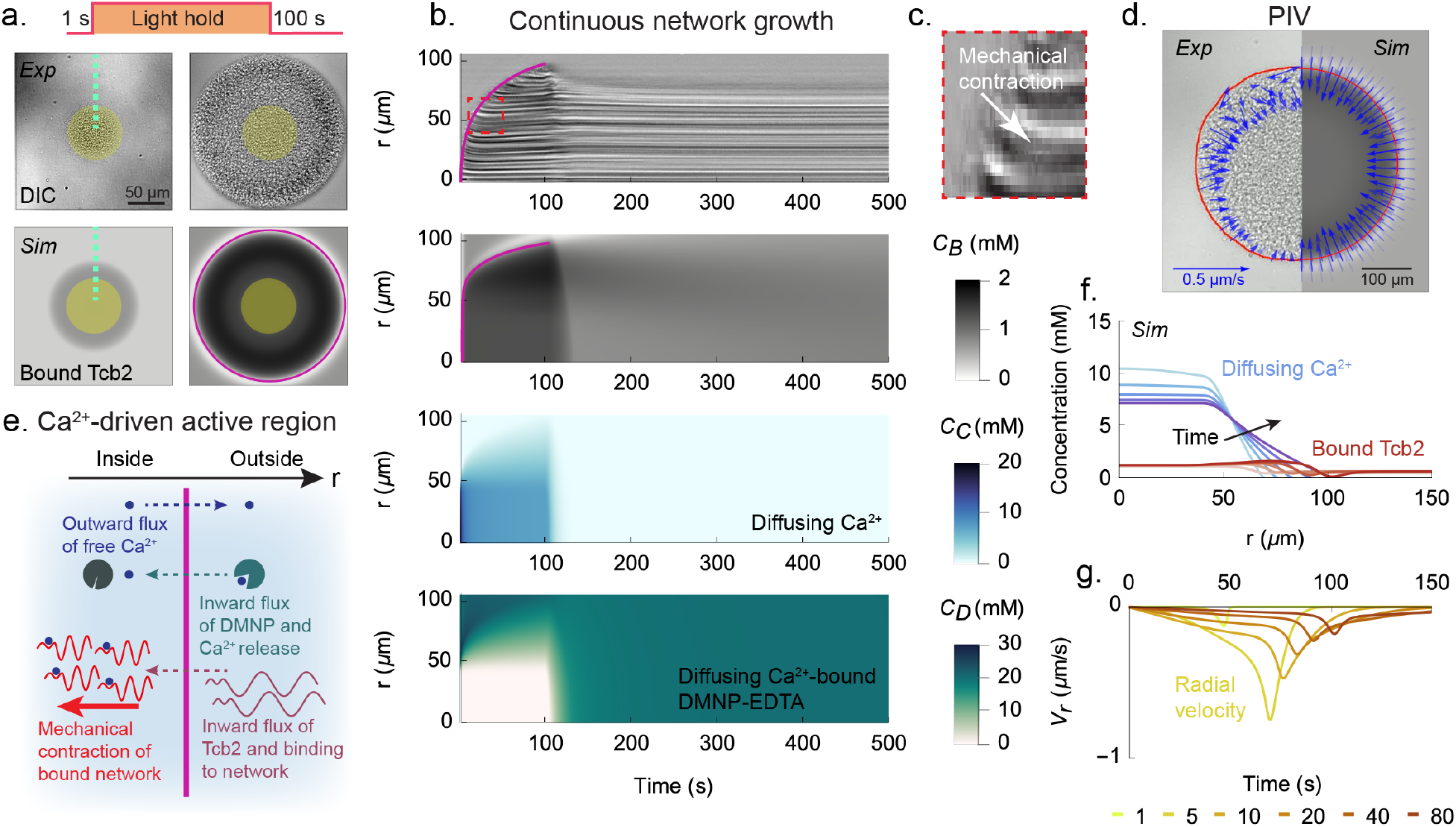
Continuous growth of the network produces an active region. (a) *Top row* : DIC images of Tcb2 network with circular illumination at 1 s and 100 s in a continuous light protocol, using an illumination diameter of 75 *µ*m. Experiments are repeated three times. *Bottom row* : Corresponding Tcb2 simulation images. (b) *First row:* Kymograph constructed from DIC images for the continuous light experiment in panel a, taken along the green dashed line. *Second row* : Simulated kymograph, showing bound Tcb2 concentration as a function of radial position. *Third row* : Same as second row, except showing diffusing Ca^2+^ concentration. *Fourth row* : Same as second row, except showing Ca^2+^-DMNP-EDTA concentration. (c) Magnified view of contracting region (red dashed box in the top row of panel b). (d) Experimental PIV field and corresponding simulation image at 100 *s*. Minor angular asymmetries occasionally visible in experimental PIV arise from stochastic experimental non-uniformities and are not reproducible across replicates. (e) Schematic illustration of the CAR. (f) Concentration of Ca^2+^ (blue) and bound Tcb2 (red) over time. The colors range from light to dark and match the times in the next panel. (g) Radial velocity profiles *V*_*r*_ (*r, t*) of the network over time.

We observe that in the growing network the region near the boundary is mechanically active. The newly formed network at the boundary contracts radially inward, as shown by a kymograph (Figs. 2b,c) and particle image velocimetry (PIV, Fig. 2d). We refer to this boundary as a Ca^2+^-driven active region (CAR, Fig. 2e) and summarize the processes within the CAR as follows. Chemically, excess Ca^2+^ ions released from photolyzed DMNP-EDTA that have not yet bound to Tcb2 diffuse outward, while unphotolyzed Ca^2+^-DMNP-EDTA complexes diffuse inward, both following their respective concentration gradients. As these complexes reach the illuminated zone they photolyze and release additional Ca^2+^, sustaining a loop that fuels network growth by continuously supplying Ca^2+^ from the inside. Tcb2 proteins diffuse inward at the CAR, where they bind to both other Tcb2 proteins and available Ca^2+^, joining the growing network. Based on molecular size, we estimate the diffusion constant of Tcb2 monomers to be 10 *µ*m^2^*/s*, and we show in SI Fig. S27 that accounting for Tcb2 diffusion is necessary to explain the network’s growth dynamics. Upon Ca^2+^ binding, Tcb2’s rest length shortens, prompting rapid, inward contraction. Since the network’s interior has reached mechanical equilibrium, contraction occurs mainly at the periphery.

To describe the contractile dynamics of this system computationally, we augment the chemical variables in the continuum model with a two-dimensional displacement field that obeys the dynamics of an overdamped linearly elastic gel [75]. We couple this mechanical representation to the chemical fields in two ways. First, because in a typical Ashby plot [76] the elastic moduli of materials increase with their density, we make the elastic moduli of the gel proportional to the local concentration of bound Tcb2. We explore non-linear dependencies between density and stiffness in SI Fig. S7, showing a qualitative insensitivity to this modeling choice. Second, following our previous model [43] for *in vivo* ultrafast myoneme contraction in ciliates, we decrease the local rest length of the material linearly with the fraction of bound Tcb2 (SI I E). This decrease of local rest length, which can be viewed as the equilibrium end-to-end length of the Tcb2 monomers, reflects the conformational change in Tcb2 monomers suggested by NMR data and AlphaFold3 predictions mentioned above. We assume that this conformational change underlies the mechanical contractility of the macroscopic network although at a coarse-grained modeling level the molecular details of this contraction is not critical [42] (SI I A).

Using the model, which allows us to spatially resolve all of the chemical components (SI Fig. S18), we observe that the boundary of the Tcb2 network closely follows the leading edge of the diffusing Ca^2+^ profile as shown in Fig. 2f. This supports the picture that the outward growth of the Tcb2 network is driven by the diffusion of Ca^2+^ out from the illuminated region and of Tcb2 into the bound network region. The model reproduces the CAR, such that the contracting radial velocity peak is located near the periphery of the Tcb2 network, moving radially outward as the network grows (Fig. 2g). We further find that, to prevent the growth from stalling at finite radius due to the balance of reaction-diffusion fluxes, it is necessary to account for the degradation of DMNP-EDTA chelators in the model (SI Fig. S22).

After the light is turned off in the experiments, Ca^2+^ rapidly unbinds from the network and is resequestered by the available DMNP-EDTA chelator. Upon unbinding of Ca^2+^ and re-elongation of the Tcb2 rest lengths (via conformational change of Tcb2 monomers back to the Ca^2+^-free state with a longer equilibrium end-to-end distance), the network quickly relaxes from its contracted state and expands outward (SI Video Part 1 Section III). It then slowly dissolves via the unbinding of Tcb2 from other Tcb2 proteins. The slow timescale of chemical growth and dissolution is distinct from the fast timescale of mechanical contraction and expansion (SI Fig. S21). A similar interplay between rapid mechanical responses and slower chemical reorganization has been observed in simulations of actomyosin networks under tensile force [77]. We next aim to leverage these two contrasting timescales to overcome the size limitation which arises from the actively contracting region advancing outward in a single sweep and leaving most of the internal region mechanically equilibrated in its path.

### C. Temporally pulsed illumination yields a larger and faster Ca^2+^-driven active region

In an effort to “recharge” the system, we next investigate the use of a pulsed, or ratcheting, illumination protocol: a 1 s flash of illumination followed by 29 s of darkness, repeated in cycles. During the initial few cycles, the network formed in the 1 s flash mostly dissolves back into solution during the light-off phase (Fig. 3a at 1 s and 29 s). After these initial cycles, a persistent network begins to form and then grow in size with each subsequent pulse (Fig. 3a at 10^th^ and 30^th^ pulse and Fig. 3b). In SI II A, II I, and II J, we study the dependence of growth dynamics on illumination size, pulse frequency, and DMNP-EDTA concentration. In SI II B and SI Video Part 2 Section VI, we demonstrate sustained contractile responses over ∼ 150 cycles without significant performance degradation. The system maintains consistent contraction speeds of ∼ 0.4 *µ*m/s across illumination diameters ranging from 50–100 *µ*m, highlighting its potential for applications requiring repeated mechanical actuation.

**FIG. 3.**
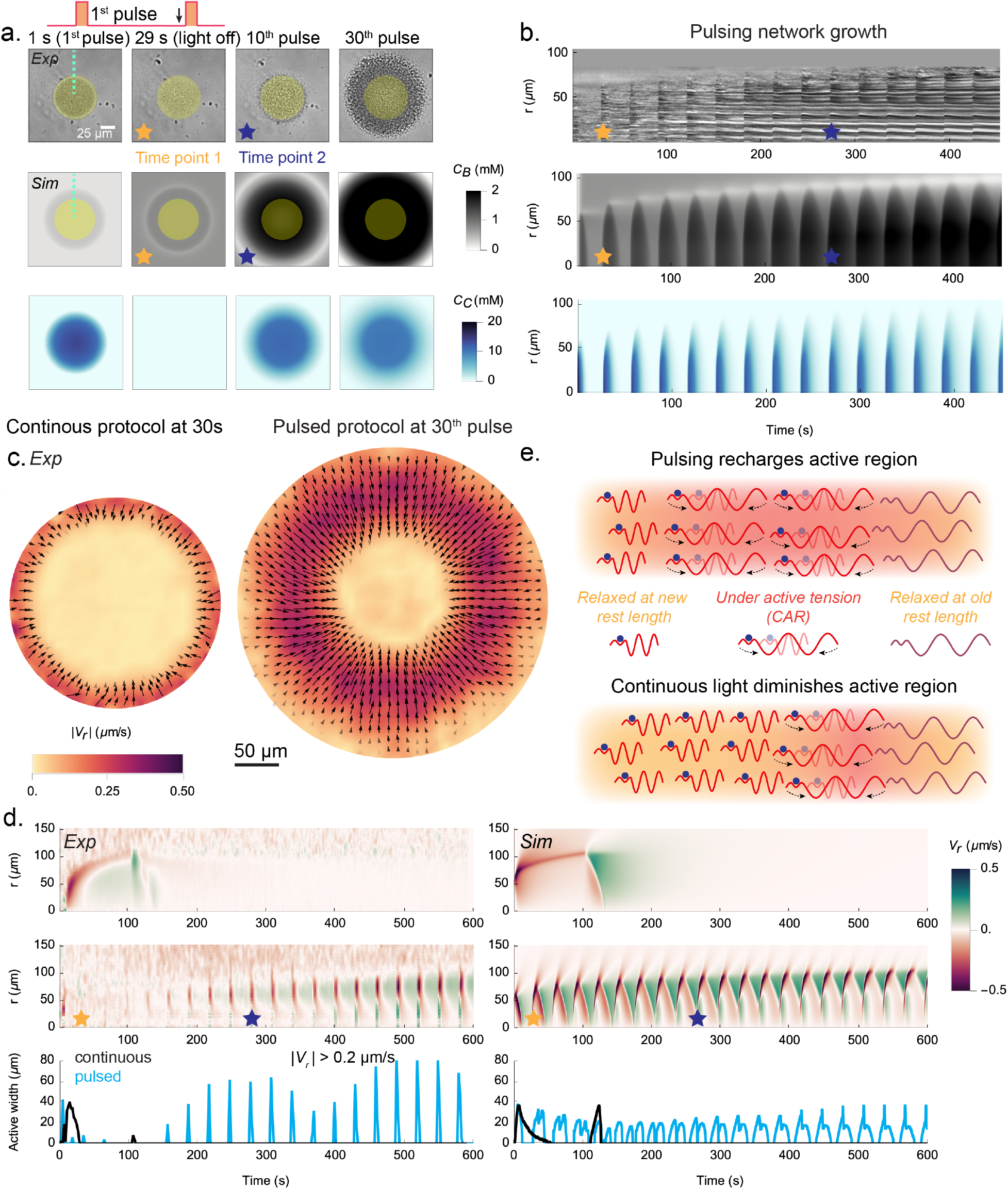
Pulsed light allows the CAR to recharge. (a) DIC images of a pulsed illumination experiment at 0 s, 29 s (right before the 2^nd^ pulse), the 10^th^ pulse, and the 30^th^ pulse. Corresponding simulation images of the bound Tcb2 and diffusing Ca^2+^ profiles are shown below. The blue and yellow stars indicate the same pulses (time points) throughout the figure. The dashed green line indicates the region used to generate the kymograph shown in panel (b). (b) Kymographs of the DIC images shown above corresponding simulation results. (c) PIV of continuous light at 30 s (left) and the 30^th^ cycle of the pulsed protocol, with arrows indicating the direction of network motion. The background color represents velocity magnitude and the arrows represent the local velocity vector. (d) *Top row* : Experimental and simulated velocity kymographs for the continuous protocol. *Middle row* : Experimental and simulated velocity kymographs for the pulsed protocol. *Bottom row* : Experimental and simulated plots of the CAR width, defined as the radial extent of the system for which |*V*_*r*_ |*>* 0.2 *µ*m/s, as a function of time for the continuous and pulsed protocols. (e) Schematic illustration of the Tcb2 recharging mechanism. See Fig. 1c for a legend of the molecular components. Experiments are repeated three times.

In Fig. 3c we show PIV measurements comparing the continuous illumination protocol, evaluated at 30 s, with the pulsed protocol, evaluated at the 30^th^ cycle (i.e., the same total time of illumination). In the continuous protocol the CAR is relatively slender and has low average radial velocity (width: ~13 *µ*m, 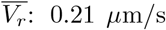), while in the pulsed protocol the CAR is significantly larger and faster (width: ~62 *µ*m, 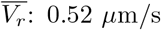); see SI Fig. S23. Importantly, pulsing also causes the CAR to remain large over many repeated cycles (middle row Fig. 3d). To quantify this we measure the width of the radial domain for which the norm of the radial velocity exceeds 0.2 *µ*m/s (chosen as an arbitrary cutoff value), which we plot as a function of time at the bottom of Fig. 3d. These measurements demonstrate that, in contrast to the continuous protocol, pulsed light effectively recharges the system and allows for repeated generation of large contractile forces that act over nearly the whole network.

Simulations qualitatively reproduce these findings, allowing closer investigation of the recharging mechanism (Fig. 3d,e). Each Tcb2 filament has a rest length that shrinks upon binding Ca^2+^, supported by earlier NMR data [42] and consistent with structural changes suggested by AlphaFold3 (SI I A). Each filament can thus be viewed as being in one of two states with different rest lengths, depending on whether Ca^2+^ is bound to the filament. Upon Ca^2+^-binding, but before the filament contracts toward the shorter rest length, there is a build up of mechanical potential energy in the system (SI Fig. S19). The Ca^2+^-bound filaments then contract in length to reduce this energy. In the pulsed light protocol, the dark phase allows Ca^2+^ to quickly unbind from contracted filaments, restoring their longer rest lengths. The filaments then mechanically relax to these lengths, but due to the slower unbinding of Tcb2 molecules from each other the network does not fully dissolve during the dark phase. In this way the bound Tcb2 state is “ratcheted” while the mechanical state is “reset.” The extended Tcb2 filaments are then ready to contract again during the subsequent pulse of light. By contrast, in the continuous protocol the network grows steadily, binding to new Ca^2+^ at the periphery while the interior remains static and stress-free once mechanically equilibrated under the shortened rest lengths. We emphasize that these networks demonstrate rapid and repeatable mechanical force generation through rapid Ca^2+^ binding and unbinding kinetics. However, the Tcb2 network assembly itself is not rapidly reversible due to the slow dissolution process governing Tcb2-Tcb2 unbinding. The recharging mechanism in the pulsed protocol enables repeatable seconds-scale contractile dynamics, offering potential for Tcb2 networks to generate controlled micron-scale mechanical forces using light-based protocols.

### D. Spatially heterogeneous patterns of simultaneous contraction and expansion

We observe that during network growth in the pulsed protocol the system not only contracts inward but also exhibits regions that contract radially outward, as shown in the PIV images from the 10^th^ and 25^th^ pulses in Fig. 4a. This is also illustrated in Fig. 4b, which further shows that at even later times (30^th^ pulse) the contraction direction is uniformly radially inward again. This reversal of contraction direction is also observed in the simulations (Fig. 3d).

**FIG. 4.**
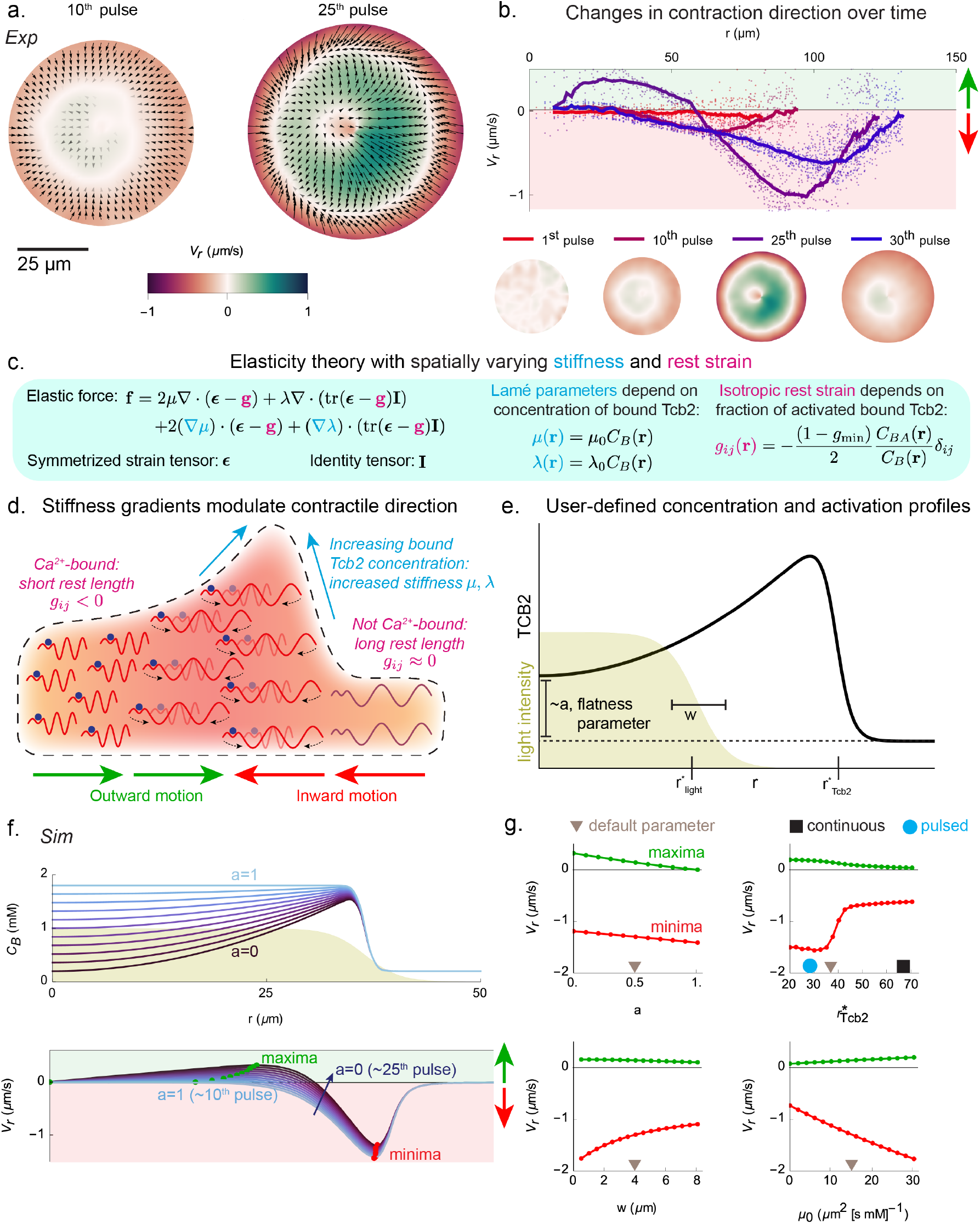
Reversal of contraction direction depends on inhomogeneous density. (a) Experimental PIV images at the 10^th^ and 25^th^ pulses. (b) *Top row* : Velocity profiles at the 1^st^, 10^th^, 25^th^, and 30^th^ pulses plotted against radial distance. *Bottom row* : Experimental PIV fields at the corresponding time points, with the same velocity scale as in panel a. We omit the arrows here for visual clarity and show only color. (c) Summary of the elastic theory used in our model (SI I E for more details). Schematic illustration of the mechanism by which contraction direction reverses due to gradients of stiffness. See Fig. 1c for a legend of the molecular components. (e) Schematic depiction of the user-defined density profile used in the remaining panels. (f) *Top row* : Illustration of the bound Tcb2 profile as the user-defined flatness parameter *a* (which is small for large boundary accumulation) is varied. *Bottom row* : Radial velocity profiles during light activation for the corresponding density profiles in the top row. Qualitative correspondences to the pulses in panel b are indicated parenthetically. (g) The maximal and minimal radial velocities (green and red dots in panel f) plotted as *a* (the flatness parameter), 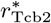 (the boundary of the Tcb2 network), *w* (the width of the light profile), and *µ*_0_ (the slope of stiffness dependence on *C*_*B*_) are varied. Default values used in the plots are shown indicated with a brown triangle. The black square and cyan circle symbols suggest correspondences to the continuous and pulsed light protocol.

What causes the change in direction of radial contraction? Analysis of the simulations reveals that a key element is an accumulation of bound Tcb2 near the periphery of the growing network, which appears as a density profile that peaks near the network boundary. In SI Figs. S26 and S27, we show peaked experimental and simulated concentration profiles and demonstrate that boundary accumulation largely depends on the diffusion of Tcb2 inward across the CAR. These new Tcb2 monomers first have a chance to bind when they encounter the network periphery, leading to a buildup of density there. The dynamical implications of these density inhomogeneities can be understood by considering the new qualitative features of our mechanical model beyond standard linear elasticity, as summarized in Fig. 4c; see SI I E for details. The density of the material is inhomogeneous, which causes the first and second Lamé parameters *λ*(**r**) and *µ*(**r**) to be inhomogeneous as well, and the instantaneous rest strain tensor **g**(**r**) is also inhomogeneous due to uneven binding of Ca^2+^ to Tcb2. These two modifications produce additional contributions to the total elastic force beyond those in a homogeneous material, and these additional forces depend on spatial gradients in the material density and bound Ca^2+^ concentration. In the presence of a peak in the density of bound Tcb2, these additional forces can overcome the inward contractile force and cause the material to move up the gradient of material density, which is radially outward for interior parts of the network (Fig. 4d).

To illustrate this picture, we perform reduced simulations in which we turn off chemical reactions, impose a user-defined concentration profile of bound Tcb2 and light-patterned Ca^2+^ activation, and simulate the dynamics of contraction. This allows us to vary the width and accumulation of the Tcb2 profile as an input. The relevant parameters of this setup are illustrated in Fig. 4e (see SI I E for details). The parameter *a* represents how much accumulation of Tcb2 there is near the boundary of the network. We find that this directly controls the positive component of the radial velocity profile, as shown in Figs. 4f,g, supporting the hypothesis that the accumulation of bound Tcb2 near the boundary underlies the reversal in contraction direction (see Fig. 2a and SI Fig. S26 for experimental observation of accumulation). We further find that as the radius of the Tcb2 network increases beyond the light illumination region, the positive component smoothly decreases to zero in magnitude. This explains why at later cycles in the pulsed protocol, when the network has grown well beyond the illumination zone, the network again contracts uniformly inward. We also find that, as the width of the light activation profile increases, weakening the gradients in rest length, the contractile velocities decrease in magnitude. Finally, increasing the proportionality constant *µ*_0_, which links the shear modulus to the bound Tcb2 density, results in a proportional increase in both positive and negative contraction velocities (Fig. 4g).

Relating these trends to the experiments, during the early pulsed cycles, the contraction direction is inward, as there is little accumulation near the boundary. At later cycles there is appreciable boundary accumulation which leads to regions of both inward and outward contraction. Because the network grows less quickly in the pulsed protocol than in the continuous one, it tends to overlap more with the illumination zone (shown schematically as blue circles and black squares in Fig. 4g). At late cycles, though, when the network has grown beyond the illumination zone, the contraction direction is predominantly inward again, as predicted by the model. These complex elastic forces suggest a rich design space in which density gradients and Ca^2+^ profiles can be dynamically sculpted to achieve fine-grained spatiotemporal control over the motion of, for example, particles interacting with the Tcb2 network. We next consider strategies for exerting such control.

### E. Leveraging Tcb2 network growth and contraction for active particle transport

To operationalize the pull and push of Ca^2+^-driven Tcb2 networks for active transport, we perform experiments in which the networks interact with liposomes, lipid particles (Figs. 5a,b) and polystyrene beads (SI Fig. S28 and SI Video Part 3 Section VII); see Methods for details on the liposome preparation. We conducted two experiments demonstrating active particle transport under different illumination protocols. First, we examine how far and how long the system can transport particles under continuous illumination. Upon illuminating a fixed region, the Tcb2 network grows outward and interacts with lipid particles. Some particles are pulled inward, others pushed outward, and some experience sequential push-pull motion (Fig. 5c and SI Section II L). This behavior depends on each particle’s initial radial distance from the illumination zone: particles near the zone become enmeshed in the rapidly forming Tcb2 network and are pulled inward as it contracts, while distant particles are sterically repelled by the expanding network periphery (SI Video Part 3 Section VII). The liposome moves outward by ∼ 30 *µ*m within 10 s. We note that the rapid steric repulsion of the liposome soon after light activation is consistent with a model prediction of a lower-density peripheral network set up throughout the sample due to rapid initial Ca^2+^ diffusion upon first light exposure (SI II M). This peripheral network, at lower Tcb2 concentration than the primary network, does not give an appreciable DIC pattern. Quantifying its density should be accessible to rhodamine-based assays [74], but we leave this for future work.

**FIG. 5.**
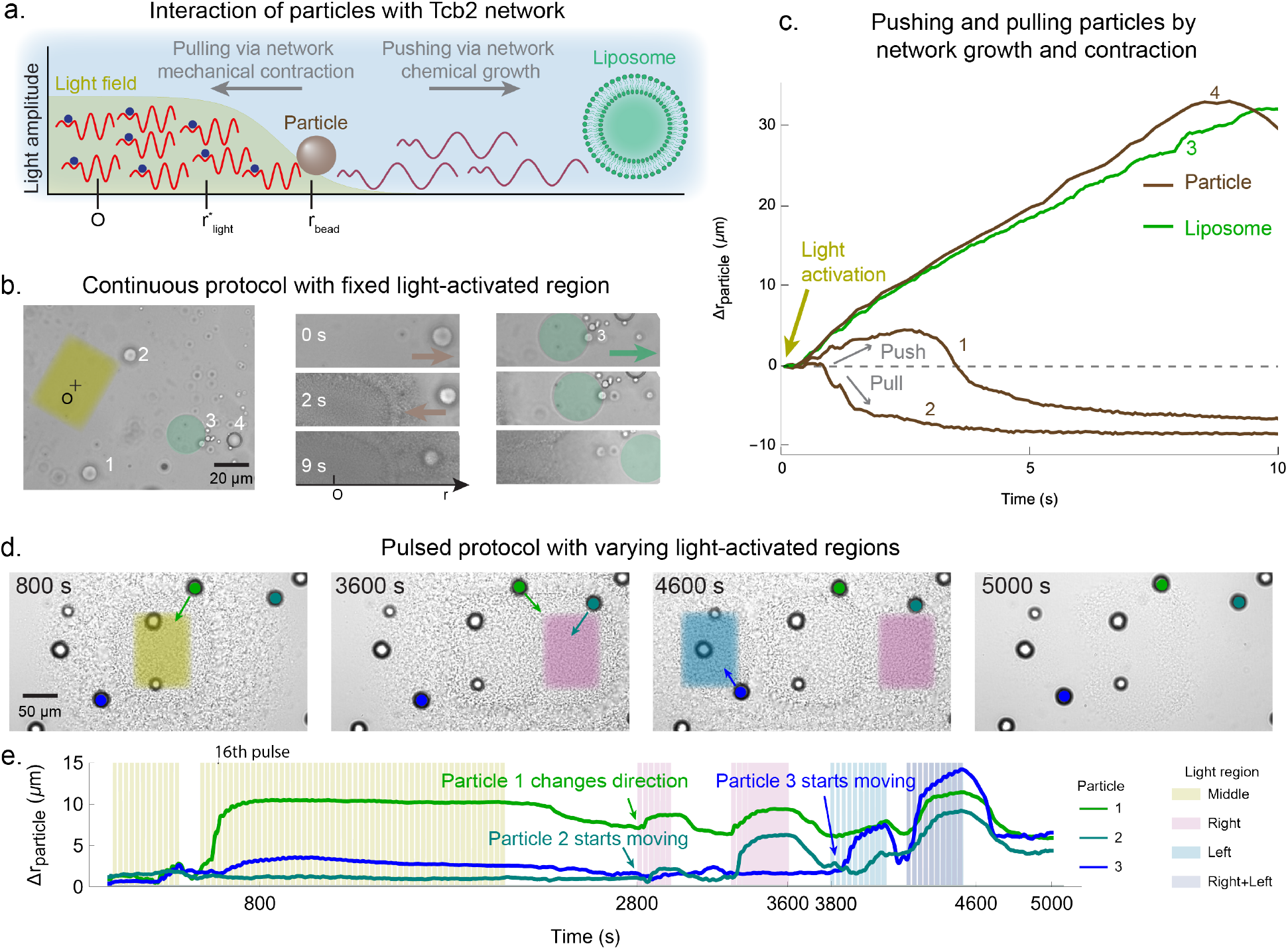
Controlling Tcb2 networks to move external particles. (a) Schematic illustration of light-controlled Tcb2 network interacting with embedded particles and liposomes. (b) Time-series images showing particle and liposome motion during continuous illumination of the yellow rectangular region. Particles 1, 2, and 4 are lipid particles and particle 3 is a liposome. (c) Displacement trajectories of three particles (brown) and one liposome (green) under continuous light activation, achieving outward displacements of up to 30 *µ*m within 10 s. (d) Time-series images demonstrating particle motion as three rectangular regions are sequentially illuminated using a pulsed protocol. (e) Displacement trajectories of three tracked particles from panel (d) over 5000 s. Particle 1 demonstrates two directional changes—moving outward by ∼ 15 *µ*m after the 16^th^ cycle, then reversing rightward at 2800 s and leftward at 3800 s. Colored bands indicate activation periods: yellow (center), purple (right), blue (left), and dark blue (right + left) illumination regions.

Second, we apply a pulsed illumination protocol to explore the system’s transport behavior across multiple illumination patterns. As shown in Fig. 5d and SI Video Part 2 Section VII, the illumination region is sequentially positioned at the center, right, and left of a sample containing embedded particles. Particle movement directly corresponds to the illumination location, demonstrating precise spatial control. Through reprogrammable light patterns, we transport multiple particles achieving displacements of 5–15 *µ*m with two directional changes, sustaining pulsed actuation for approximately 5000 s. For example, in Fig. 5e, Particle 1 reverses direction twice in response to shifts in the illuminated region, moving outward by approximately 15 *µ*m after the 16th cycle and then reversing rightward at 2800 s and leftward at 3800 s. This demonstrates tight coupling between particle motion and activation position. Additionally, at these same time points (2800 s and 3800 s), we selectively activated particles 2 and 3, directing them toward the lower left and upper left respectively. Across the field of view, particle response varied with proximity to the illuminated zone and pulse duration, revealing localized and tunable transport capabilities. By 5000 s, following a 500 s dark period, the system underwent near-complete chemical disassembly while retaining reactivation capacity. We observed small residual particle displacements after illumination cessation, suggesting viscoelastic behavior. Capturing these dynamics in our continuum model will require transitioning in the future from the current damped elastic solid rheology to a viscoelastic fluid description. We show additional particle movement statistics in SI Section II L and SI Fig. S28.

These applications of Tcb2 network growth and contraction to active particle transport suggest a diverse set of options for dynamical control over external objects, although we leave a full exploration of experimental control strategies to future work.

### F. Using reinforcement learning to control Tcb2 contraction

Finally, we test the feasibility *in silico* of closedloop control for positioning particles using light-actuated Tcb2 networks. We consider a task inspired by the experimental application in Fig. 5 of displacing a particle at a specified location in the network (assuming cylindrical symmetry) by a target amount and then maintaining the particle’s position (Fig. 6a). Specifically, we target the time-dependent radial displacement *U*_*r*_(*r*_0_, *t*) evaluated at a fixed radial location *r*_0_ and time *t*. Recent work showed that proportional–integral control can stabilize the spatially averaged velocity in chaotic active nematic channel flow using uniform light intensity [54], illustrating the general feasibility of closed-loop control in active matter at micron scales. Our goal here, however, is particle positioning with multiple inputs and highly nonlinear input-output relationships. To this end we employ reinforcement learning (RL) [78], a data-driven approach suited to high-dimensional non-linear systems. Similar RL-based methods have been applied *in silico* to control dynamics in active nematics, flocking systems, and active crystals [57, 79, 80].

**FIG. 6.**
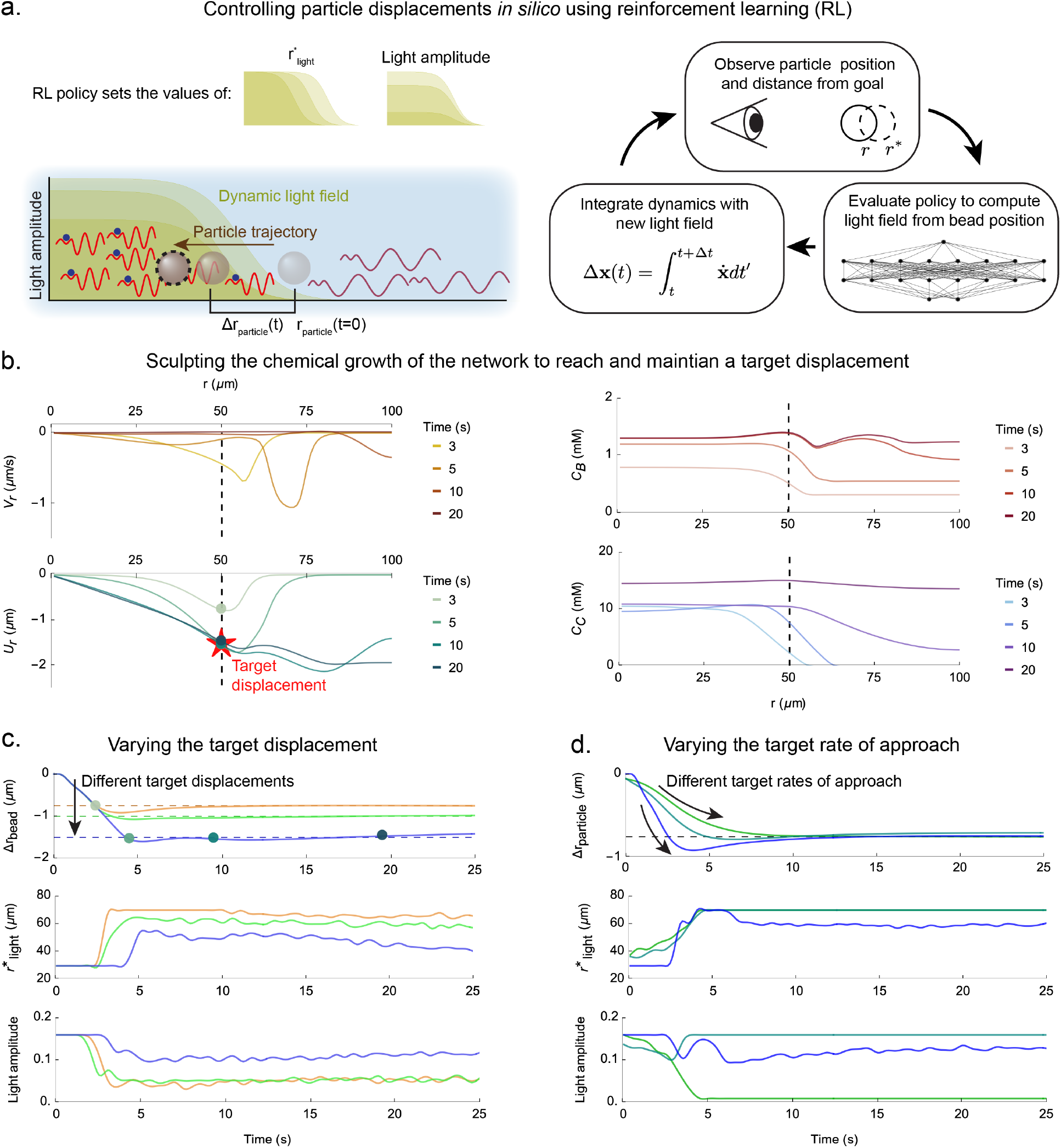
Using reinforcement learning to control displacements. (a) Schematic illustration of the control task considered here and the feedback loop for controlling particle displacements *in silico* using RL. The position and amplitude of a sigmoidal radial light profile is varied dynamically to cause the displacement at certain point to reach a desired value. (b) Plots of the radial velocity *V*_*r*_, radial displacement *U*_*r*_, bound Tcb2 concentration *C*_*B*_, and diffusing Ca^2+^ concentration *C*_*C*_ at different times during an episode. The trained RL policy shines light to sculpt these fields and cause the displacement at *U*_*r*_ (*r*_0_ = 50 *µ*m) to reach its target value of 1. − 5 *µ*m, indicated by the red star. The dashed line throughout indicates the position *r*_0_ = 50 *µ*m. (c) *Top row* : Trajectories of the displacements over time for three different dynamical control tasks targeting different displacements. *Middle row* : The policy choices for the light position corresponding to the trajectories in the top row. *Bottom row* : The policy choices for the light amplitude corresponding to the trajectories in the top row. (d) Same as panel c, but for dynamical tasks targeting different rates of approach.

As discussed more in the Methods and SI I F, we train an actor-critic policy using the deep deterministic policy gradient algorithm [78, 81, 82]. At each time step *t* the RL agent observes the radial displacement *U*_*r*_(*r*_0_, *t*) at a chosen location *r*_0_, and selects as actions the radius and amplitude of a sigmoidal, azimuthally symmetric light profile (Fig. 6a). The reward for training is constructed to minimize the deviation from a specified target *U*_goal_ in a manner mimicking an overdamped spring connecting *U*_*r*_(*r*_0_, *t*) to *U*_goal_. Training is performed *in silico* using our coupled reaction-diffusion-elasticity model. Even though the RL agent observes only a coarse description of the system state, it learns policies that solve different control tasks by sculpting the chemical concentration and velocity fields in time (Fig. 6b), with the effect of driving the displacement at the chosen location to distinct target values (Fig. 6c). The learned strategies typically apply high light amplitude initially to contract the network and move the particle to the target location, after which the light position and amplitude oscillate and drift to compensate for deviations from the target. In SI Fig. S10 we demonstrate that this feedback control is robust to simulated latency of up to at least 3 *s* between system observation and update of the control fields.

To further probe the capabilities of RL-guided control, we considered a more challenging task: driving the displacement to the target value at different rates of approach, mimicking overdamped springs with different stiffnesses. We studied a similar dynamical control task for active nematic defects in Ref. 57. As shown in Fig. 6d, RL successfully solves this problem in simulated Tcb2 networks. The learned policies realize distinct approach rates to the same displacement, demonstrating rate-shaping in addition to set-point control. These results suggest that Ca^2+^-driven Tcb2 networks, despite their highly nonlinear and spatiotemporally history-dependent dynamics, are amenable to control using lowdimensional state and input spaces, and may in the near future be harnessed to exert precise mechanical forces at micrometer and second scales.

## III. DISCUSSION

We have developed an experimental platform and accompanying continuum model to control the Ca^2+^-driven contractile dynamics of Tcb2 protein networks *in vitro*. By reconstituting these networks outside the ciliates in which they naturally occur, we decouple them from the cellular complexity and directly manipulate the chemomechanical processes of proteins homologous to those driving the fastest molecular motions known in biology [37, 62].Using a light-sensitive Ca^2+^ chelator we locally tune the Ca^2+^ concentration in the solution, allowing us to exert fine-grained spatiotemporal control over the network’s growth and contraction. The disparate timescales of these two processes—slow chemical growth and comparatively fast mechanical response—give rise to rich material dynamics. By periodically pulsing light, we can sustain large spatial areas of contractile response for ∼ 150 cycles through a ratchet-like effect. Moreover, the repeated mechanical contractions reflect the slowly evolving concentration profile of the growing network, leading to surprising reversals of contraction direction due to strong density inhomogeneities. The dynamical malleability of this system presents an excellent opportunity for generating on-demand micron-scale forces, which we have explored both *in vitro* by moving lipid particles and *in silico* using reinforcement learning.

This material is different from reconstituted actomyosin or microtubule-kinesin networks in several ways. First, the Tcb2 networks are largely amorphous and isotropic, such that the direction of contraction is not restricted to local filament orientations [19]. Second, our system assembles rapidly (in seconds), compared to cytoskeletally-derived systems (minutes to hours) [73]. The contraction rate of Tcb2 networks is ∼ 0.5 *µ*m/s, which is higher than that of actomyosin networks driven by non-muscle myosin II (∼ 0.05 *µ*m/s) [28, 83, 84] and comparable to the rate of muscle myosin II (∼ 0.25 *µ*m/s) [85–87] and of mixed myosin IIA and myosin IIB networks (∼ 0.4 *µ*m/s) [88]. This system is slower than microtubule-based transport driven by kinesin under light activation, which reaches speeds of ∼ 2.5 *µ*m/s [17, 23, 30, 36], and slower than synthetic moysin XI system [22, 29]. See SI I B for a comparison to specific studies. We note that ultrafast contractile velocities observed in protists such as *Spirostomum ambiguum*, which can reach 0.1 m/s [37, 43], involves a fishnet arrangement of Ca^2+^-binding proteins homologous to Tcb2 with a secondary scaffolding protein [49]. In future work it may be possible to refine the *in vitro* system to account for these biological details to reach similar speeds. Third, this system has a minimal chemical composition, containing just three key components: Ca^2+^, which acts as an external chemical trigger distinct from ATP, competing chelator molecules, and Tcb2 protein. In contrast, in most reconstituted actomyosin networks there are several additional accessory proteins and crowding agents that are typically used to regulate actin polymerization and confine filaments to a surface [89, 90].

The physics of Tcb2 networks presents several opportunities for model development, unification, and control. The system’s mechanics are tightly coupled to its chemistry: diffusive network growth creates strong density inhomogeneities, and Ca^2+^ binding to Tcb2 alters the elastic rest length, necessitating extensions to standard elasticity theory. The reversal of contractile forces due to these density inhomogeneities is a general physical phenomenon that may explain contractility in other systems like cytoskeletal networks [91]. Additionally, the coupling of stress generation to chemistry in our system is reminiscent of earlier works on periodic swelling in synthetic polymer gels due to pattern-forming chemical reaction networks [2, 3], although these swelling motions are much slower. That pulsing the light activation allows for greater force generation through a recharging mechanism may be relevant to understanding the function of, for example, pulsatile dynamics in excitable RhoA-actin systems [92, 93]. Finally, our efforts to control this soft material using reinforcement learning builds on a broader current research thrust of using control theory techniques to design policies that enable precise, ondemand force production in soft active matter systems [50, 51, 53, 54, 57].

This demonstration of light-controlled Tcb2 network dynamics highlights its potential for programmable force generation in synthetic biology. Looking forward, we envision an integrated, closed-loop platform in which live imaging guides feedback control—potentially through reinforcement learning—to precisely manipulate soft active materials *in vitro* and *in vivo* (Fig. 7a). One promising direction involves biotinylated Tcb2: via streptavidin–biotin linkage, the network can be anchored to lipid membranes, enabling controllable membrane deformation and mimicking cellular processes such as cytokinesis or endocytosis [85] (Fig. 7b). The same strategy could also be applied to tether Tcb2 networks to intracellular organelles to facilitate transport or remodeling. Additionally, the AM ester form of DMNP-EDTA can be loaded into live cells by incubation, allowing for localized Ca^2+^ modulation and enabling reconstitution of the system in a cellular context [94]. This offers a feasible approach for studying Tcb2-driven actuation under native-like conditions. The feasibility of light-guided feedback control has also been demonstrated in bulk active nematic flow systems [54], further supporting potential implementations based on Tcb2 networks.

**FIG. 7.**
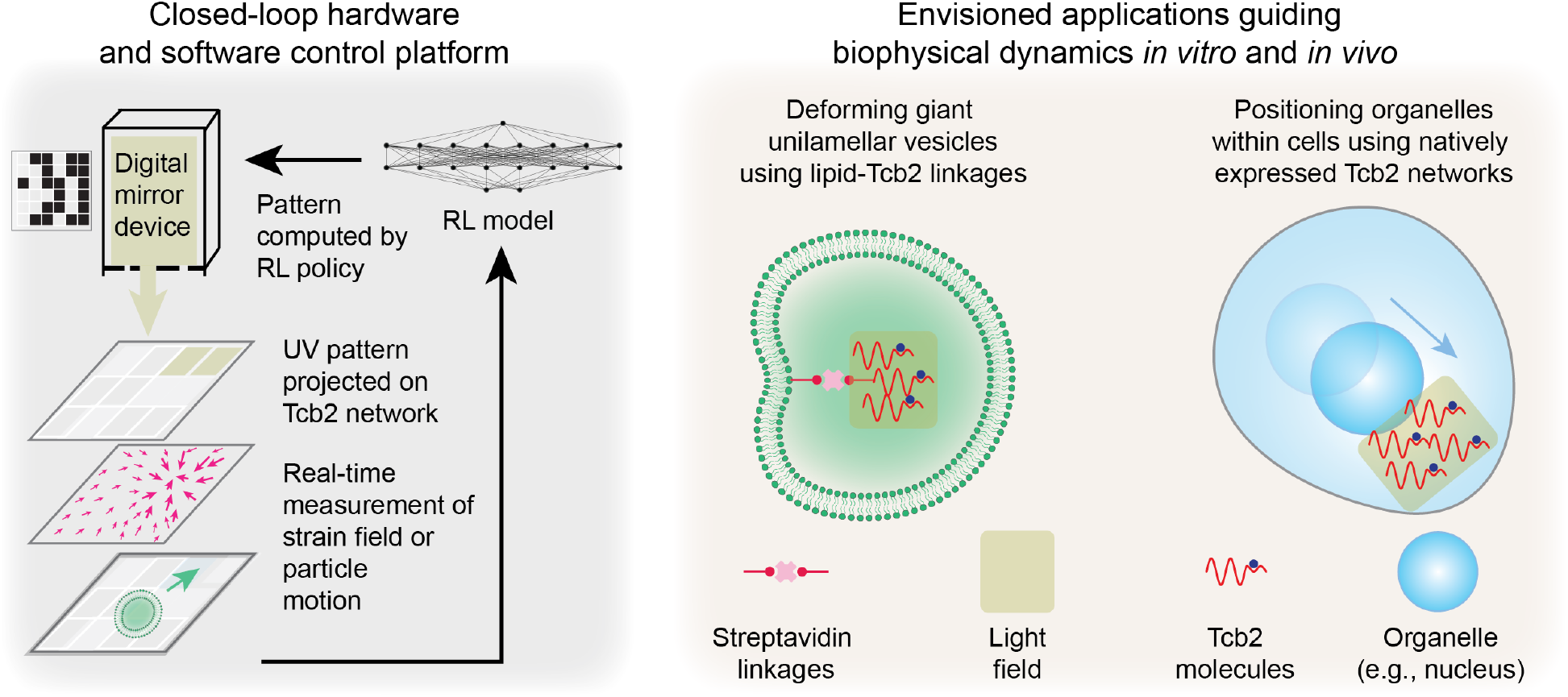
Future applications integrating hardware, software, and biology to control soft active materials. *Left* : Schematic illustration of an integrated experimental platform in which dynamical light patterning is controlled using real-time feedback from imaging of strain fields or particle displacements. A trained RL model or other control architecture maps imaging data to updates in the light field. *Right* : Possible applications of this control platform include deforming lipid vesicles with streptavidin–biotinylated Tcb2 networks, as well as manipulating organelle motion in vivo using light-controlled Tcb2 networks.

In parallel with these applications, the Tcb2 platform may also serve as a tool to uncover fundamental insights into how ultrafast contraction occurs in nature. Reconstituting such behavior in a minimal system could provide mechanistic understanding of contraction in heterotrich protists. Moreover, these supramolecular protein networks could be used to construct synthetic cytoskeletal assemblies with capabilities distinct from those of microtubule- or actin-based systems. For example, they could introduce novel mechanical capabilities, including ultrafast contraction, localized force generation without the need for filamentous tracks, and light-controlled activation that is programmable and decoupled from ATP- or GTP-driven enzymatic pathways. These features open possibilities for intracellular actuation, sensing, therapeutic delivery, and the development of mechanically responsive components in synthetic cells.

## IV. METHODS AND MATERIALS

### A. Tcb2 construction, storage, and purification

We synthesize a codon-optimized full-length version of the Tcb2 protein and clone it into a pJ411 expression plasmid, which we then transform into the BL21 strain of *E. coli* (see SI I C for sequence details). We synthesize GFP- and mCherry-tagged variants of Tcb2 to facilitate experiments shown in Fig. 1a. To prevent aggregation during storage, we store the synthesized proteins in 4 M − urea solution at −80^*°*^C. Prior to experiments, we remove urea by overnight dialysis against a buffer containing 25 mM Tris (pH 7.5) and 1 mM EGTA, using a 3.5–5 kDa MWCO membrane (Spectra-Por Float-A-Lyzer G2, black). After dialysis, we concentrate the protein to approximately 30 mg/mL using a centrifugal filter (Amicon Ultra, UFC500324). We determine the final protein concentration by measuring absorbance at 280 nm with a UV spectrometer.

### B. Light control technique

To ensure complete chelation of Ca^2+^, we prepare a premixed solution of DMNP-EDTA and Ca^2+^ in a 2:1 molar ratio, with final concentrations of 0.54 mM Tcb2, 6.8 mM DMNP-EDTA and 3.4 mM Ca^2+^. We prepare the sample in a “sandwich structure” with a glass slide, a coverslip and a spacer (9 × 9 mM, 25 *µ*L; BioRad, SLF0201) to maintain precise height. We perform imaging using a Nikon Ti2 Eclipse inverted microscope with a 20 × and 40 × objective. To initiate the release of Ca^2+^ from the DMNP-EDTA-Ca^2+^ complex, we project a 365 nm light pattern through a Mightex Polygon 1000-G DMD (SI I D).

### C. Liposome formation

We prepare liposomes using the cDICE method [95] with EggPC (840051P, Avanti Polar Lipids) and an osmotic pressure difference of 10 mOsm/kg between the inner and outer solutions. The inner solution contains 290 mOsm/kg glucose, while the outer solution contains 300 mOsm/kg sucrose in ultrapure MQ water. We centrifuge the resulting aqueous phase containing liposomes at 1000 rpm for 5 minutes to concentrate the liposomes at the bottom of the sample due to the density difference between the inner and outer solutions. We then transfer the 2 *µ*L aliquot from the concentrated liposome layer to a 30 mg/mL Tcb2 solution in a 10% v/v ratio, resulting in a total solution volume of 20 *µ*L. We place the mixture on a glass slide that has been pretreated with a 10 mg/mL poly-L-lysine hydrobromide solution [25, 96]. The lipids particles shown in Fig. 5b are the byproducts of liposomes preparation [97]. We then prepare the sample in the same “sandwich structure” as described in Methods IV B.

### D. PIV analysis and particle preparation

We segment DIC videos using the AI model Segment Anything [98, 99] to track the growth of the Tcb2 network. We employ the prompt-based initialization model to enhance segmentation speed and accuracy for the Tcb2 network. For PIV analysis, we adapt the PIV MATLAB code to track Tcb2 filaments [100], with images pre-processed using a Gaussian filter (standard deviation: 12 pixels), tracked using a nearest-neighbor algorithm within a search window grid of 48 pixels and post-validated using a correlation coefficient threshold of 0.6 to 0.8. We perform tracking and trajectory analysis of liposomes, lipid particles, and fluorescent particles using the AI tool Co-Tracker, with a manually set initial prompt-point and a grid size of 50 pixels [101]. We use 10-22 *µ*m diameter Fluorescent Red Polyethylene Microspheres (cospheric UVPMS-BR-0.995) for SI Video Part 2 Section V.

### E. Continuum model

Our chemomechanical model uses reaction-diffusion dynamics, for the evolution of the chemical concentration fields coupled to an overdamped elastic solid with viscous drag for the evolution of the displacement fields, whose density and rest strain depend on local chemical concentrations (see SI I E for details). To determine the motion of the system we assume an instantaneous force balance between viscous drag and the local the elastic force, which comprises several non-standard terms due to inhomogeneity of the Tcb2 density. The external light field is a treated as a non-autonomous source function in the model whose magnitude spatiotemporally sets the − rate of Ca^2+^ release by DMNP-EDTA. We distinguish between diffusing and bound Tcb2, as well as between Ca^2+^-bound and unbound Tcb2. As a first approximation, we assume a simple model for nucleation and growth of the Tcb2 network, in which available Tcb2 binding sites depend non-monotonically on the local concentration of bound Tcb2 to capture the competing effects of bound Tcb2 concentration locally creating new binding sites while also contributing to steric interference. In SI Fig. S8 we explore the effect on varying the saturation concentration of Tcb2 networks. Ca^2+^-induced growth of the bound network is modeled by taking the unbinding rate of Ca^2+^-bound Tcb2 from the network to be significantly less than that of Tcb2 not bound to Ca^2+^. We incorporate the degradation of DMNP-EDTA upon photolysis as a fractional contribution to the viable DMNP-EDTA concentration following Ca^2+^ release. For simplicity we neglect possible advection of diffusing particles by motion of the network; in SI Fig. S9 we show that including advection makes little quantitative difference for this low (0.07 − 2) Péclet number system. To accelerate calculations for azimuthally symmetric light patterns we solve the equations of motion in cylindrical coordinates and neglect the angular dependence. We constrain parameterization of the model by the known reaction rates and otherwise determine parameters through exploration to produce dynamics which match the experimental observables. We give parameter values in SI I E.

### F. Simulation methods

To integrate the dynamical equations we developed custom Julia code which implements Heun’s method, a finite difference predictor-corrector scheme. We impose no-flux Neumann boundary conditions for the chemical species, preventing them from diffusing away from the simulation volume. We impose Dirichlet conditions **u** = **0** for the displacement field at the boundary. We choose sufficiently fine spatial and temporal discretization to ensure numerical stability and convergence.

### G. Reinforcement learning

We define the RL learning task in terms of an imposed dynamical law on the displacement *U*_*r*_(*r*_0_, *t*) at a fixed radial coordinate *r*_0_ of the material. We use RL to learn a mapping (called a policy) from *U*_*r*_(*r*_0_, *t*) to the radius of the azimuthally-symmetric sigmoidal light profile and its amplitude. This mapping should mimic the dynamics of an overdamped spring connecting the displacement at position *r*_0_ to the target displacement *U*_goal_. We vary *U*_goal_ and the overdamped spring constant to test the ability to learn different dynamical tasks, as in our previous work on active nematics [57]. We train the system using the deep deterministic policy gradient algorithm (DDPG), which is a variant of the actor-critic method [78, 81, 82]. For this we combine our custom numerical integrator of the Tcb2 dynamics with the DDPG implementation provided by the Julia package ReinforcementLearning.jl [102]. Additional details of the RL problem formulation and training algorithm are given in SI I F.

## ACKNOWLEDGMENTS

We wish to thank Douglas Chalker for providing the Tcb2-GFP tagged *Tetrahymena* strain, Heidi Sleister and students for creating the GFP/mCherry versions of Tcb2 and Mary Elting, Fred Chang, Jane Maienschein, and Suri Vaikuntanathan for helpful discussions. We also ac knowledge the use of ChatGPT to refine the language in the supplementary information. A.R.D. acknowledges support from National Science Foundation (NSF) award 2313725 and the University of Chicago Materials Research Science and Engineering Center funded by NSF award 2011854. S.C. acknowledges support from NSF award 2313723 and the David and Lucille Packard Fellowship for Science and Engineering. J.H. acknowledges support from NSF award 2313727. S.B. acknowledges support from NSF award MCB-2313724, National Institutes of Health (NIH) award R35GM142588 and Schmidt Sciences, LLC. The authors acknowledge the University of Chicago’s Research Computing Center for computing resources.

## Supplemental Material

## I. SUPPLEMENTARY METHODS

### A. Biological background on Tcb2 and its homologs in ciliates

**FIG. S1.**
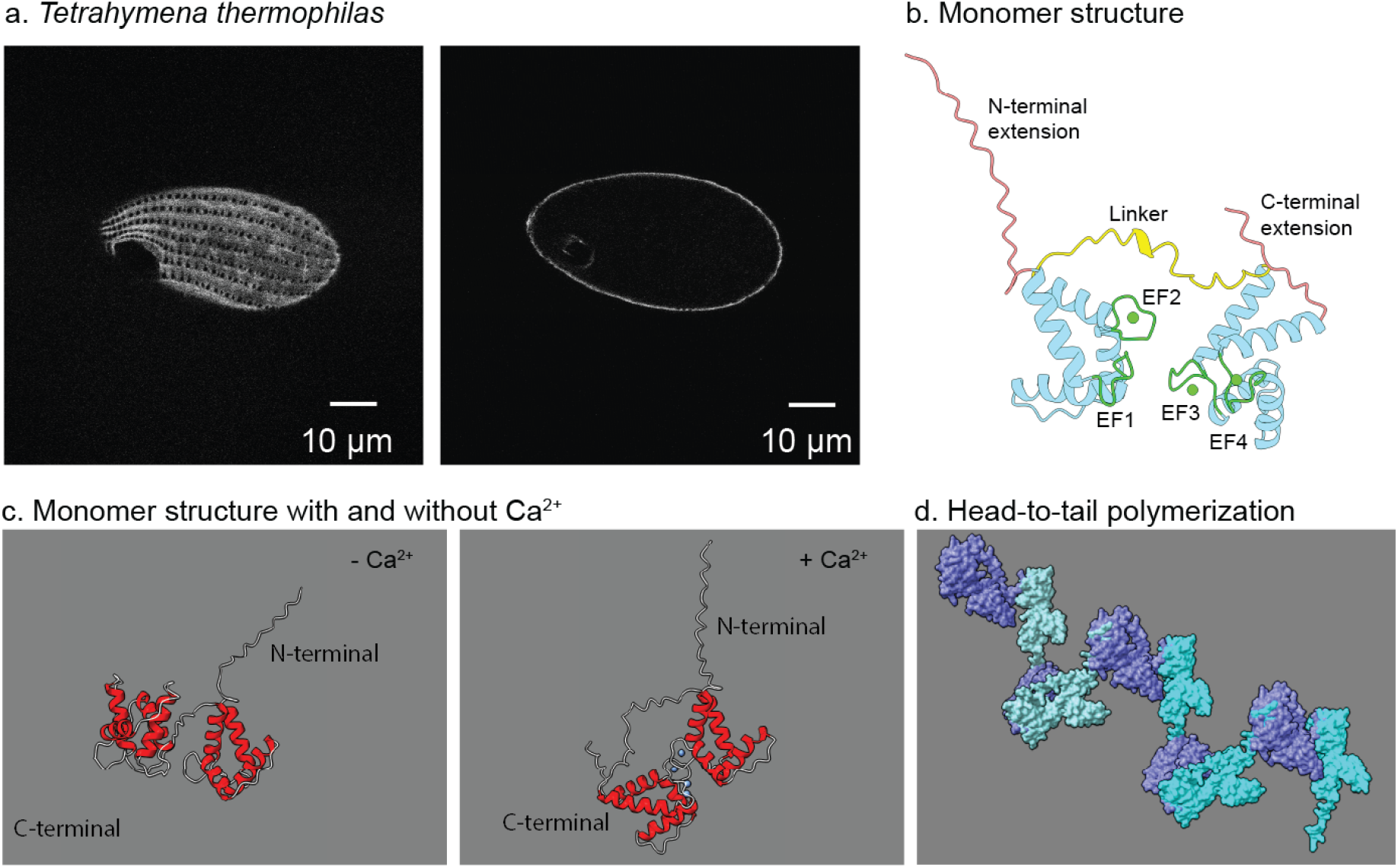
*Tetrahymena* and Tcb2 structure. (a) Confocal laser scanning immunofluorescence microscopy images of *Tetrahymena thermophila* cell expressing a GFP-tagged version of Tcb2. The left panel shows the top plane, while the right panel captures the midway plane, highlighting the specific localization of Tcb2-GFP to the submembranous epiplasmic layer. (b) AlphaFold3 [1] prediction of Tcb2 monomer showing EF-hand domains (EF1 to EF4), disordered N-terminal extension, flexible linker and C-terminal extension. (c) AlphaFold3 prediction of Tcb2 monomer structure in the absence and presence of Ca^2+^, illustrating structural changes upon Ca^2+^ binding. (d) AlphaFold3 prediction of Tcb2 polymerization with a head-to-tail connection.

*Tetrahymena* Tcb2, first designated as TCBP-25, is a 25-kDa Ca^2+^-binding protein [2, 3]. It has been found localize to the cortical cytoskeleton (epiplasm) of *Tetrahymena thermophila* cells [2, 4], which we also observe through flurouescence imaging *in vivo* in Fig. S1a. Tcb2 has been implicated in Ca^2+^-mediated gametic pronuclei exchanged during mating and conjugation [3, 5]. It was also identified as a key component of a contractile gel formed from an alkaline low ionic strength extract of the *Tetrahymena* membrane skeleton, along with several other proteins including a high molecular weight protein Epc1 [6, 7]. We expressed a synthetic gene encoding Tcb2 in bacteria and discovered that the purified protein had Ca^2+^-triggered contractile properties independent of its association with Epc1. Additionally, we found that a Ca^2+^-responsive network can be reconstituted solely with Tcb2, leading us to focus specifically on the role of this protein.

Tcb2 is reported to have four potential Ca^2+^-binding sites, along with a longer flexible linker between its N- and C-terminal EF-hand domains [3, 8]. Upon Ca^2+^ binding, the C-terminal domain of Tcb2 undergoes a substantial conformational change, as evidenced by NMR data [7]. Additionally, our predictions using Alphafold3 suggest that the C-terminal flips after binding to Ca^2+^ (Figs. S1b,c). Unlike calmodulins, Tcb2 forms filaments. In response to Ca^2+^ ions, Tcb2 assembles into a contractile network, as shown in SI video Part 1, Section I. Structurally, in addition to the longer linker already mentioned, Tcb2 also features an extended N-terminal structure and a shorter C-terminal extension (Fig. S1b). We note that these features are absent in calmodulin, another Ca^2+^ binding protein which is not thought to play a major structural or mechanical role.

Gel filtration chromatography suggests that Tcb2 forms polymers even in the absence of Ca^2+^. Mass photometry supports this, further revealing that in the presence of Ca^2+^ there is a change in mass distribution indicating the formation of larger polymers. However, more research is required to determine the exact molecular role of Ca^2+^ in these processes. Here we propose two hypotheses on how Ca^2+^ reacts with Tcb2. First, Ca^2+^ binding may cause Tcb2 to assemble into longer filaments through head-to-tail assembly (Fig. S1d). Alternatively, Ca^2+^ may facilitate the aggregation of these filaments, leading to branching. However, it remains unclear whether Ca^2+^ binding drives the elongation of these polymers by adding more subunits or whether it promotes the formation of branching networks, as observed in fluorescence images in the presence of Ca^2+^ (Fig. 1 c,d). It is also possible that both mechanisms are involved.

**TABLE S1.**
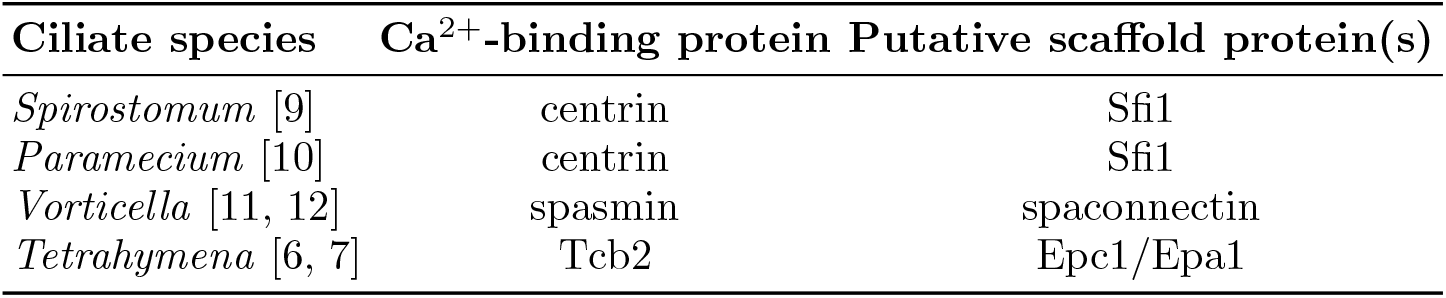
Comparison of Ca^2+^-triggered contractile proteins in ciliates.

Ciliates can utilize ATP-independent Ca^2+^ systems for motility, such as in the contractile stalk of *Vorticella* [11, 12] and the ultra-fast contraction of *Spirostomum* [17]. In these systems, contractility is hypothesized to result from a Ca^2+^-dependent interaction between a Ca^2+^-binding protein and an alpha-helical scaffold protein, commonly referred to as the “beads-on-a-string” model [18]. Centrins, EF-hand proteins, and associated alpha-helical scaffold proteins that contain tandem repeats of motifs are identified in Ca^2+^-triggered ATP-independent contractile systems in *Spirostomum* [9].

Several centrin-like proteins have been identified in ciliates, as shown in Table S1 and Fig. S2. In *Paramecium*, centrin, together with the scaffold protein Sfi1, forms filaments within the infraciliary lattice that are hypothesized to coil or kink in response to Ca^2+^, leading to contraction [18]. Similarly, the contractile stalk of *Vorticella* contains a related protein, spasmin, which requires an alpha helical scaffold protein called spaconnectin to form Ca^2+^-sensitive contractile filaments [11, 12]. The centrin, spasmin, and Tcb2 proteins illustrate the diversity of Ca^2+^-triggered contractility that has evolved in ciliates. In particular, unlike centrin and spasmin, subsequent bacterial expression of a synthetic gene encoding Tcb2 revealed that Tcb2 exhibits Ca^2+^-triggered contractile properties independently of the Epc1 association.

**FIG. S2.**
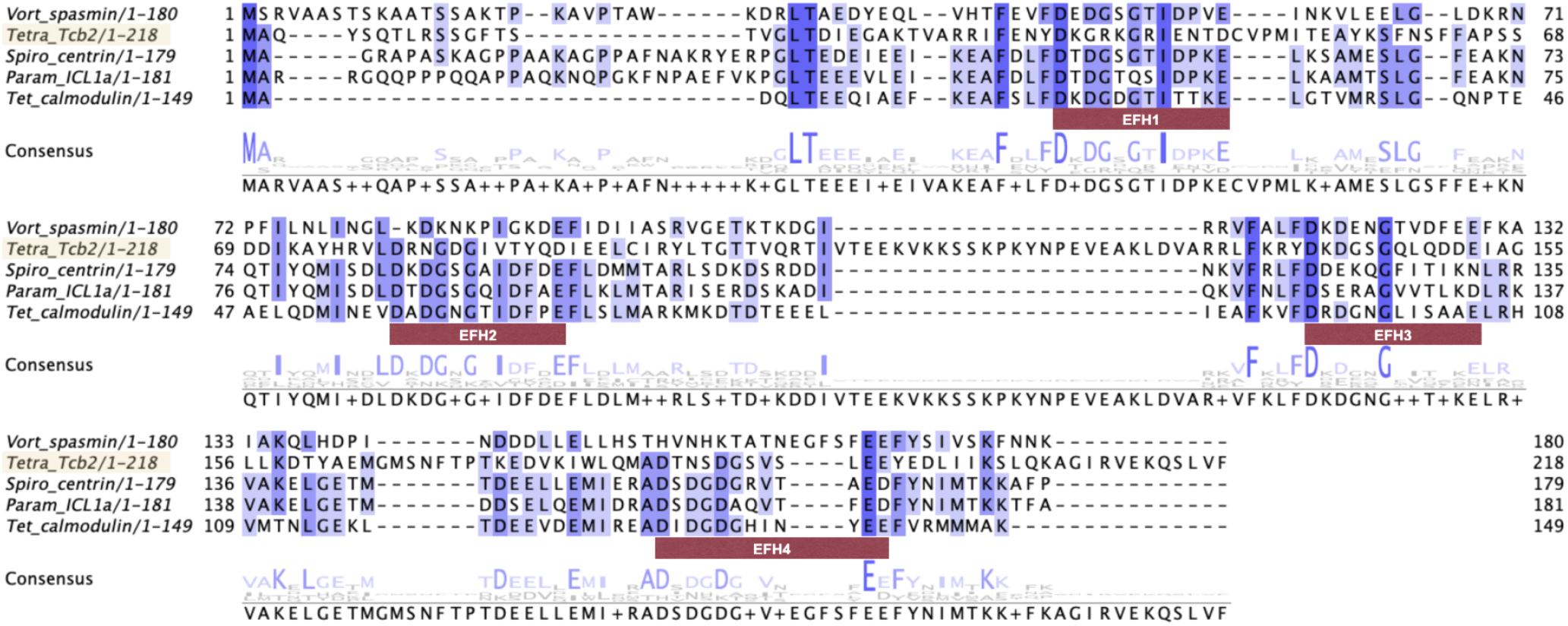
Homolog sequence comparison of Tcb2 in*Tetrahymena*. Protein sequences observed or predicted to be involved in calcium-dependent contractile processes in ciliates and a ciliate calmodulin as a reference were retrieved from the National Center for Biotechnology Information. *Vorticella* spasmin: GenBank: AAD00995.1 [13]; *Tetrahymena* Tcb2: UniProtKB/Swiss-Prot: P09226.2 [14]; *Paramecium* ICL1a: UniProtKB/Swiss-Prot: Q27177.2 [15]; *Tetrahymena* calmodulin: PIR: S28954 [16]. The sequence of *Spirostomum* centrin is taken from our unpublished data. These sequences were aligned in Jalview using default settings for the MAFFT multiple sequence alignment server. Predicted EF-hand calcium-binding sites in Tetrahymena calmodulin, as predicted by PROSITE, are indicated.

### B. Comparison with other biomimetic contractile systems

In Table S2, we compare two categories of cytoskeletal reconstitution systems: actomyosin systems (with both natural myosin motors and synthetic light-activated motors), and microtubule-kinesin systems (with both natural kinesin and synthetic light-activated motors). We note that in each case the reported velocity may correspond to slightly different measurements (such as filament motion, motor motion, or contractility rate), and we report these values as rough estimates of the typical system speed.

A key distinguish feature among these system is whether they can be dynamically regulated. Some systems employ light-activated motors [19, 20] or light-activated inhibitors such as blebbistatin [21–25] that enable external control over contraction, allowing researchers to externally control motor activity. In contrast, many reconstituted systems lack such regulatory mechanisms and exhibit irreversible dynamics: once activated, they proceed uni-directionally though continuous contraction [26, 27], directional migration [22, 28] or turbulent-like flows [29] until their ATP fuel is exhausted.

### C. Tcb2 construction and characterization

We cloned a synthetic Tcb2 gene (shown in I C 1), optimized for expression in *E. coli*, into plasmid pJ411 (high copy number, kanamycin resistance plasmid, with expression controlled by a T7 promoter), and transformed it into BL21 strain *E. coli* cells. We grew cells in lysogeny broth (LB) or super broth (SPB) media containing kanamycin, and we induced expression by adding IPTG to 1 mM. We grew cells overnight at 18^*°*^C and harvested by centrifugation. We lysed cells using B-PER detergent solution and centrifuged the lysate centrifuged at 15,000 g for 15 minutes. We extracted Tcb2 protein from the post-lysate pellet using 4M urea, 0.25 mM EGTA, 25 mM Tris-HCl, pH 7.5 buffer. We loaded the clarified extract onto a 25 mL Q-Sepharose anion exchange column and eluted with a NaCl step gradient in the presence of 4M urea. We concentrated and stored the Tcb2-containing fractions in urea-containing buffer at -80^*°*^C.

**TABLE S2.** A comparison of flow rates, motor proteins, and dynamical regulation mechanisms across different cytoskeletal systems.

| System | Maximal flow velocity | Motor | Dynamically regulated | Refs. |
| --- | --- | --- | --- | --- |
| Actomyosin | 0.06 $\mu\text{m/s}$ | 24 nM non-muscle myosin II | Yes (blebbistatin photocontrol) | [21] |
| Actomyosin | Continuous: 0.42 $\mu\text{m/s}$, Periodic: 3.33 $\mu\text{m/s}$ | 2.0 $\mu\text{M}$ myosin IIA, 0.2 $\mu\text{M}$ myosin IIB | Periodic waves | [26] |
| Actomyosin | 0.3 $\mu\text{m/s}$ | Skeletal muscle myosin II | No (irreversible) | [30] |
| Actomyosin | 0.25 $\mu\text{m/s}$ | Skeletal muscle myosin II | Yes (blebbistatin photocontrol) | [22] |
| Actomyosin | 0.17 $\mu\text{m/s}$ | Skeletal actin from chicken skeletal muscle | No (steady contraction) | [27] |
| Actomyosin | 1 $\mu\text{m/s}$ | Skeletal muscle myosin, or heavy meromyosin from rabbit muscle | Yes (blebbistatin photocontrol) | [23] |
| Actomyosin | 4.8 $\mu\text{m/s}$ | Non-processive heavy meromyosin | No (unidirectional motion) | [28] |
| Actomyosin | 4 $\mu\text{m/s}$ | Actin-intact cytoplasmic extracts from *Xenopus* eggs | Periodic oscillations | [31] |
| Actomyosin | 0.9 $\mu\text{m/s}$ (speed of transported lipid droplet) | Actin-intact cytoplasmic extracts from *Xenopus* eggs | No (irreversible, directional locomotion) | [32] |
| Actomyosin-rho | 0.2 $\mu\text{m/s}$ (speed of wave) | *Xenopus* embryos | Oscillatory waves | [33] |
| Actomyosin | 2.5 $\mu\text{m/s}$ (filament motion) | Synthetic (light-activated) myosin XI | Yes (optogenetic control) | [19] |
| Actomyosin | 10 $\mu\text{m/s}$ (filament motion) | Synthetic (light-activated) myosin XI | Yes (optogenetic control) | [20] |
| Microtubule-kinesin | 2.2 $\mu\text{m/s}$ | Kinesin | No (turbulent-like flows) | [29] |
| Microtubule-kinesin | 1.8 $\mu\text{m/s}$ | Kinesin | No (extensile dynamics) | [34] |
| Microtubule-kinesin | 2.5 $\mu\text{m/s}$ | Synthetic (light-activated) kinesin | Yes (optogenetic control) | [24] |
| Microtubule-kinesin | 1 $\mu\text{m/s}$ | Synthetic (light-activated) kinesin | Yes (optogenetic control) | [25] |

#### 1. Protein sequence

We used the sequence of TcBP-25 in the National Center Biotechnology Information (NCBI) database, UniProtKB/Swiss-Prot: P09226.2 [14]. The sequence is (see also Fig. S2):

1 MAQYSQTLRSSGFTSTVGLTDIEGAKTVARRIFENYDKGRKGRIENTDCVPMITEAYKSFNSFFAPSSDD 69

70 IKAYHRVLDRNGDGIVTYQDIEELCIRYLTGTTVQRTIVTEEKVKKSSKPKYNPEVEAKLDVARRLFKRY 139

140 DKDGSGQLQDDEIAGLLKDTYAEMGMSNFTPTKEDVKIWLQMADTNSDGSVSLEEYEDLIIKSLQKAGIR 209

210 VEKQSLVF 218

#### 2. Ca^2+^ concentration for assembly

We conducted two experiments to determine the critical Ca^2+^ concentration for Tcb2 assembly. We first perfoemd a pelleting assay test. We mixed freshly diluted Tcb2 with a gradient of Ca^2+^ concentrations. At high centrifugal speed, the Tcb2 protein network pelleted (bottom sediment), while monomeric Tcb2 remained in the supernatant (top layer solution). We analyzed the concentration of Tcb2 in both the pellet and the supernatant using SDS-PAGE [35]. We observed a critical shift, where the bulk of Tcb2 moved from the supernatant to the pellet. The critical concentration for assembly is roughly 0.47 *µ*M Ca^2+^. In the second experiment, we monitored the progression of the Tcb2-Ca^2+^ reaction using a UV spectrometer at 280 nm, with a gradient of Ca^2+^ concentrations. We selected this wavelength based on prior determinations of Tcb2 concentration. Higher absorbance values indicate a denser Tcb2 network. We observed rapid and stable assembly at a critical concentration of approximately 4 *µ*M Ca^2+^.

### D. DMD and optical setup

We customized our microscope by integrating a multi-port illuminator to enable 365 nm patterned illumination, while retaining the original light source for fluorescence. The components we use are listed in Table S3. The excitation filter was moved from the microscope’s filter cube to the Multi-Port Illuminator. We use two dichroic mirrors (see the light path diagram in Fig. S3): a 450 nm shortpass filter reflects mCherry/green excitation light and transmits UV, allowing a combination of fluorescence and patterned illumination from the polygon projector, and a Multi-band Dichroic Mirror II transmits mCherry/green excitation light and 365 nm light toward the sample, while reflecting mCherry/green emission light from the sample to the camera. With this setup, the DMD pattern projector achieves sub-micron resolution with various patterns, as shown in Fig. S4. The smallest observed assembly size is approximately 20 *µ*m × 20 *µ*m.

To control Ca^2+^ release using the DMD, we use the photolyzable chelator DMNP-EDTA, whose chemical structure is shown in Fig. S5.

**FIG. S3.**
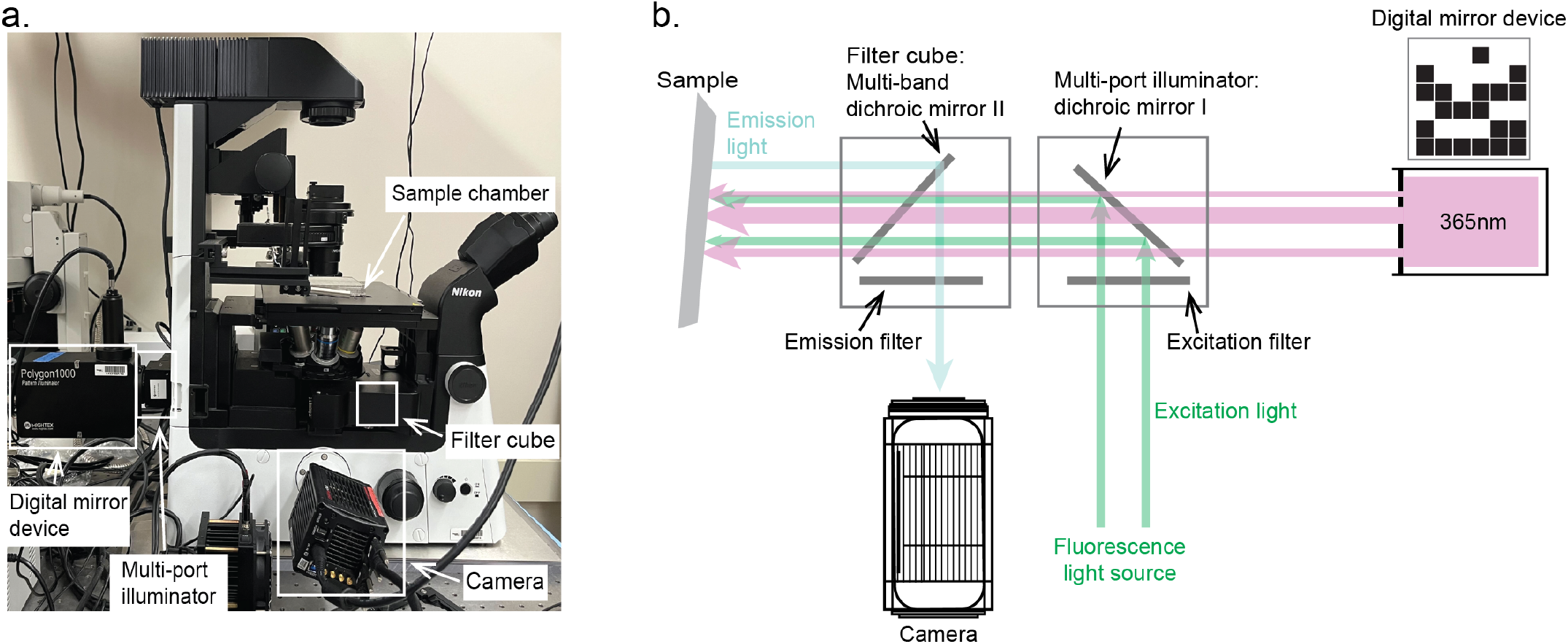
Experimental setup for patterned light illumination. (a) Microscopy system with main components highlighted. Illustration of the optical path with two main modules: the multi-port illuminator and filter cube alignment corresponding to panel a.

Throughout this paper, we performed comparative experiments on Tcb2 network assembly and contraction under a range of conditions. We performed these experiments with three to four replicates for each condition, analyzing distinct regions positioned at least 200-300 *µ*m apart on the sample coverslip to ensure spatial separation for each sample. This ensured that the regions were not interconnected and minimized any potential effects of protein aggregates due to diffusing Ca^2+^.

**TABLE S3.** Optical components used in the experimental setup.

| Optical Component | Model |
| --- | --- |
| Dichroic mirror I | Mightex FLTR-DCH-450S, Dichroic, 450 nm shortpass |
| Multi-band dichroic mirror II | Chroma 343448, Reflection: 365 nm LED and 59022x, Transmission: 59022m |
| Emission filter | Chroma 368163 (Transmission: 500-550 nm, 600-650 nm) |
| Excitation filter | Chroma 369288 (Transmission: 450-500 nm, 550-600 nm) |

### E. Continuum model

#### 1. Model motivation

We develop a physical model for the light-activated Tcb2 networks based on principles of reaction-diffusion chemistry and elasticity theory, with the goal of providing mechanistic explanations for several phenomena observed in our experiments. These include the localization of contractile force to the periphery of a continuously illuminated sample (main text Fig. 2d), a systematic heterogeneity of Tcb2 density along the radial direction (main text Fig. 2a and Fig. S26 below), the ability to repeatably recharge the network contraction upon pulsing the light rather than shining it continuously (main text Fig. 3d), and the surprising reversal of contraction direction from radially inward to outward at intermediate pulsation cycles (main text Fig. 4a). We find that a model based on first-principles considerations of the chemical reaction-diffusion dynamics and elasticity theory with a simple scaling between stiffness and network density suffice to recapitulate each of these phenomena. The model is not quantitatively accurate, however, largely due to a simplified treatment of network nucleation and growth, unaccounted-for influence of buffers in the solution, lack of treatment of viscoelastic effects and a general lack of experimentally constrained model parameters. We therefore view this model as a first iteration to be refined in subsequent work, but emphasize that it currently reproduces and explains key experimental phenomenology as discussed in the main text.

**FIG. S4.**
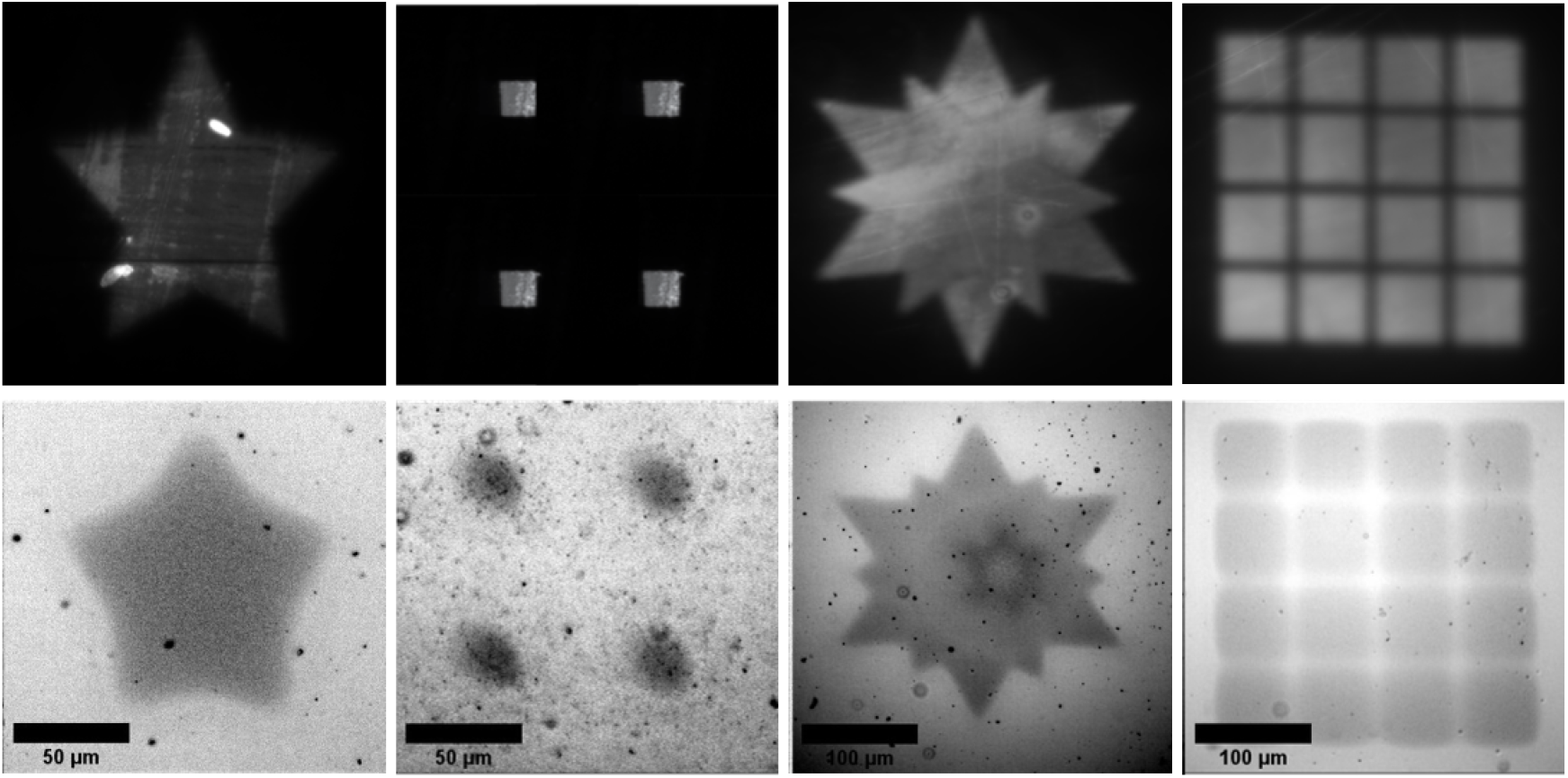
Patterned illumination. *Top row* : A digital mirror device (DMD) projects various patterns onto a highlighter-coated mirror slide to visualize the UV pattern. We note that the uneven distribution of the light pattern is due to the highlighter strokes on the slides and visible scratches on the mirror, not the projector. *Bottom row* : The Tcb2 with DMNP-EDTA-Ca^2+^ system responds to the projected light patterns by forming protein assemblies with sharp boundaries, imaged using DIC.

**FIG. S5.**
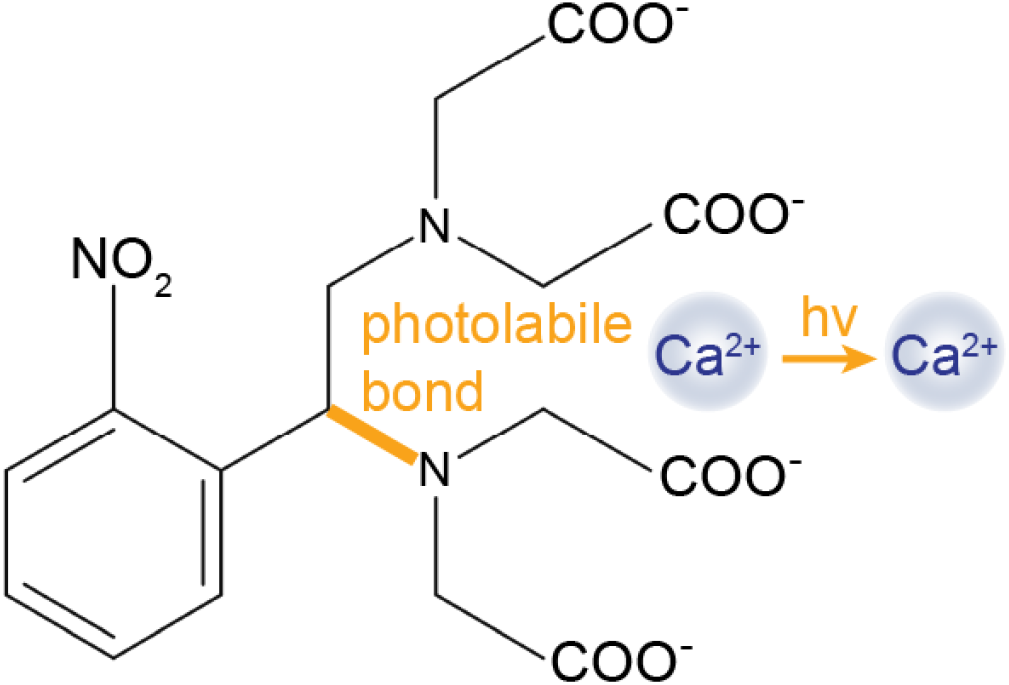
Chemical structure of DMNP-EDTA. DMNP-EDTA has a photolabile bond which releases Ca^2+^ in response to light.

#### 2. Equations of motion

Here we outline the model equations of motion for a mixture of Ca^2+^, Tcb2 molecules, and DMNP-EDTA chelators. We assume that diffusing Tcb2 molecules can bind to a network of bound Tcb2 filaments, and the bound filaments’ mechanical rest lengths change upon Ca^2+^ activation. Ca^2+^ ions are sequestered by the available DMNP-EDTA chelators, and their release from these chelators is modulated by an externally applied light field. We track spatiotemporal concentrations of the following chemical species:

- Diffusing Tcb2 molecules in their inactivated (not Ca^2+^-bound) state, denoted *DI*.
- Diffusing Tcb2 molecules in their activated (Ca^2+^-bound) state, denoted *DA*.
- Bound Tcb2 molecules in their inactivated state, denoted *BI*.
- Bound Tcb2 molecules in their activated state, denoted *BA*.
- Diffusing Ca^2+^ ions, denoted *C*.
- Diffusing DMNP-EDTA molecules which do not contain a Ca^2+^ ion, denoted *D*^∗^.
- Diffusing DMNP-EDTA molecules which contain a Ca^2+^ ion, denoted *D*.

These species participate in the following reversible reactions:

- (In)activation of diffusing Tcb2 by Ca^2+^ (un)binding

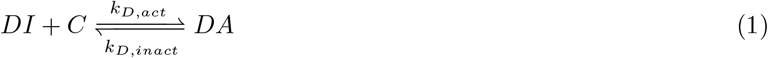
- (In)activation of bound Tcb2 by Ca^2+^ (un)binding

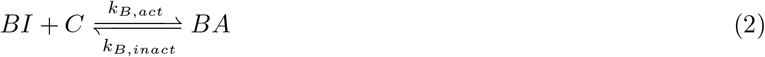
- Trapping (release) of Ca^2+^ by DMNP

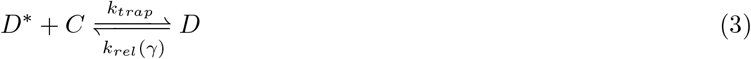

Here, *γ*(**r**, *t*) is a non-autonomous function representing a field of light that speeds up the released rate of DMNP-EDTA via the dependence *k*_*rel*_(*γ*).
- (Un)binding of *DA* to an open binding site *O* in the network

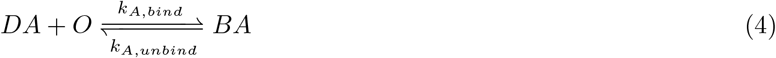
- (Un)binding of *DI* to an open binding site *O* in the network

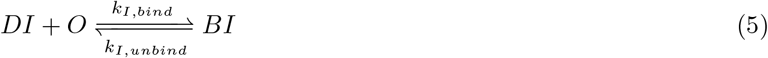

The precise nature of how Tcb2 binds to other Tcb2 monomers to form the bound network is unknown, so we use a simple phenomenological model for how the concentration of available binding sites *O*(*C*_*B*_) depends on the concentration of bound Tcb2 *C*_*B*_ ≡ *C*_*BA*_ + *C*_*BI*_. We take this function to be non-monotonic because of two competing effects: binding of Tcb2 to the network opens up new binding sites, and when there is too much locally bound Tcb2 then binding sites become sterically blocked. This steric hindrance is necessary to prevent a run-away positive feedback of binding, and it causes the concentration of bound Tcb2 to saturate at some level *C*_*sat*_. We take the simple non-monotonic form

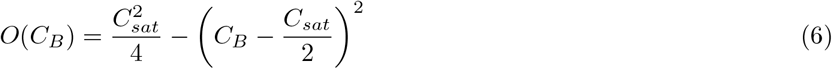

which is a downward facing parabola that intercepts *O* = 0 at *C*_*B*_ = 0 and *C*_*B*_ = *C*_*sat*_ and has a maximum at

The concentrations of these seven species are dynamical variables of the model, being functions of position **r** and time *t*. The dynamics of the diffusing molecules (*DI, DA, C, D, D*^∗^) have spatial dependence due to their concentration gradients.

To treat the linear elastic dynamics of the bound Tcb2 network, we introduce an additional vector field **U**(**r**, *t*) and its derivative **V** = ∂_*t*_**U** which represent the displacement and velocity vectors of the network. The mechanical degree of freedom **U** is coupled to the chemical concentration variables in that the constitutive equation for the elastic deformation depends on the local chemical variables *C*_*B*_ and thr fraction of Ca^2+^-bound Tcb2 in the network *p*_*A*_ ≡ *C*_*BA*_*/C*_*B*_. Introducing the elastic stress tensor *σ*_*ij*_ (where the indices run over Cartesian directions *x* and *y*), the overdamped elastic dynamics have the force balance

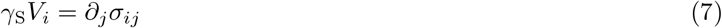

where *γ*_S_ is the Stokes drag coefficient. Introducing the symmetrized elastic strain tensor *ϵ*_*ij*_ = (1*/*2) (∂_*i*_*U*_*j*_ + ∂_*j*_*U*_*i*_), we use the isotropic elastic constitutive equation

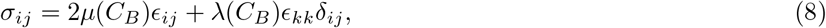

where *µ*(*C*_*B*_) and *λ*(*C*_*B*_) are the first and second Lamé parameters of the Tcb2 network. Because the network should be stiffer where it is denser [36], we assume that these elastic moduli increase linearly with local concentration of bound Tcb2:

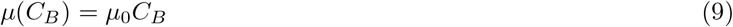

where *µ*_0_ captures the slope of the dependence on *C*_*B*_. We use a similar relation for *λ*(*C*_*B*_).

To account for the fact that Ca^2+^ activation of bound Tcb2 induces a change in the rest length of the Tcb2 molecules, we introduce an autogeneous (rest) strain *g*_*ij*_(*p*_*A*_) which depends on the local proportion of activated Tcb2 molecules *p*_*A*_ as in our previous work [37]. We assume that this strain is isotropic, that is

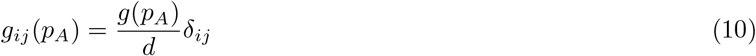

where *g*(*p*_*A*_) = *g*_*kk*_(*p*_*A*_) is the the trace of *g*_*ij*_ in *d* dimensions. Using this, we update Equation 8 to

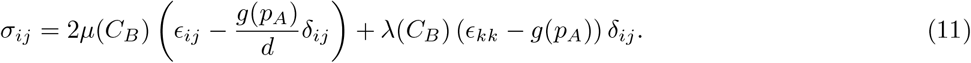

We take *g*(*p*_*A*_) to be a linear function between 0 (when *p*_*A*_ = 0, no autogeneous strain) and −(1−*g*_min_) (when *p*_*A*_ = 1):

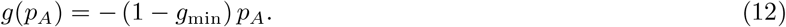

Putting everything together into a closed set of dynamical equations for the variables *C*_*DA*_, *C*_*DI*_, *C*_*C*_, *C*_*D*_, *C*_*D*∗_, *C*_*BA*_, *C*_*BI*_, and **u**, we have

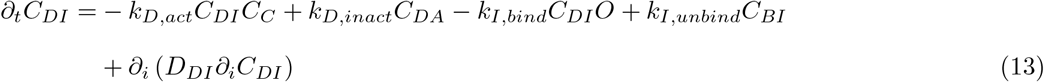

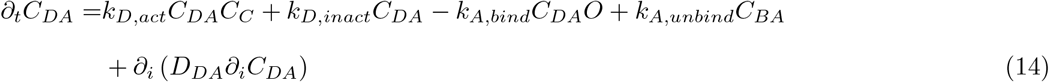

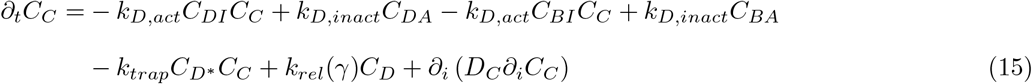

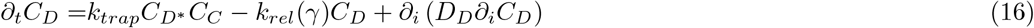

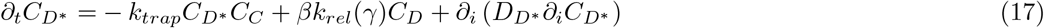

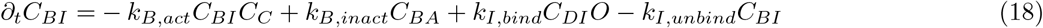

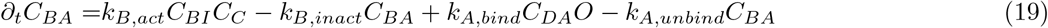

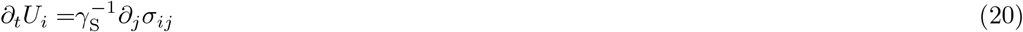

In Equation 17 we introduce the parameter *β* to control the degree of degradation of DMNP-EDTA chelators upon light-induced cleavage. If *β* = 1, then every molecule of DMNP-EDTA which release Ca^2+^ after photolysis is available again to bind a new Ca^2+^ ion. If *β* = 0, then every such DMNP-EDTA molecule is destroyed upon photolysis and hence does not enter the pool of available chelators.

The derivative ∂_*j*_*σ*_*ij*_ is

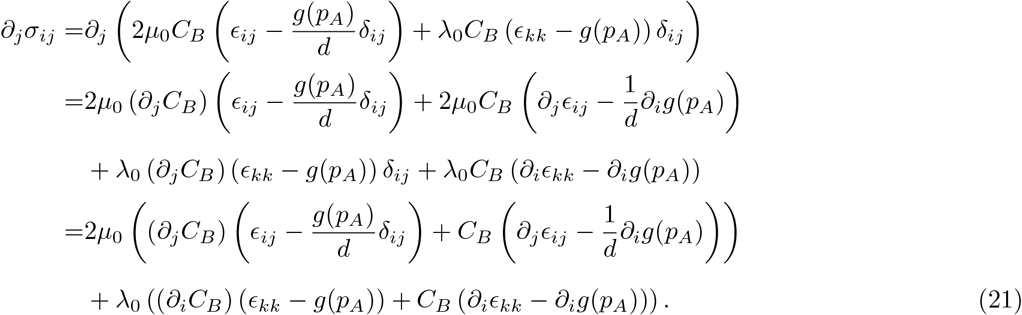

The derivative ∂_*i*_*g*(*p*_*A*_) is

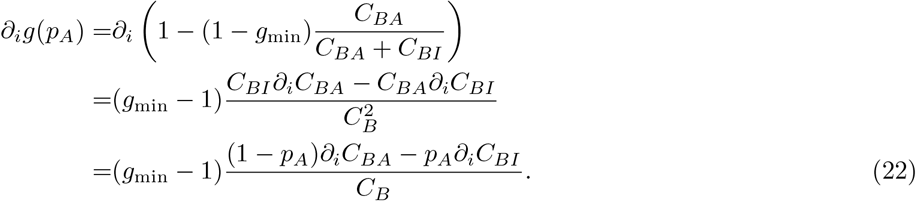

For azimuthally symmetric light protocols, we convert these equations to cylindrical coordinates and solve for only the radial dependence.

Unless otherwise specified we use the spatially uniform initial conditions with **u**(**r**, 0) = **0**. We set all concentrations to zero except *C*_*DI*_ (**r**, 0) = 1.25 mM, *C*_*D*_(**r**, 0) = *C*_*D*∗_ (**r**, 0) = 20 mM, and we uniformly nucleate the first growth of the bound Tcb2 network by setting *C*_*BI*_ (**r**, 0) = 0.05 mM.

#### 3. Light field

We decompose the non-autonomous function *γ*(**r**, *t*) = *γ*_*r*_(**r**)*γ*_*t*_(*t*), which mimics spatiotemporal light fields from the DMD, into a part depending only on position and a part depending only on time (Fig. S6). For circular profiles, the spatial part is a decreasing sigmoid function of the radial distance *r* = ||**r**||,

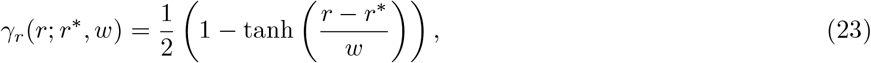

having an offset *r*^∗^ (the illumination radius) and a width *w* (the spread of light at the cutoff). Unless otherwise noted we set *w* = 4 *µ*m throughout. The star pattern is implemented in 2D using a tunable parametric formula for the offset *r*^∗^ as a function of polar angle. The time-dependent part *γ*_*t*_(*t*) is also constructed using sigmoidal curves with finite width of 0.5 s. In the pulse protocol we chain several sigmoidal bumps with pulse length of 1 s and a cycle length of 30 s, while in the continuous protocol we use a single bump with duration 100 s.

#### 4. User-defined concentration profile

In Fig. 4 of the main text we show results using a user-defined concentration profile for *C*_*B*_(*r*) to systematically explore the role of boundary accumulation on reversing the contraction direction. The user-defined function is (cf. Equation 23)

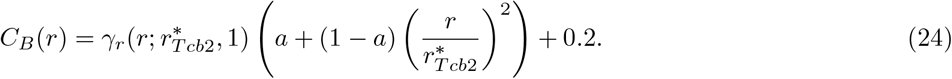

During simulations with this imposed concentration profile, we mimic the effect of light by directly setting the ratio 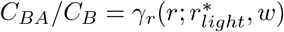 using Equation 23, and we then solve for the radial displacement fields *U*_*r*_(*r, t*) (cf. Equation 20) after a 1 s pulse of light.

#### 5. Parameterization

In Tables S4, S5, and S6 we provide the default parameters used in simulation. We set the initial concentrations in Table S4 according to the experimental conditions. We constrained the reaction rates in Table S5 where possible by available literature data, such as the DMNP-EDTA release rate of Ca^2+^ under light exposure [38, 39]. We set the unknown values by exploring parameter space to find reasonable agreement with experimental results. We estimated diffusion constants in Table S6 in order of magnitude using approximate knowledge of the size of the different molecules.

**FIG. S6.**
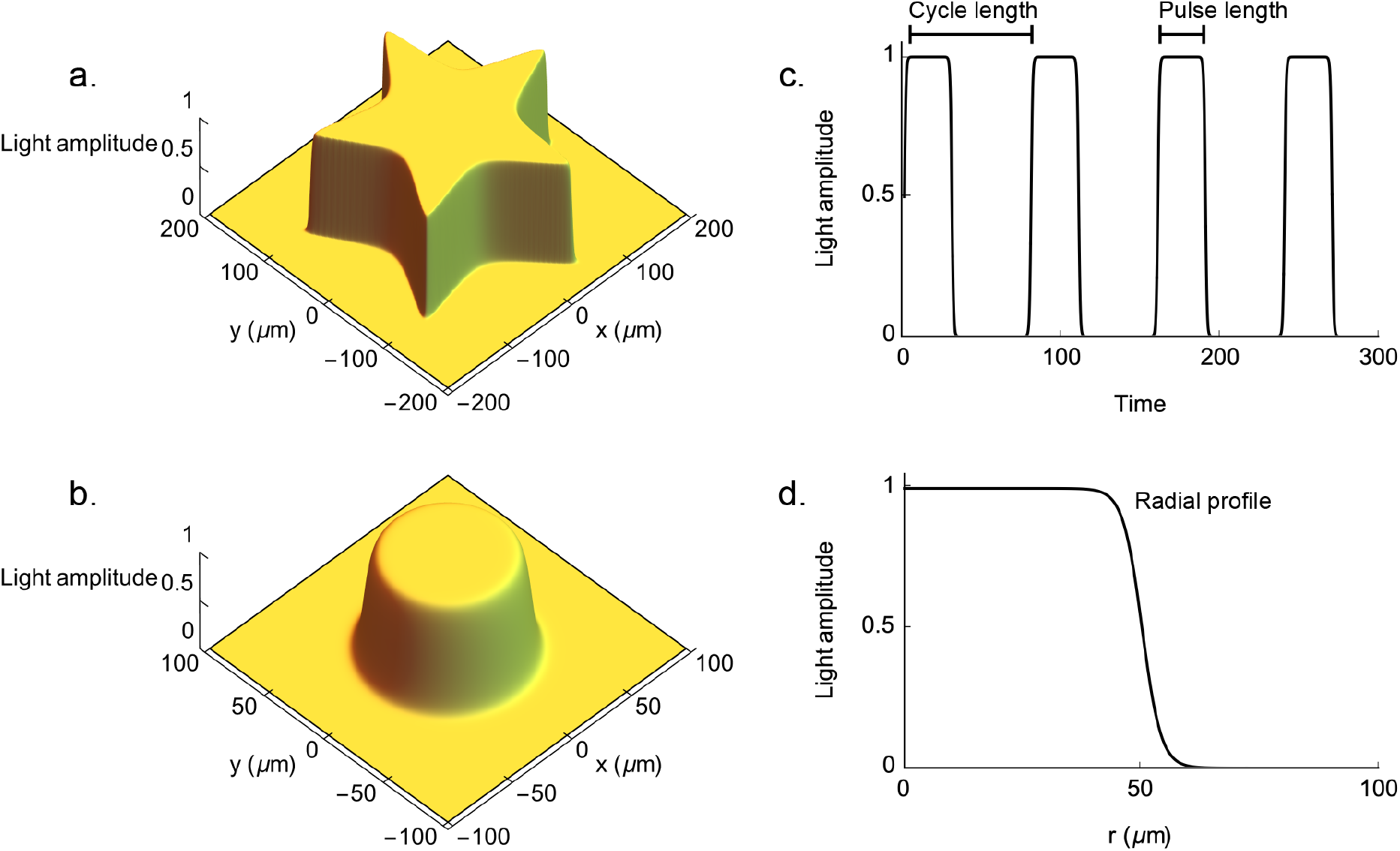
Light protocol. (a) Spatial pattern *γ*_*r*_ (**r**) of the star-shaped illumination used in simulation. (b) Spatial pattern *γ*_*r*_ (**r**) of the circular illumination used in simulation, here with a radius of 50 *µ*m. (c) Illustration of a pulsed temporal protocol *γ*_*t*_(*t*), with the the cycle length and pulse length labeled. (d) For circularly symmetric illumination patterns, cylindrical coordinates are used and only the radial part of *γ*_*r*_ (*r*) is considered. This profile corresponds to Equation 23 with *r*^∗^ = 50 *µ*m and *w* = 4 *µ*m.

**TABLE S4.** Initial concentrations used in simulation.

| Parameter | Value | Units |
| --- | --- | --- |
| $C_C$ (diffusing $\text{Ca}^{2+}$) | 0 | mM |
| $C_{DI}$ (diffusing inactivated Tcb2) | 1.25 | mM |
| $C_{BI}$ (bound inactivated Tcb2) | 0.05 | mM |
| $C_{DA}$ (diffusing activated Tcb2) | 0.0 | mM |
| $C_{BA}$ (bound activated Tcb2) | 0.0 | mM |
| $C_D$ (DMNP-EDTA with $\text{Ca}^{2+}$) | 20.0 | mM |
| $C_{D^*}$ (DMNP-EDTA without $\text{Ca}^{2+}$) | 20.0 | mM |

#### 6. Varying the model

The continuum model is under-constrained by the available data. While future work will focus on kinetic parameter estimation, rheological measurements, and other characterizations to better constrain it, our present goal is not quantitative accuracy but rather to capture the key dynamical features observed experimentally, providing qualitative agreement that yields physical insight. In particular, the model recapitulates the localization of contractile force near the network periphery, the accumulation of bound Tcb2 near the periphery, and the occasional reversal of contraction direction toward the periphery from the inside. In this section we alter certain assumptions of our minimal model to verify that they do not change these qualitative results.

**TABLE S5.** Chemical reaction rates used in simulation. ^∗^ - These values are based on Refs. 38, 39. To avoid stiff numerical integration steps, both the binding and unbinding rates have been decreased by the same factor (preserving their ratio) and keeping this process fast compared to others in the system.

| Parameter | Value | Units |
| --- | --- | --- |
| $k_{D,act}$ (binding of Ca to diffusing Tcb2) | 50 | $(\text{mM} \cdot \text{s})^{-1}$ |
| $k_{D,inact}$ (unbinding of Ca to diffusing Tcb2) | 20 | $\text{s}^{-1}$ |
| $k_{B,act}$ (binding of Ca to bound Tcb2) | 50 | $(\text{mM} \cdot \text{s})^{-1}$ |
| $k_{B,inact}$ (unbinding of Ca to bound Tcb2) | 20 | $\text{s}^{-1}$ |
| $k_{trap}$ (trapping of Ca by DMNP-EDTA when light is on or off) | 30* | $(\text{mM} \cdot \text{s})^{-1}$ |
| $k_{rel}(0)$ (release of Ca by DMNP-EDTA when light is off) | 0 | $\text{s}^{-1}$ |
| $k_{rel}(1)$ (release of Ca by DMNP-EDTA when light is on) | 700* | $\text{s}^{-1}$ |
| $k_{I,bind}$ (binding of inactivated Tcb2 to bound Tcb2) | 0.2 | $(\text{mM} \cdot \text{s})^{-1}$ |
| $k_{I,unbind}$ (unbinding of inactivated Tcb2 from bound Tcb2) | 0.5 | $\text{s}^{-1}$ |
| $k_{A,bind}$ (binding of activated Tcb2 to bound Tcb2) | 0.2 | $(\text{mM} \cdot \text{s})^{-1}$ |
| $k_{A,unbind}$ (unbinding of activated Tcb2 from bound Tcb2) | 0.5 | $\text{s}^{-1}$ |
| $\beta$ (fraction of DMNP-EDTA returning to available pool after release) | 0.2 | — |
| $C_{sat}$ (maximum concentration of bound Tcb2) | 5 | mM |

**TABLE S6.** Diffusion constants used in simulation.

| Parameter | Value | Units |
| --- | --- | --- |
| $D_C$ ($\text{Ca}^{2+}$) | 300 | $\mu\text{m}^2/\text{s}$ |
| $D_D$ (DMNP-EDTA with $\text{Ca}^{2+}$) | 100 | $\mu\text{m}^2/\text{s}$ |
| $D_{D^*}$ (DMNP-EDTA without $\text{Ca}^{2+}$) | 100 | $\mu\text{m}^2/\text{s}$ |
| $D_{DA}$ (activated Tcb2) | 10 | $\mu\text{m}^2/\text{s}$ |
| $D_{DI}$ (inactivated Tcb2) | 10 | $\mu\text{m}^2/\text{s}$ |

**TABLE S7.**
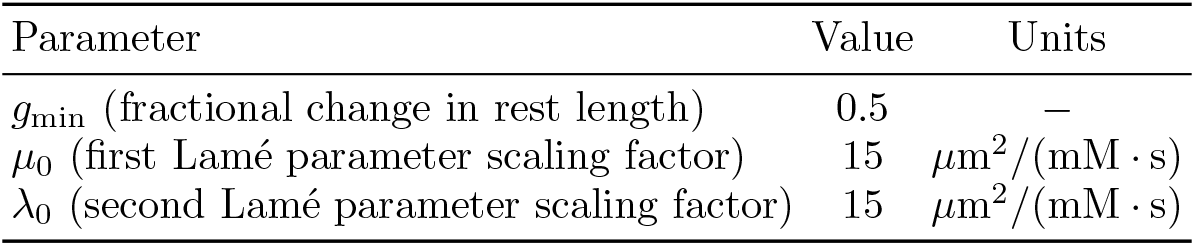
Mechanical parameters used in simulation.

We first consider how the mechanical contraction dynamics depend on the manner in which stiffness (the Lamé parameters *µ, λ*) depends on Tcb2 density. Based on typical Ashby plots relating stiffness to material density [36], our simple assumption in the default model is that *µ*(*C*_*B*_) ∝ *C*_*B*_ and similarly for *λ*(*C*_*B*_). We generalize this to a power law dependence 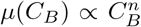 such that at the saturation density *C*_*B*_ = *C*_*sat*_ all stiffness models coincide at We thus consider the form

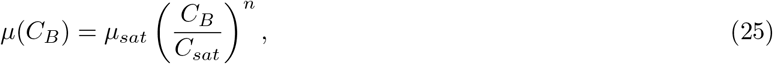

and similarly for *λ*(*C*_*B*_). See SI Figure S7a for an illustration of this curve. We note that in the elastic equations of motion the spatial gradients of stiffness become more complicated using these power law models than in the default case when *n* = 1. We find different powers of *n* yield quantitatively different contraction dynamics, yet the qualitative features of the radial velocity curves (such as peaks near the boundary and presence of radially outward components) are preserved for each choice.

**FIG. S7.**
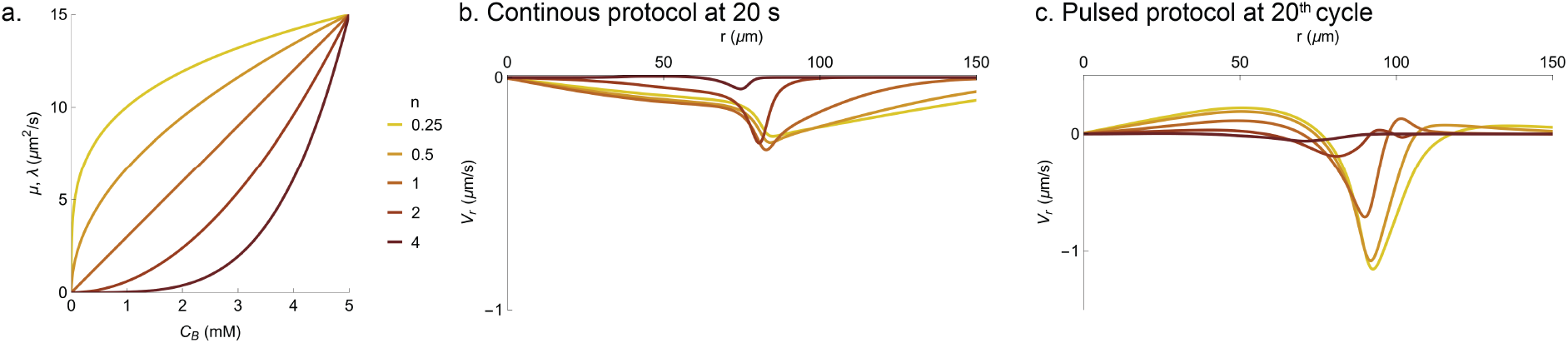
Comparison of simulated network dynamics for different values of *n* in the elastic model. (a) Plots of the Lamé parameters *µ* and *λ* as a function of bound Tcb2 concentration *C*_*B*_ as *n* is varied among the values 0.25, 0.5, 1, 2, and 4. (b) Radial profiles of radial velocity *V*_*r*_ for simulations with different values of *n* for the continuous light protocol at 20 s. Colors correspond to the legend in panel a. (c) Same as panel b, but for the 20^th^ cycle of the pulsed protocol.

Next, we test the effect of different values of the saturating bound Tcb2 concentration *C*_*sat*_. Above this threshold, we assume that remaining Tcb2 binding sites are sterically inaccessible, causing the network binding rate to drop to zero. Although our phenomenological model for Tcb2 network nucleation and growth is simplified and will require refinement with improved data constraints, it is sufficient here to capture the essential features of growth via self-limiting aggregation. In SI Figure S8 we show that variations in this parameter around its default value of 5 mM yield negligible differences in the diffusing Ca^2+^ profiles (as Ca^2+^ binding by Tcb2 is taken to not depend on Tcb2-Tcb2 binding), and only minor quantitative differences in the profiles of bound Tcb2 and radial velocity.

**FIG. S8.**
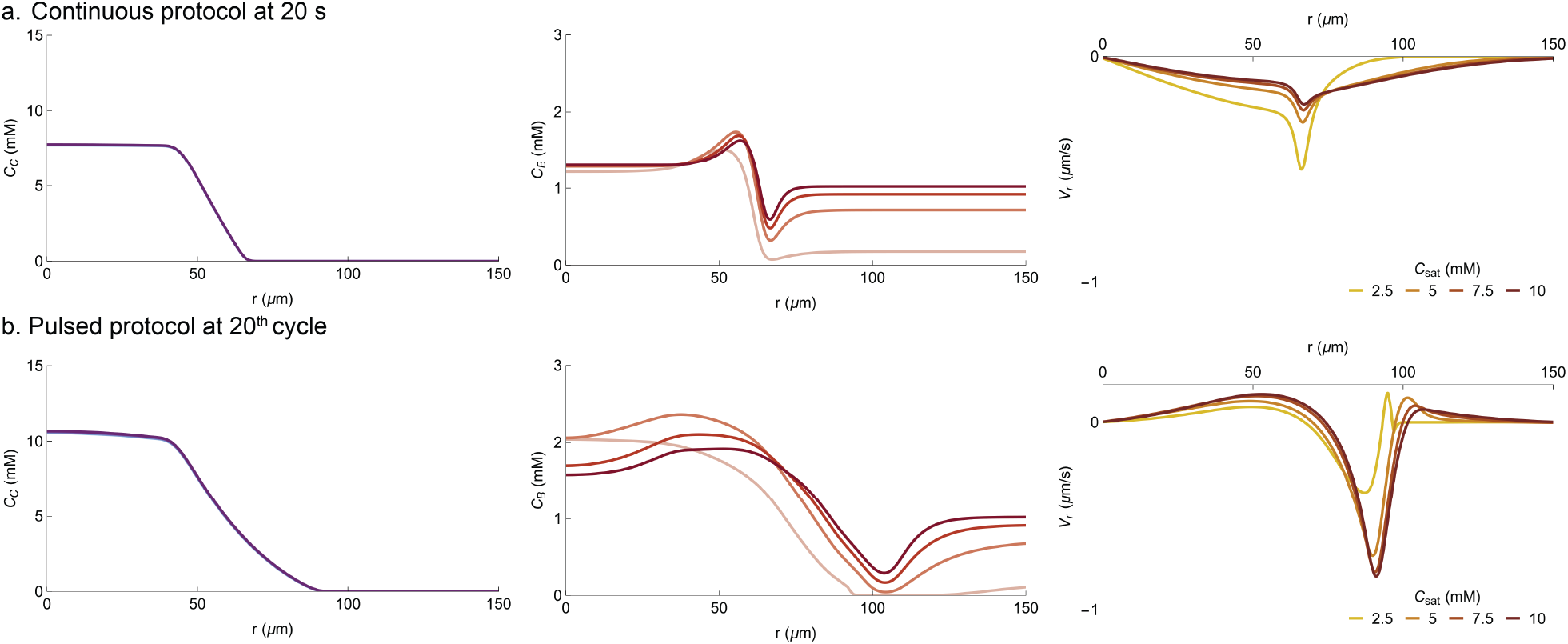
Comparison of simulated network dynamics for different values of *C*_*sat*_ in the chemical model. (a) Radial profiles of diffusing Ca^2+^ concentration *C*_*C*_, bound Tcb2 concentration *C*_*B*_, and radial velocity *V*_*r*_ for simulations at 20 s during the continuous light protocol for different values of the parameter *C*_*sat*_. (b) Same as panel a, but for the 20^th^ cycle of the pulsed protocol.

Finally, we test whether advection of chemical species, which we neglect in the default model, has a significant effect. We run simulations with advective terms in the equations of motion for all chemical species and compare the resulting behavior with the default model which neglects advection. The results indicate that advection produces a small systematic correction to the diffusing Ca^2+^ and radial velocity profiles and negligible correction to the bound Tcb2 profiles, for both the continuous and pulsed light protocols; see SI Figure S9. Advection thus appears to not qualitatively affect the conclusions of the model.

To understand this, we estimate the Péclet number for the different diffusing species in our system (Ca^2+^, DMNP, and Tcb2) using their diffusion constants of 300, 100 and 10 *µ*m^2^*/*s respectively (these order-of-magnitude values are obtained based on molecular weights and are used in our simulations). The maximum contractile velocity is approximately 1 *µ*m*/*s, and as a length scale we take the typical CAR width, which is roughly 20 *µ*m. We thus have Péclet estimates of 1*/*15, 1*/*5, and 2 for Ca^2+^, DMNP, and Tcb2. These values suggest that advection may introduce minor quantitative corrections to the observed dynamics but should not overwhelm the physics. Indeed this is what we observe when we include advection in the dynamics.

**FIG. S9.**
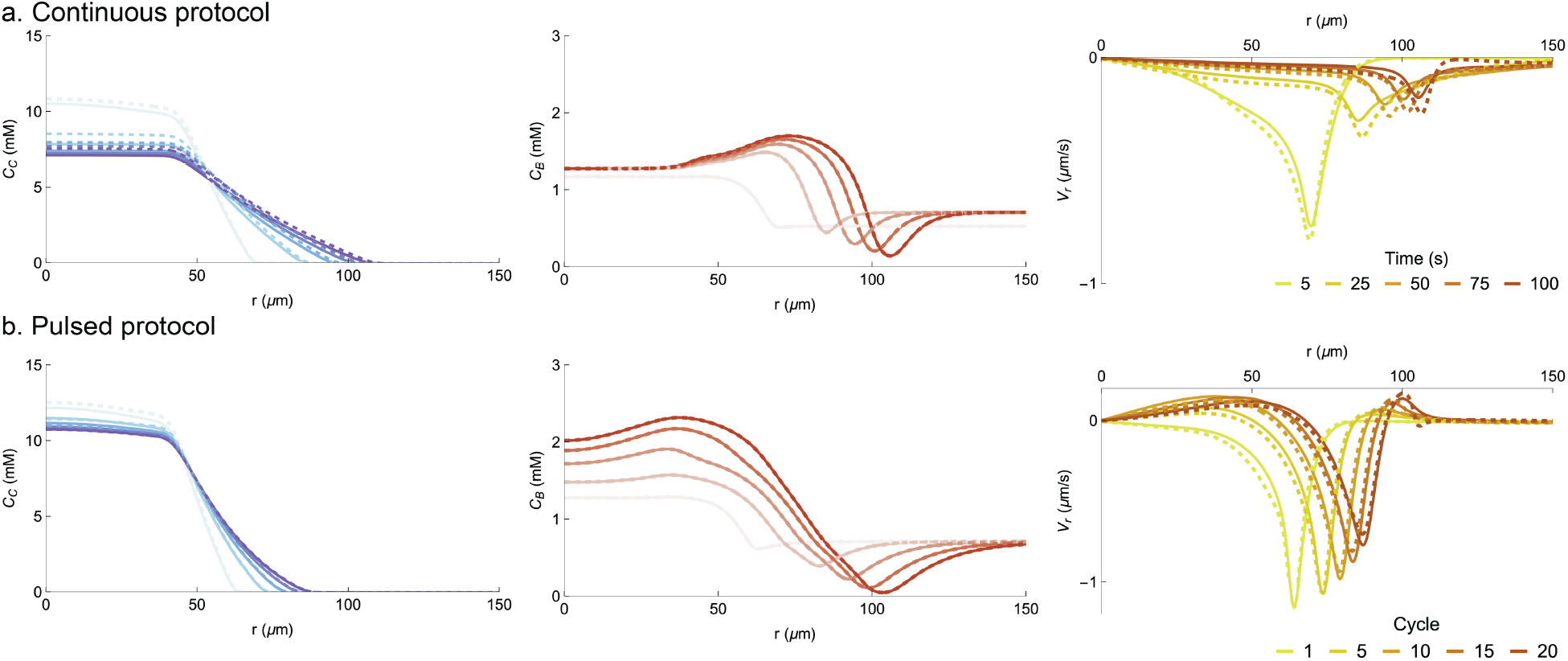
Comparison of simulated network dynamics with and without advection. (a) Radial profiles of diffusing Ca^2+^ concentration *C*_*C*_, bound Tcb2 concentration *C*_*B*_, and radial velocity *V*_*r*_ for simulations with (dotted lines) and without (solid lines) the inclusion of advection in the dynamics for the chemical species. The colors range from light to dark as time increases during the simulation of the continuous light protocol, as indicated by legend on the right. (b) Same as panel a, but for the pulsed protocol.

We conclude that while the model lacks quantitative accuracy, its qualitative features of boundary accumulation, contraction localization, and contractile direction reversal are fairly robust to the modeling choices made here.

### F. Reinforcement learning

#### 1. Problem definition

Reinforcement learning (RL) is a set of techniques for training an agent to take actions in an environment so that it can optimize a reward function [40]. For us, the agent has control over the spatial light pattern which illuminates a Tcb2 network, and its goal is to move a certain spot in the network to a given position and then hold it there. Specifically, in radial coordinates we pick a position *r*_0_ and imagine attaching a “spring” to between the radial displacement at this point and a goal displacement *U*_goal_. The “spring force” experienced at *r*_0_ is *F*_*s*_ = *k*_*s*_(*U*_goal_ − *U* (*r*_0_, *t*)), where *U* (*r*_0_, *t*) is the displacement at coordinate *r*_0_. Throughout this section all displacements are radial displacements but we drop the subscript *r* to simplify notation. We want the agent to shine light in a way that mimics the effect of this imaginary spring, thereby causing the point to move to position *r*^∗^ and stay there.

The light field’s spatial pattern is a sigmoidal decreasing function parameterized as in Equation 23. The width is fixed at *w* = 4 *µ*m, and the agent has control over the amplitude *γ*_*t*_ ∈ [0.01, 0.16] and offset *r*^∗^ ∈ [*r*_0_ − 20, *r*_0_ + 20] *µ*m.

An episode of training is set as a fixed duration of time *T* = 25 s. The numerical integration time step is *dt* = 2×10^−4^ s, and the agent updates the light field parameters every 0.5 s. It chooses new parameters according to a neural network function which maps the state of the system, which it sees as (*U* (*r*_0_, *t*) − *U*_goal_)*/U*_goal_, into values of *γ*_*t*_ and *r*^∗^. We divide by *U*_goal_ so the result takes values of roughly order 1.

During an increment Δ*t*, under the newly set of parameters chosen by the policy, the displacement will change to *U* (*r*_0_, *t* + Δ*t*). If the point were really attached to a spring, then the change in *U* during this time would be

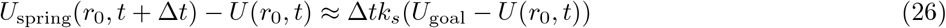

where we have assumed overdamped dynamics for the spring with a drag coefficient that we absorb into the definition of *k*_*s*_, which we set as *k*_*s*_ = 0.01 s^−1^ throughout. Because the agent is not perfectly imitating the imaginary spring, the real displacement will be *U* (*r*_0_, *t* + Δ*t*) ≠ *U*_spring_(*r*_0_, *t* + Δ*t*). To encourage the agent to better imitate the spring, we thus provide rewards after each Δ*t* as

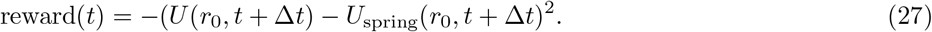

This reward is used to provide updates to the function which maps the state *U* (*r*_0_) into actions, sigmoidal light amplitude *γ*_*t*_(*t*) and offset *r*^∗^(*t*), so that the reward is eventually maximized. See Ref. 41 for additional details on our similar application of RL to control active nematic defects [42].

We emphasize that the agent only sees the the current displacement *U* (*r*_0_, *t*) when deciding what light parameters to use, which is a very coarse view of the system state. That the agent nonetheless learns good policy functions given this information, at least for moderately long simulation trajectories, suggests that despite the complicated physics of the full system, a coarse projection of the system state provides enough information to guide the system’s dynamics.

#### 2. Algorithm

To train the RL agent, i.e., learn a policy which maps *u*(*r*_0_) into light field parameters *γ*_*t*_ and *r*^∗^, we use a variant of the actor-critic algorithm [40] called deep deterministic policy gradient (DDPG) [43, 44], which is well-suited for continuous actions in deterministic environments. We combined our custom numerical integrator of the Tcb2 dynamics with the DDPG implementation provided by the Julia package ReinforcementLearning.jl [45]. The actor and critic neural networks each have 1 hidden layer with 32 neurons, which are trained using stochastic batches of 128 (state, action, reward, termination bool) tuples that are collected during training. The agent first chooses actions stochastically for ∼ 9, 600 steps after which it begins to use its learned policy. The neural networks are updated using stochastic batches of 128 tuples every 10^th^ step. We use a discount factor *γ* = 0.99 and weight transfer factor *τ* = 0.995 [44]. The neural networks are trained using the ADAM optimizer with a learning rate of 0.005 and a weight norm clip of 0.5.

#### 3. Latency

To address robustness of the RL algorithm against common experimental challenges, we include *in silico* a latency between the current state of the system (when the actions are applied) and the RL agent’s view of it. This mimics the finite processing time required to do image processing and evaluate the control policy in real time in our envisioned hybrid experimental and software platform (see Fig. 7 of the main text and Ref. 46). In Fig. S10 we show trajectories of trained policies for four levels of latency, including a severe latency of 3 *s*. In all cases the RL algorithm is able to learn a policy which brings the displacement to its target value, indicating that RL is a flexible enough approach to counteract latencies and learn effective feedback control.

### G. Area detection

#### 1. Measuring area experimentally

To segment the protein network from the background in the differential interference contrast (DIC) microscopy videos, we implemented the AI model Segment Anything [47] and used a modified version of the code from GitHub for video tracking [48]. To enhance both segmentation accuracy and speed for Tcb2 network (single-object) detection in DIC videos, we employed a prompt-based initialization method. We initialized the model with a prompt point at the center of the image, which is aligned with the center of the light activation area and the protein network. In the Supplementary Videos using this analysis, the red curve indicates the AI-detected protein boundary. We manually reviewed the videos to ensure the accuracy of the segmentation.

**FIG. S10.**
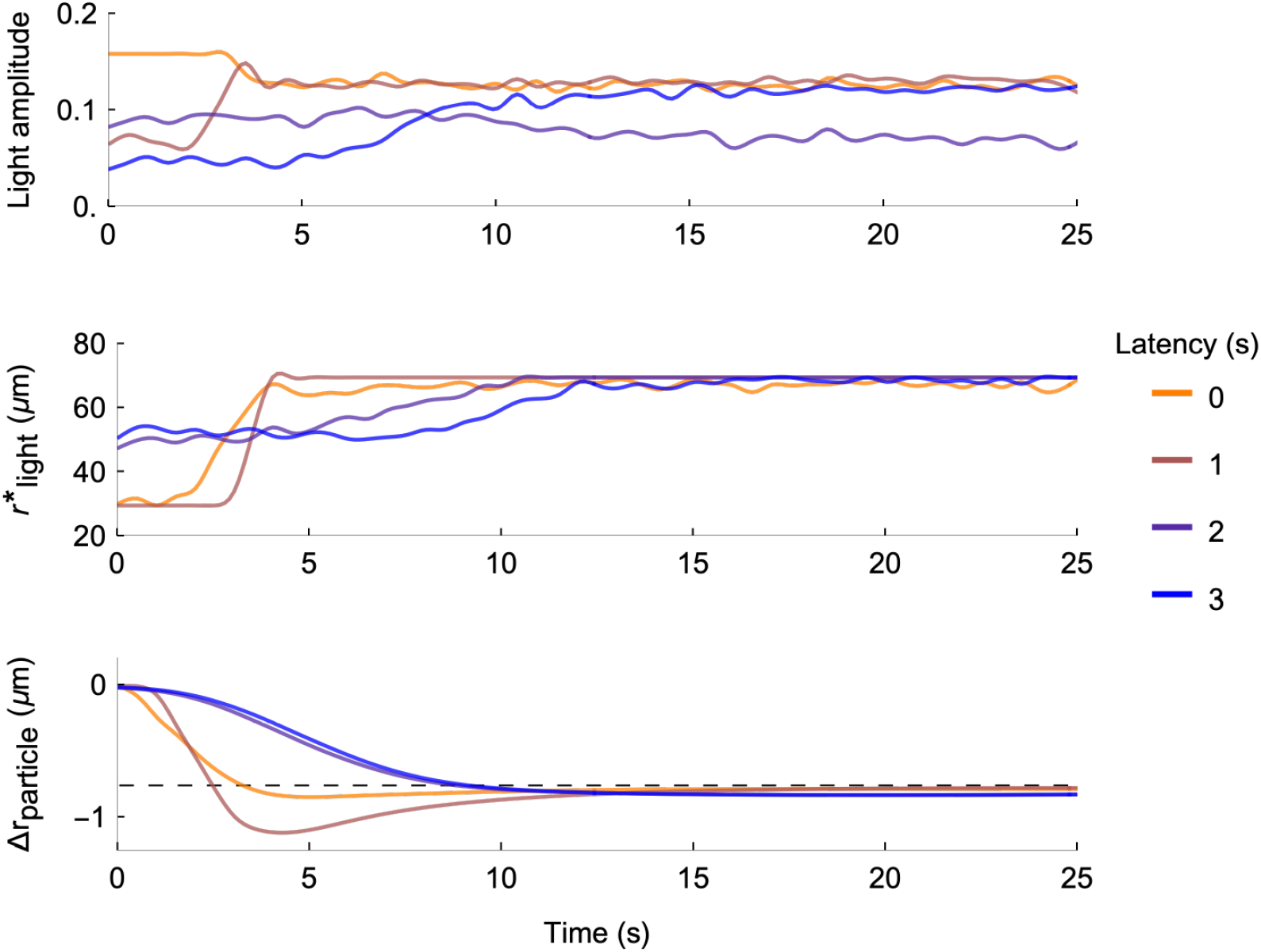
Including latency in the feedback control. Trajectories of the light amplitude, light position, and displacement at the target location in the system for four levels of latency between the current state of the system and the RL agent’s view of it.

#### 2. Measuring area in simulation

To track the area of simulated Tcb2 networks, we defined a threshold on the simulated function *C*_*B*_(*r, t*) = *C*_*BA*_(*r, t*) + *C*_*BI*_ (*r, t*) of total bound Tcb2. Using thresholds in the range 0.75 − 0.9 *µ*M, we define the radius of the network as the most distant value of *r* which exceeds the threshold. We note that this approach is different from the approach for tracking area in experiments, which does not have direct access to the value of *C*_*B*_(*r, t*) and relies on an adaptive AI method for segmentation. Despite these differences in approach, we find semiquantitative agreement between simulation and experimental results for the areal growth of the network.

## II. SUPPLEMENTARY RESULTS

### A. Varying the illumination area

Here we describe how properties of the network growth and contraction depend on the size of the circular illumination area, in both pulsed and continuous light protocols.

#### 1. Effect on the growth

In Fig. S11 we show experimental and simulation results for how the network’s growth in time varies with the illumination size using the continuous light protocol. We represent the network size using its effective radius *R*, obtained from its measured area *A* as 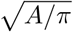. The networks grow faster with a larger illumination size in both the experiments and simulations.

We note that the simulations do not fully capture the degree to which the growth rate increases with the illumination size. Extensive parameter tuning failed to alleviate this disagreement, which we speculate may be due to an overly simplified model of the bound Tcb2 network’s nucleation dynamics as well as unaccounted-for contributions from chemical buffers in the experimental solution. We leave resolution of this quantitative discrepancy as an avenue for future work, which may benefit from more dedicated measurements of the initial network formation.

The curves of effective radius *R*(*t*) suggest that the growth of the network beyond the illuminated region is primarily determined by diffusion. The area covered by molecules diffusing from a source grows in time as *A*(*t*) ∝ *Dt* with *D* the effective diffusion constant. The corresponding radius thus grows as 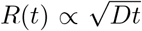. Accounting for an initial radial size (coming from the finite illumination region), we expect a diffusion-limited network’s radius to grow in time according to the functional form

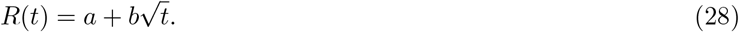

This functional form fits both the experimental and simulated growth curves reasonably well. In the experiments, the value *a* matches roughly with the initial illumination radius, although in simulations the growth occurs so rapidly in the beginning that the fitted value of *a* is larger than this illumination radius. Despite this difference between experiments and simulations, the fitted values of *a* both scale linearly as a function of illumination diameter and with similar slopes. The fitted values of *b* for experiments and simulations similarly differ in numerical values but scale in the same way with the illumination diameter.

In Fig. S12 we show experimental and simulation results for how the network’s growth in time varies with the illumination size using the pulsed light protocol. As with the continuous protocol, the rate at which the network grows increases with illumination diameter. The growth rate averaged over cycles is approximately linear in both the experimental and simulated results, although we find that the simulations do not quantitatively match the slopes of these linear growth dynamics. For the initial cycles, the network does not accumulate enough bound Tcb2 to leave a detectable area, either through the adaptive AI-based segmentation method used to analyze experiments or through the fixed concentration threshold-based method to analyze simulation results. For the smallest illumination diameter in experiments, the network fails to nucleate a detectable area after many repeated cycles, suggesting that insufficient Ca^2+^ is released by the light illumination to persistently grow the network. We note that we repeated each experiment two to four times per condition but for clarity we only plot one trial per condition in Figs. S11, S12.

In Fig. S13 we show simulation kymographs of both the bound Tcb2 and Ca^2+^ radial profiles as a function of time across conditions of illumination diameter and light protocol. We observe qualitatively similar behavior as the illumination diameter is varied.

**FIG. S11.**
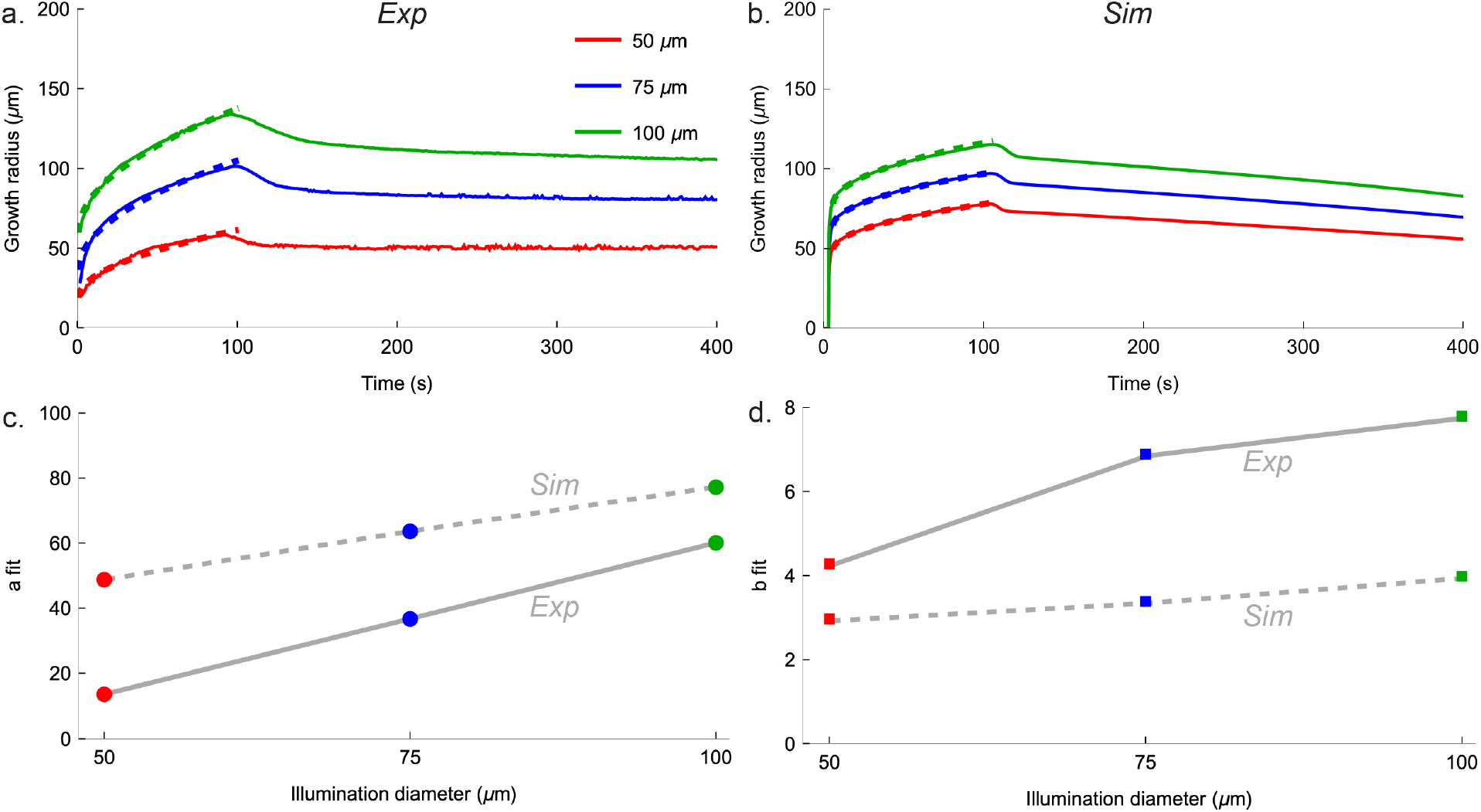
Network growth dynamics with varying illumination diameters for the continuous protocols. (a) Experimentally measured growth radius as a function of time for different illumination diameters. Dashed curves indicate fits of Equation 28 to the 100 s of light being held on. We repeat each experiment 2-4 times but only plot one here for visibility. (b) Simulated growth radius as a function of time for different illumination diameters. (c) Fitted values of *a* in Equation 28. (d) Fitted values of *b*.

**FIG. S12.**
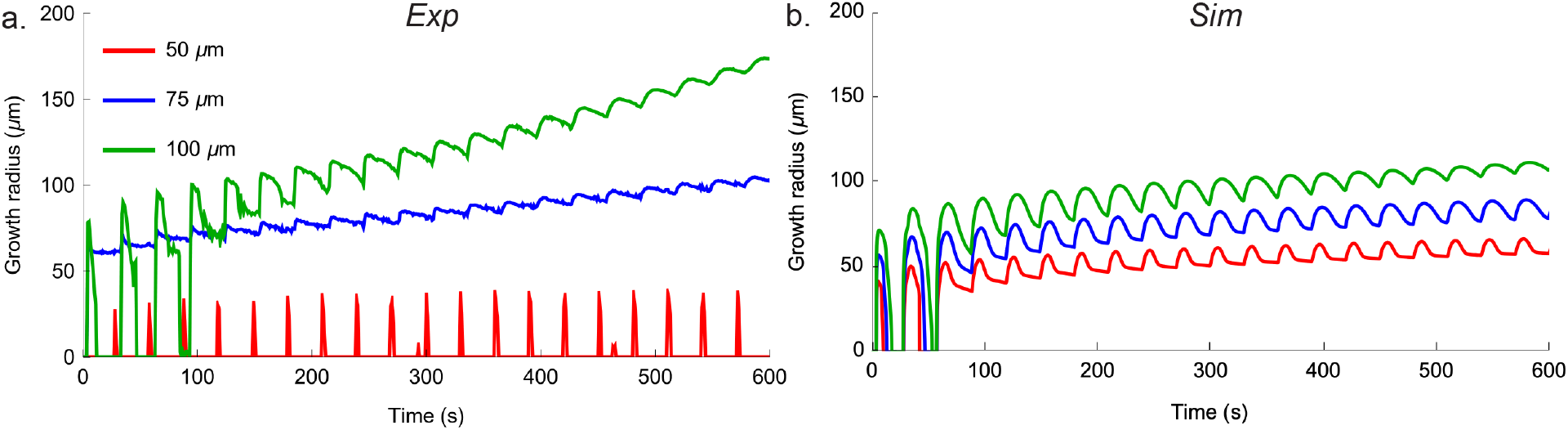
Network growth dynamics with varying illumination diameters for the pulsed protocols. (a) Experimentally measured growth radius as a function of time for different illumination diameters. (b) Simulated growth radius as a function of time for different illumination diameters.

#### 2. Effect on the contractility

In Fig. S14 we show experimental and simulation results for the network’s radial velocity as a function of time, as we vary the illumination diameter and the light protocol. In experiments we observe qualitatively similar behavior of the network as the illumination size increases, with the exception of the smallest illumination size which fails to nucleate fully. The radial velocity data for this condition is noisy and we do not discern any clear trends.

**FIG. S13.**
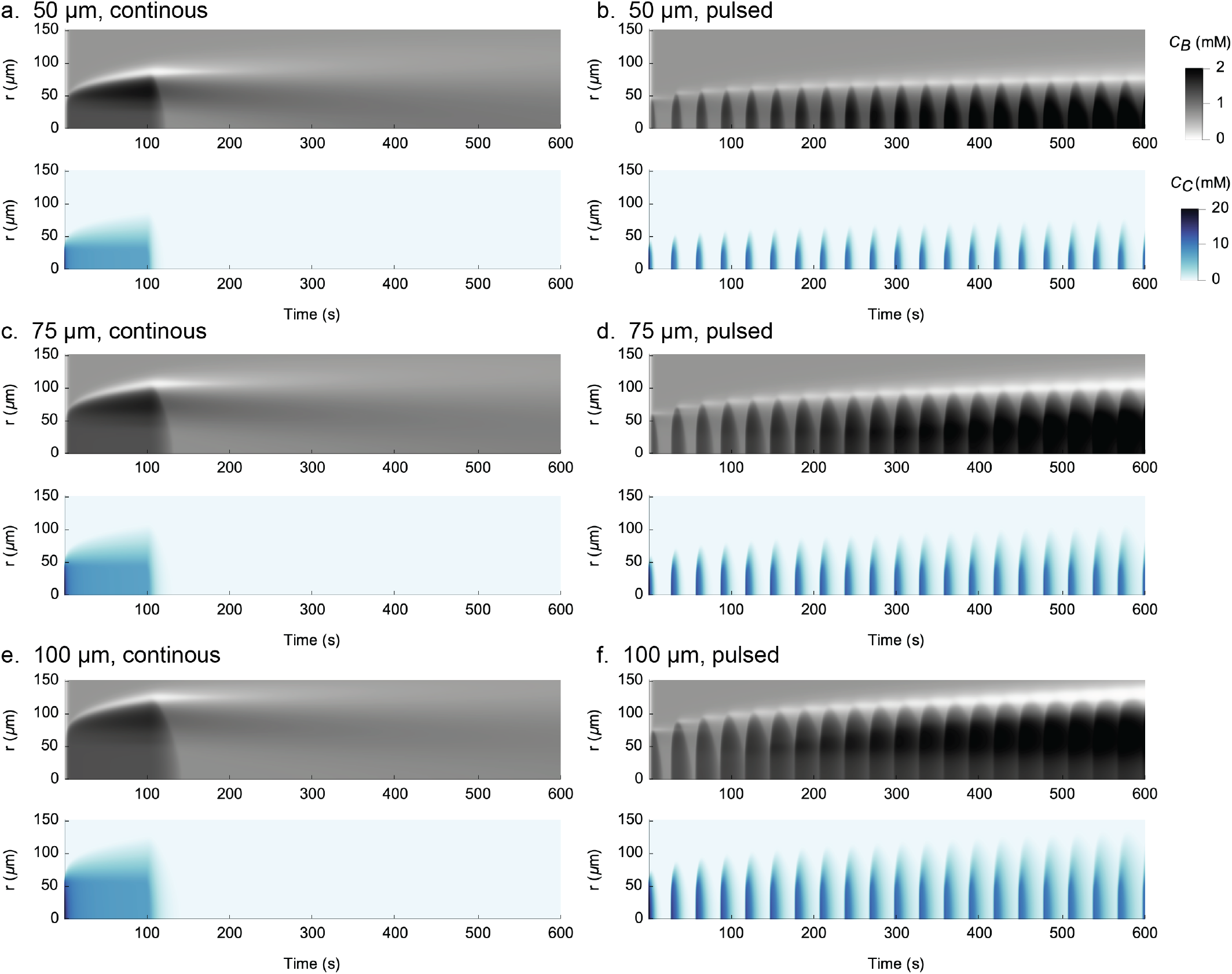
Network concentration dynamics with varying illumination diameters and light protocols. (a) Simulated kymographs for the bound Tcb2 and diffusing Ca^2+^ concentrations as a function of radial position for the continuous light protocol, and with an illumination diameter of 50 *µ*m. (b) Same as panel a, for the pulsed light protocol. (c) Same as panel a, for an illumination diameter of 75 *µ*m. (d) Same as panel c, for the pulsed light protocol. (e) Same as panel a, for an illumination diameter of 100 *µ*m. (f) Same as panel e, for the pulsed light protocol.

We find slight discrepancies between the experimental measurement and simulation results for this data. We noted above that the network growth dynamics are not in complete agreement alignment, leading to a difference in the spatial extent of radial velocity. In addition, we find in the experiments that during the continuous protocol the interior of the network has a small radially outward velocity for illumination diameters of 75 and 100 *µ*m, while this feature is not observed in the simulations. We do observe radially outward velocities in the network’s interior during the pulsed protocols, which we explain in Fig. 4 of the main text as resulting from an accumulation of bound Tcb2 concentration near the network periphery. Although this accumulation also exists in simulations of the continuous protocol, it does not give rise to an outward velocity. This suggests that further refinement of the model assumptions, such as the assumptions underlying nucleation and chemical growth, isotropy of the autogeneous strain due to Ca^2+^ activation, linear growth rate of the Lamé parameters with *C*_*B*_, or parameter choices are needed in the future to achieve more quantitative agreement with experiments. We leave these issues to future work, and note that the current model suffices to semi-quantitatively explain several other features of the contraction dynamics as described in the main text.

**FIG. S14.**
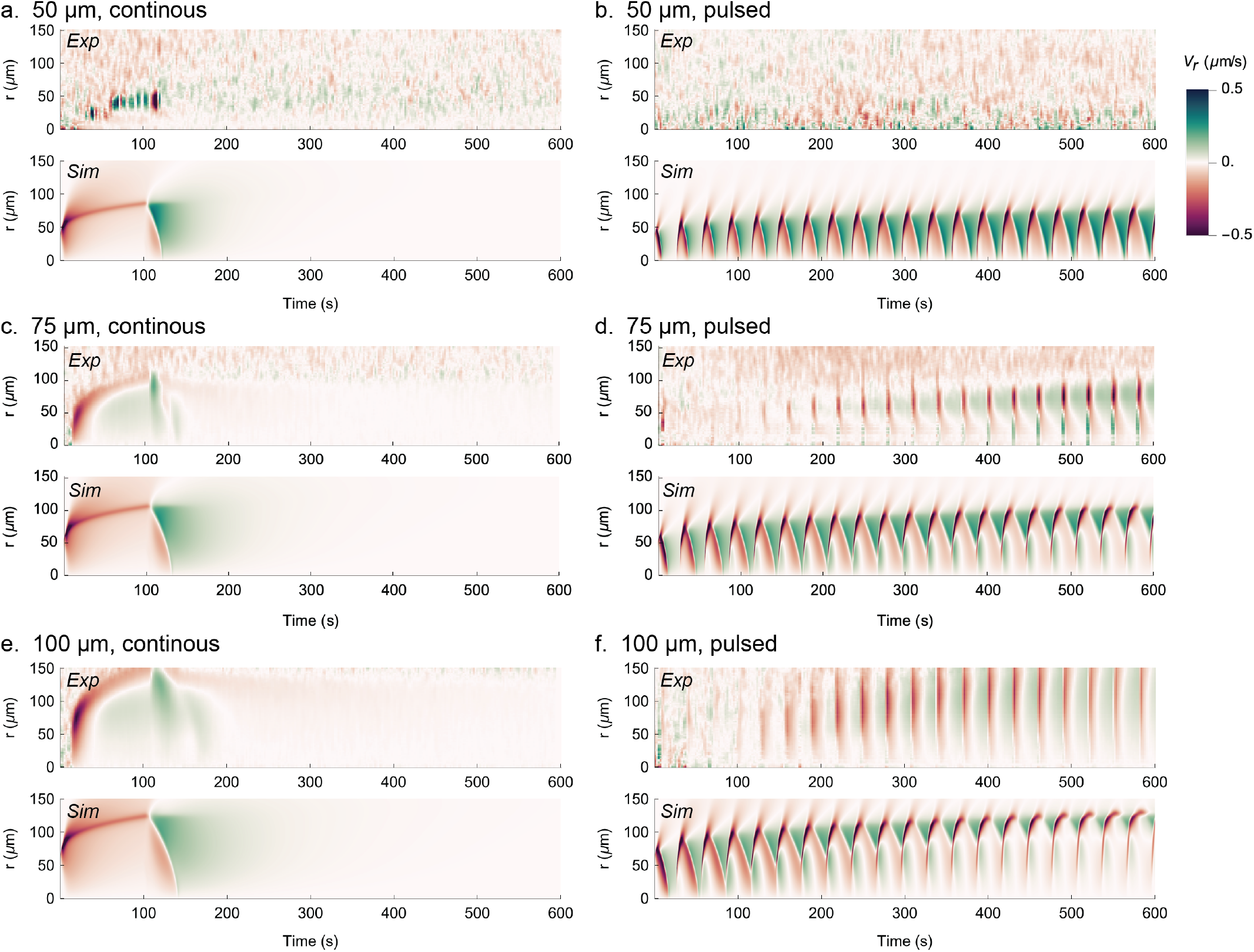
Network contraction dynamics with varying illumination diameters and light protocols. (a) Experimental and simulated kymographs for the radial velocity as a function of radial position for the continuous light protocol, and with an illumination diameter of 50 *µ*m. (b) Same as panel a, for the pulsed light protocol. (c) Same as panel a, for an illumination diameter of 75 *µ*m. (d) Same as panel c, for the pulsed light protocol. (e) Same as panel a, for an illumination diameter of 100 *µ*m. (f) Same as panel e, for the pulsed light protocol.

### B. Pulse repeatability experiments over many cycles

To test the degree of repeatability of light-induced mechanical contraction, we performed experiments with large numbers of pulsation cycles. Figure S15 presents DIC kymographs (left) displaying chemical assembly and disassembly dynamics, alongside corresponding radial velocity-colored kymographs (right) that visualize contraction speed and amplitude. We performed cycling tests using illumination regions of three diameters: 50 *µ*m (∼150 cycles; SI Video Part 2 Section VI), 75 *µ*m (∼70 cycles; SI Video Part 2 Section V), and 100 *µ*m (∼30 pulses). The maximum number of observable cycles was constrained by our 20× objective’s 300 × 300 *µ*m field of view, as contracting regions eventually extended beyond the imaging area. Figure S16 quantifies these dynamics through measurements of both the active contracting radius (defined as regions with velocities *>*0.2 *µ*m/s) and average radial velocity within active regions. Following 150 cycles with 50 *µ*m illumination, the active contracting radius stabilized at ∼30 *µ*m at the periphery, maintaining a contraction speed of ∼0.4 *µ*m/s. With 75 *µ*m illumination, the radius expanded to ∼70 *µ*m after 70 cycles while preserving a comparable speed of ∼0.4 *µ*m/s. For 100 *µ*m illumination, the radius reached ∼50 *µ*m after 30 cycles, again demonstrating a consistent average speed of ∼0.4 *µ*m/s.

**FIG. S15.**
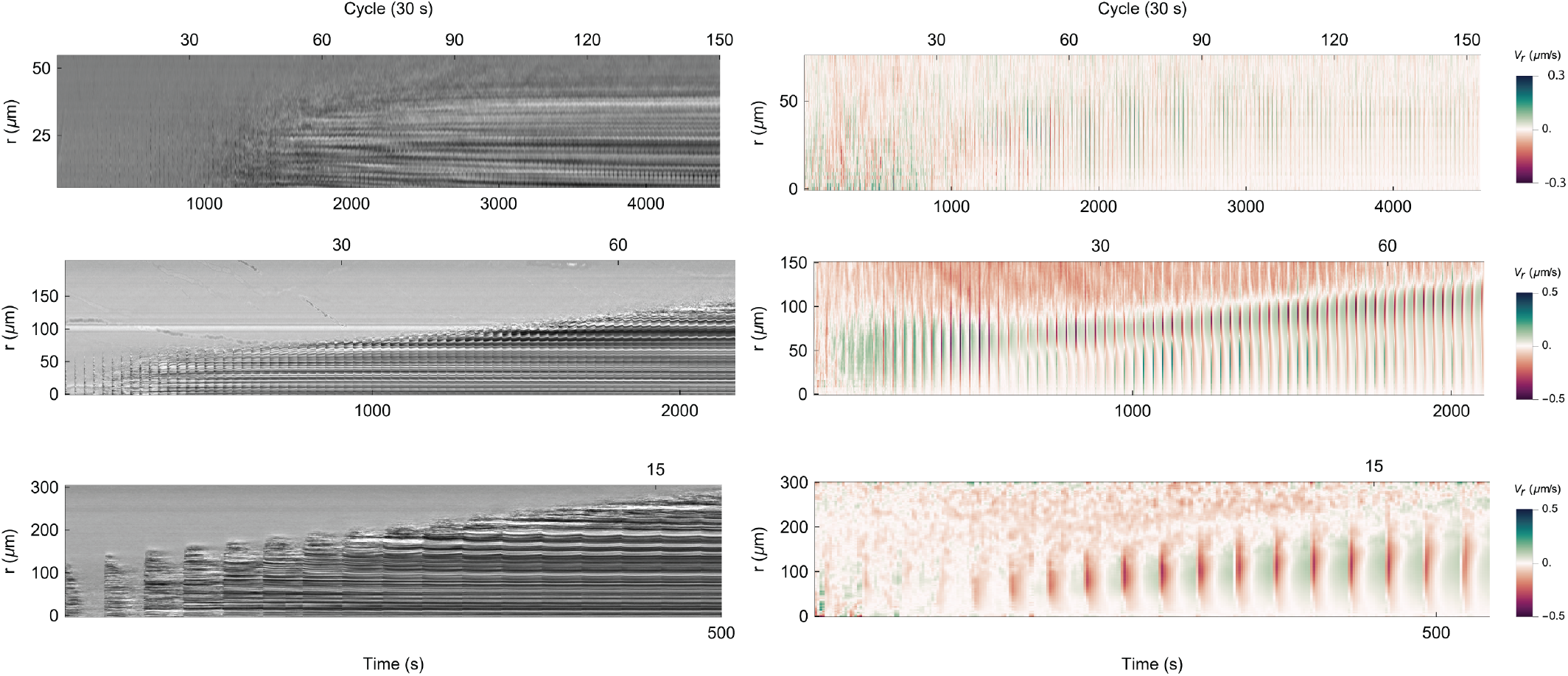
Repeatability tests under the pulsed protocol. *Left* : DIC kymographs for illumination regions of 50 *µ*m (150 cycles), 75 *µ*m (70 cycles), and 100 *µ*m (30 cycles) in diameter. *Right* : Corresponding radial velocity *V*_*r*_ kymographs.

### C. Simulated concentration fields under continuous and pulsed protocols

In Figs. S17 and S18, we show the radial profiles of the concentrations for each chemical species tracked in simulation, for the continuous and pulsed protocols.

### D. Phases during a light pulse

In Figure S19 we plot various physical fields during different phases of the 10^th^ simulated pulse of light.

**FIG. S16.**
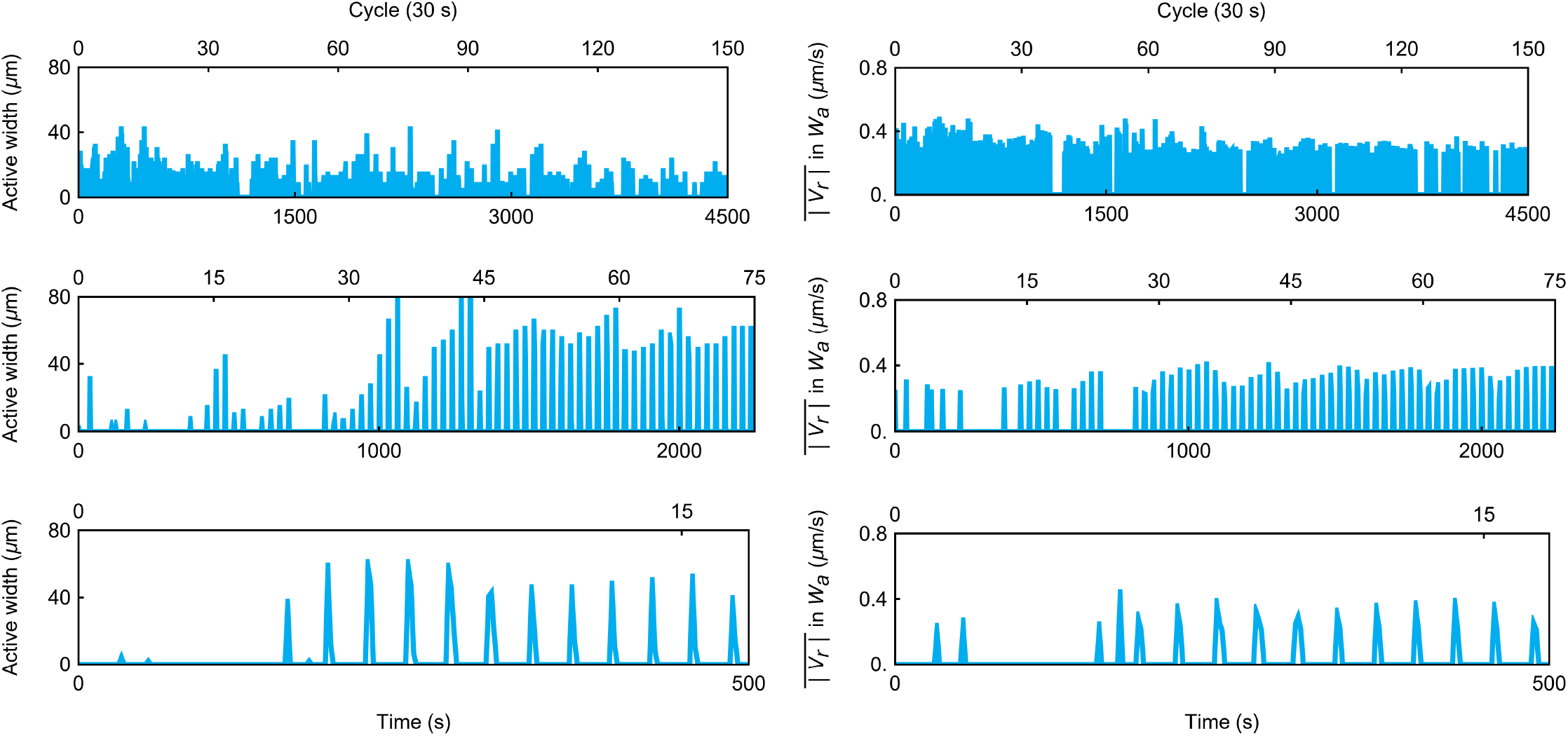
Repeatability of active width generation under the pulsed protocol. *Left* : active width *W*_*a*_ for illumination regions of 50 *µ*m (150 cycles), 75 *µ*m (70 cycles), and 100 *µ*m (30 cycles), defined as the radial extent where |*V*_*r*_ | *>* 0.2 *µ*m/s. *Right* : mean radial velocity 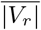 within *W*_*a*_ across cycles.

**FIG. S17.**
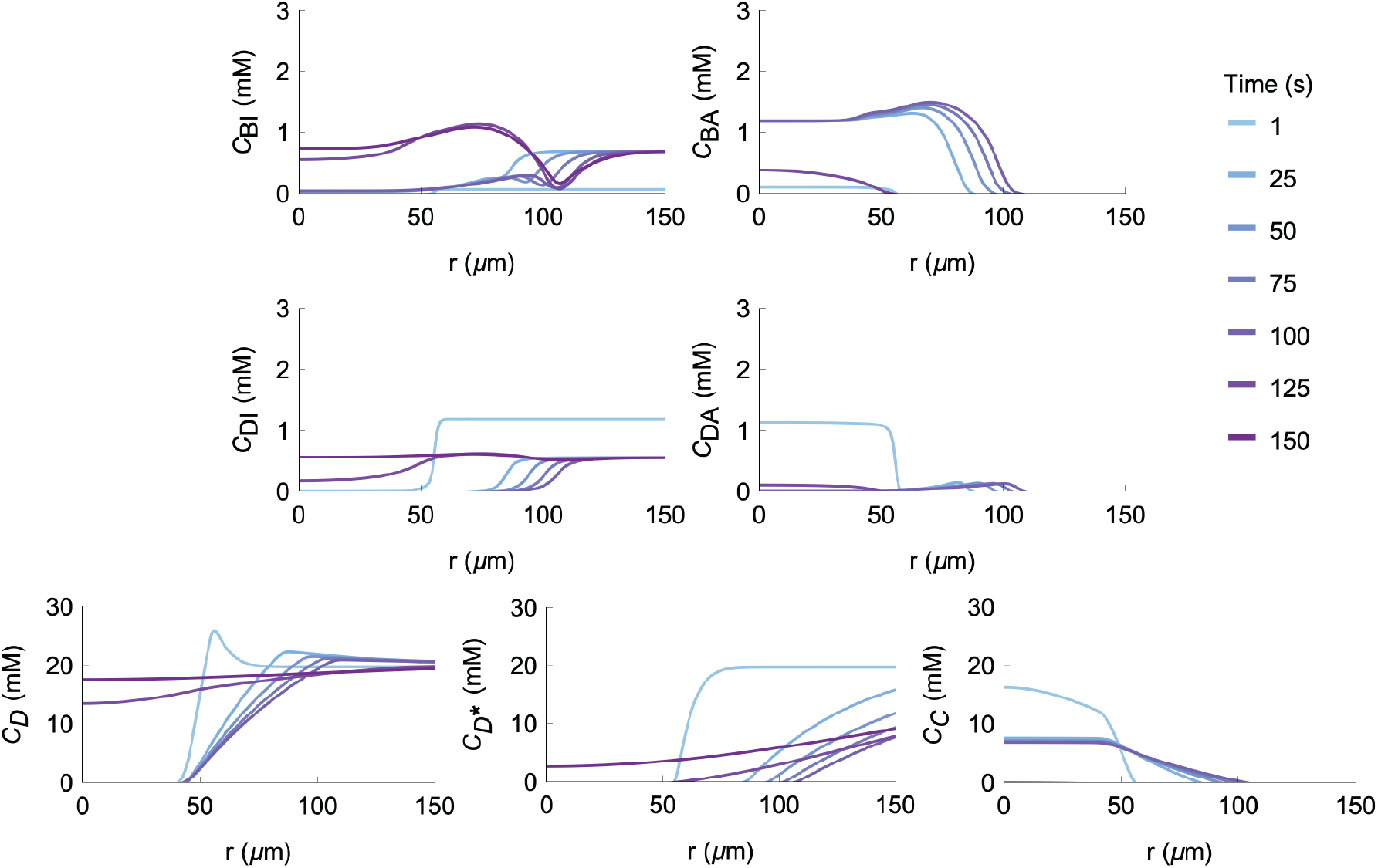
Concentration profiles during the continuous protocol. Simulated concentration profiles for the various chemical species at different time points during the continuous protocol. The light is held on for 100 s and then turned off. The illumination diameter is 75 *µ*m.

**FIG. S18.**
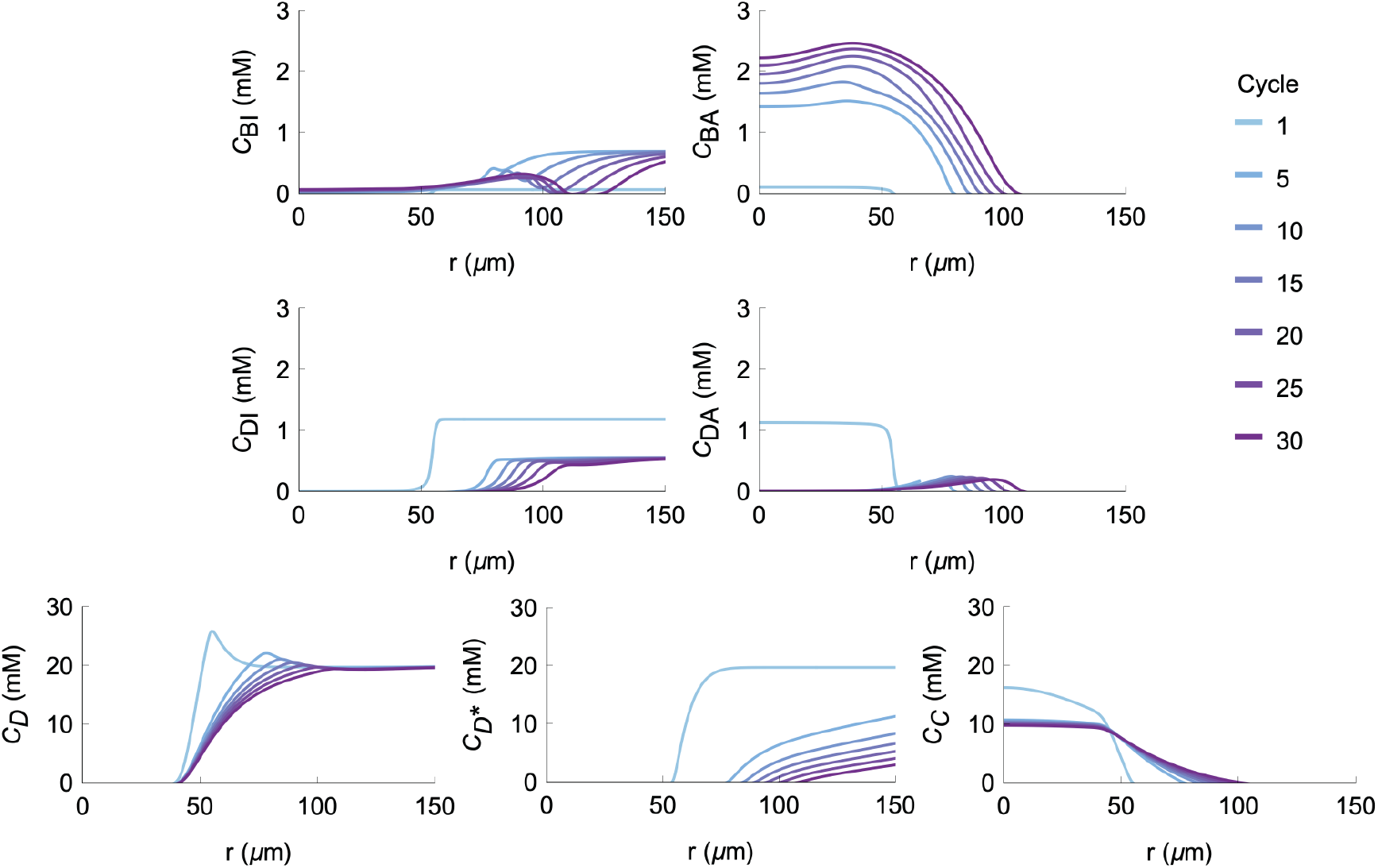
Concentration profiles during the pulsed protocol. Simulated concentration profiles for the various chemical species at different time points during the pulsed protocol. The light is held on for 1 s and then turned off for 29 s for each cycle. The profiles are illustrated at the end of each 1 s light-on pulse. The illumination diameter is 75 *µ*m.

### E. Correlation of network and Ca^2+^ concentration profile

Ca^2+^ underlies network growth and contraciton in the Tcb2 network. Unfortunately in this study we are not equipped to perform quantitative calibration using the ratiometric method to measure Ca^2+^ concentration, and we leave this measurement for future work. Instead, we conducted a quantitative characterization of the spatial distribution at 340 nm using Ca^2+^ indicator rhodamine-2. In Fig. S20, we present experimental fluorescence images immediately after light activation and after 30 s of activation. A persistent bright spot is visible in the illuminated region at both time points. By the later time point, a more diffuse halo of fluorescence is also apparent, representing the diffusion of Ca^2+^ away from the illuminated region. Simulation results, colored to match the grayscale of the experimental images, resemble the experimental images.

### F. Comparison of mechanical contraction and chemical dissociation

In Fig. S21 we highlight the disparate timescales of fast mechanical contraction at the active boundary and slow chemical dissociation of the network.

**FIG. S19.**
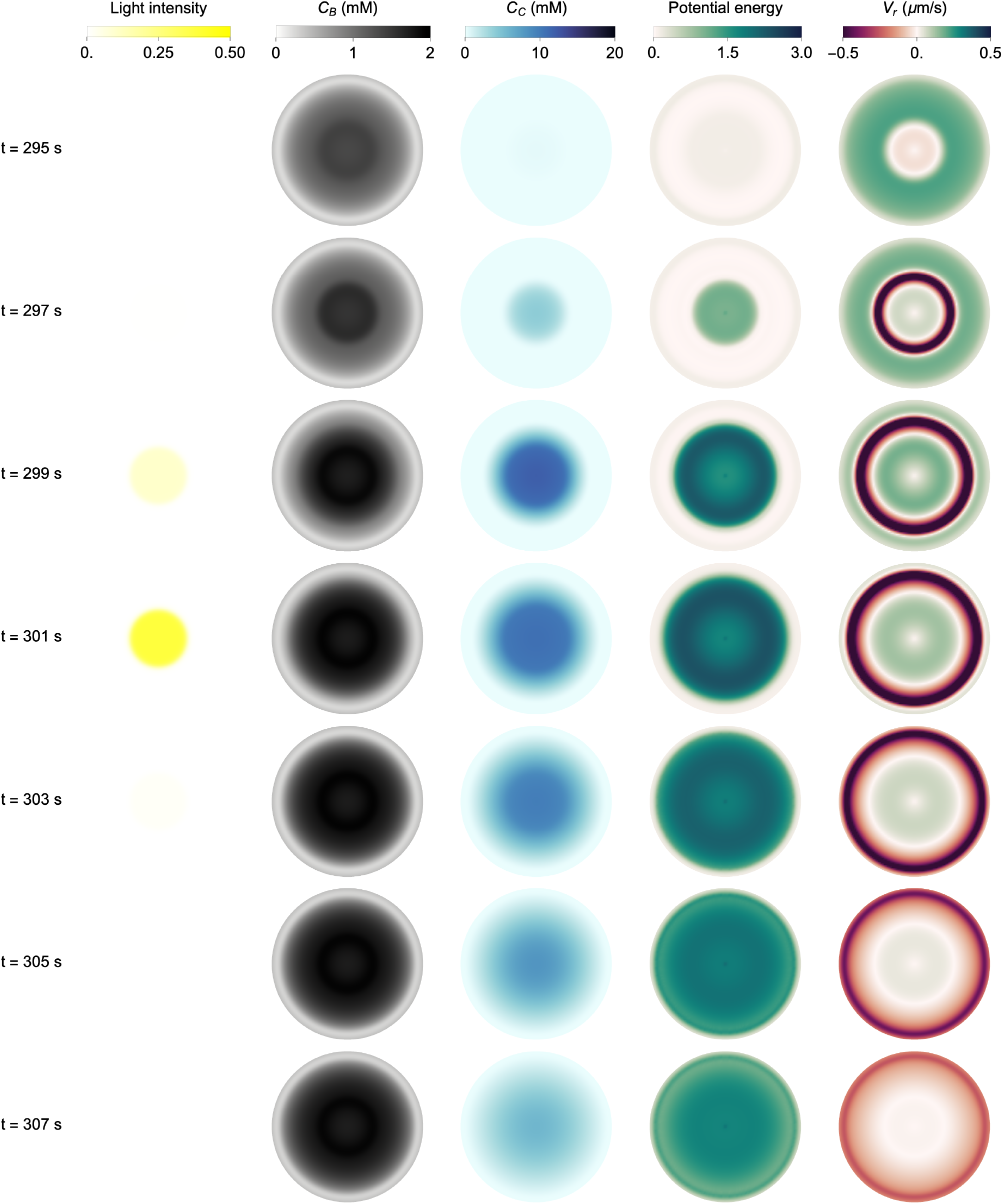
Phases of various fields during a pulse of light. During the 10^th^ simulated pulse of light, various physical fields are plotted at different phases. The mechanical potential energy density is computed as 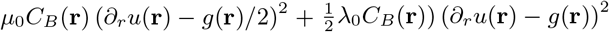. The diameter of each visualization domain is 200 *µ*m, and the diameter of the light pulse is 75 *µ*m.

**FIG. S20.**
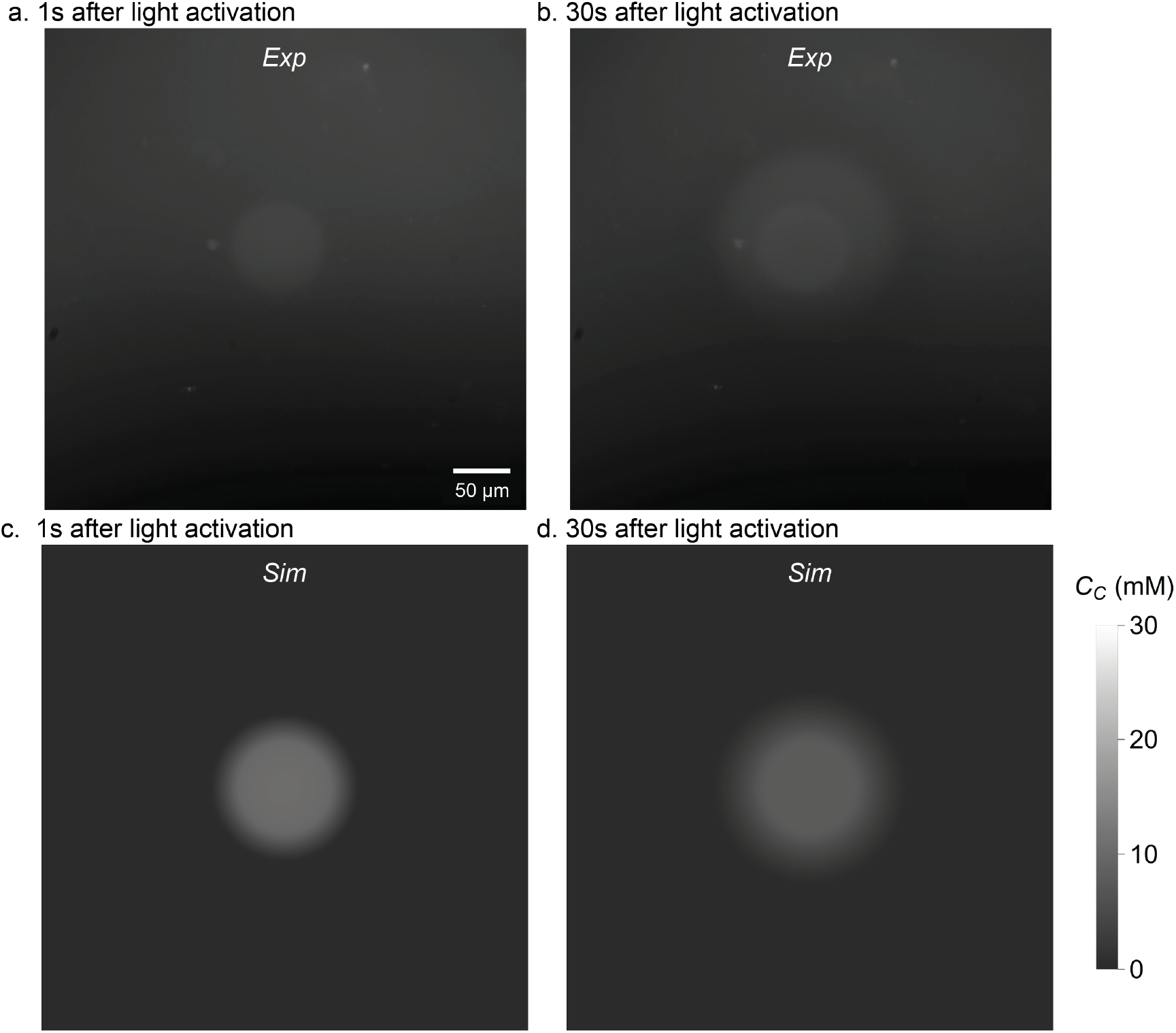
Visualization of light-induced Ca^2+^ release with Ca^2+^ indicator dye rhodamine-2. (a) Image taken immediately after 1 s of light activation, showing a faint fluorescence signal indicating Ca^2+^ release. The illumination diameter is 75 *µ*m. See also SI video Part 3, Section VIII. (b) Image taken after 30 s of light illumination, showing a further spread or diffusion of Ca^2+^. (c) and (d) Corresponding simulation images of the the diffusing Ca^2+^ concentration, colored to match the experimental images.

### G. Network growth requires degradation of chelator

In Fig. S22, we show simulation results indicating that for the network to grow in size over the course of both the continuous and pulsed light protocols, it is necessary that the DMNP-EDTA chelators degrade upon photolysis and release of Ca^2+^. If the chelators are perfectly able to re-uptake Ca^2+^ after photolysis, then the total available pool of Ca^2+^ does not increase in time indefinitely and the system reaches a steady state at finite size. Chelators thus provide a controllable, but not infinitely rechargeable, way to release free Ca^2+^ to the system.

The experimental value of *β* is not known. However, we note that the precise value of the degradation parameter *β* does not appear to have a strong effect on the growth rate of the network, provided that it is not equal to 1, as other steps presumably become rate limiting.

**FIG. S21.**
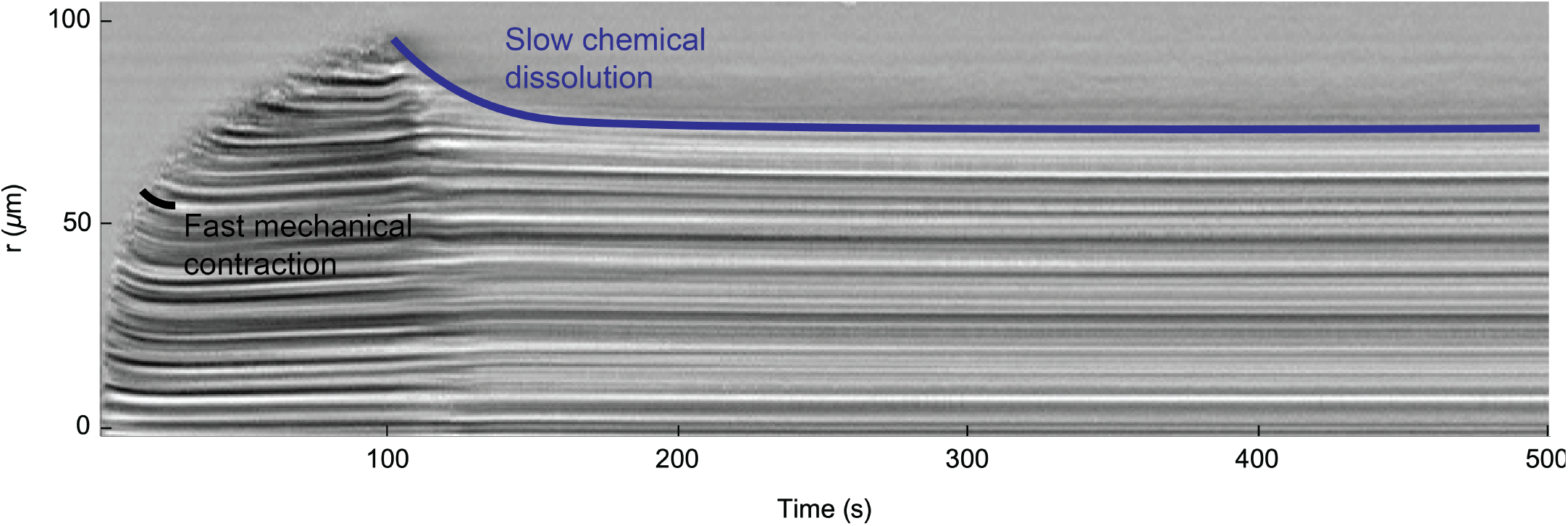
Comparison of mechanical contraction and chemical dissociation. Blow-up of Fig. 2b of the main text, highlighting the kymograph features corresponding to fast mechanical contraction at the active boundary and slow chemical dissolution after the light is turned off.

**FIG. S22.**
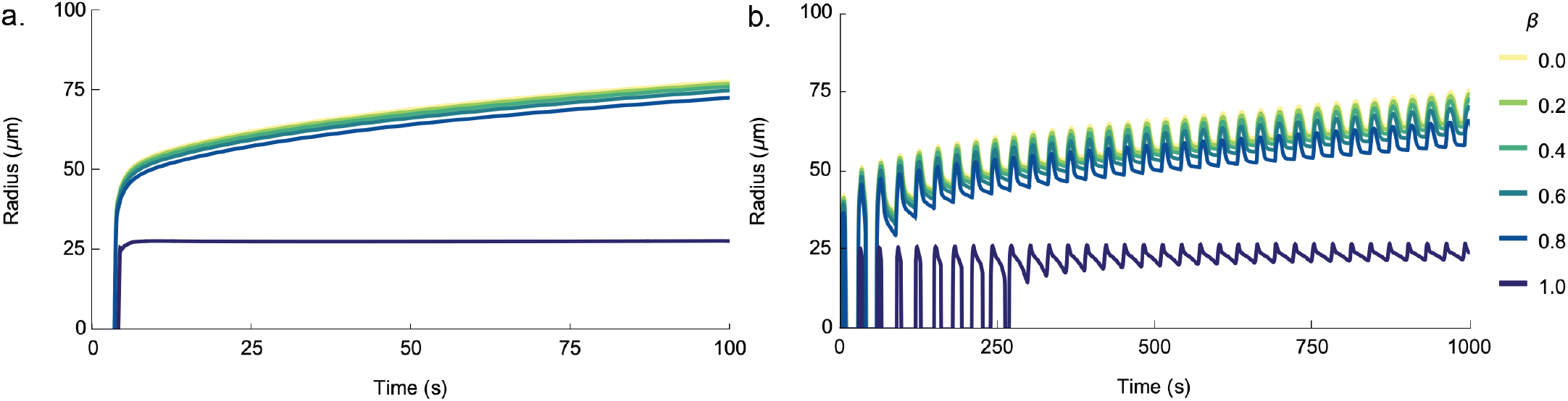
Effect of chelator degradation on network growth. (a) Simulated growth radius as a function of time for various values of the degradation parameter *β*, for the continuous light protocol. For *β* = 1, there is no degradation of chelator upon photolysis and Ca^2+^ release, and for *β* = 0 there is complete degradation. (b) Same as panel a, for the pulsed light protocol.

### H. Comparing the active region in light-on state under pulsed and continuous protocols

In Fig. 3 of the main text we show how the size of the Ca^2+^-active region remains (CAR) large for the pulsed protocol than for the continuous light protocol. In Fig. S23 we quantify this by measuring the radial size of the CAR |*W*_*a*_|, which is radial extent of the part of the system in which the radial velocity *V*_*r*_ is greater than 0.2 *µ*m/s in magnitude. We also average the radial velocity magnitude 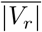 over the CAR. We average over the the time that the light is on time for each protocol, capturing 100 s of light activation in the continuous protocol and 30 pulses (30 s of light-on time) in the pulsed protocol. Simulations semi-quantitatively match the experimental data (S23b).

**FIG. S23.**
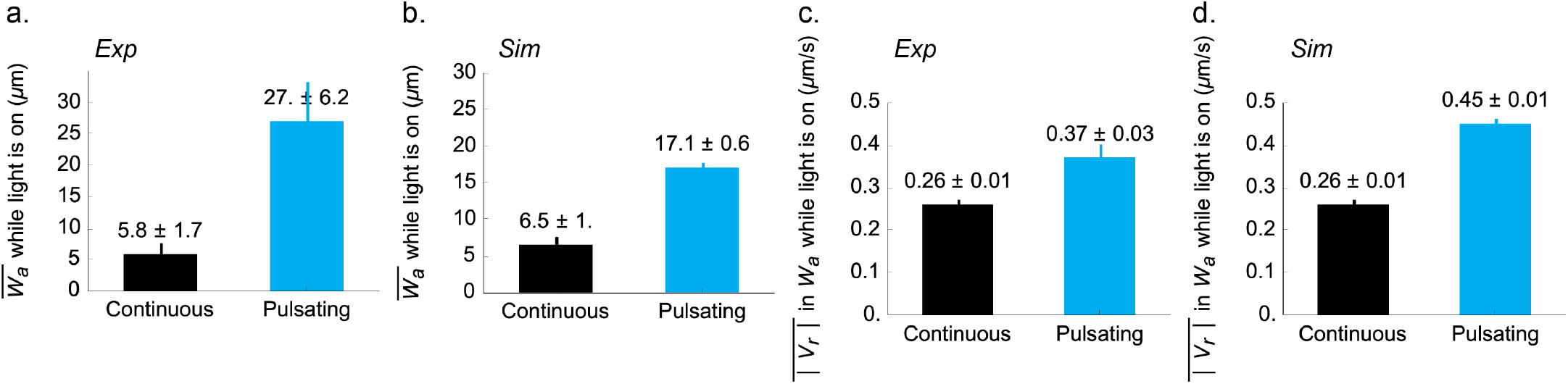
Active widths and mean radial velocities under continuous and pulsed conditions. (a) Measurements of the active width 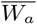 (i.e., radial spatial extent for which |*V*_*r*_ |*>* 0.2 *µ*m) for the continuous and pulsed protocols, using an illumination diameter of 75 *µ*m. Error bars represent the standard deviation aggregated over time and space of the averaging window. The data for the continuous protocol was sampled every second and averaged over the 100 s duration the light was on. The data for the pulsed protocol was sampled every 30 seconds during each 1-second light-on interval, at times 1 s, 31 s, 61 s, …, up to 601 s. (b) Same as panel a, for the simulation. (c) Same as panel a, for the average magnitude of the radial velocity in the active region. (d) Same as panel c, for the simulation.

### I. Varying the frequency of light pulses

Here we explore the effect of varying the frequency of light pulses in the pulsed protocol. In experiments, we carry out a protocol in which the pulse length and cycle length (cf. Fig. S6 above) are scaled by the same factor in a fixed ratio of 1*/*30, shown in Fig. S24a, which we also do in simulations, shown in Fig. S24b. Despite a quantitative difference in the extent of growth between simulation and experiments (discussed above), we observe qualitative agreement on two key features. First, the size of the initial network growth increases with the pulse length. Additionally, following the end of a pulse, the network radius decreases in an apparent biphasic manner, with a quick drop in size followed by a slower decay. The slopes of these two phases of network shrinking do not seem to depend strongly on the pulse length, reflecting instead the chemical kinetics of Tcb2 inactivation and unbinding from the network.

We also test in simulation a protocol in which we fix the pulse length at 1 s and increase the cycle length, shown in Fig. S24c. This makes apparent that the growth of the network using the pulse protocol depends decreases with the cycle length (i.e., the frequency). The network size decays in a roughly linear manner following the cessation of light, and the network will grow faster if this period of decay has a shorter duration. The increase in network size upon each cycle of light is roughly, but not exactly, the same across cycle number and cycle length conditions. Given these results, our choice of default pulse length and cycle length as 1 s and 30 s respectively is a practical choice that makes the experimental run time more manageable without qualitatively changing the behavior we expect with longer cycle lengths.

### J. Varying the ratio of DMNP-EDTA to Tcb2

Here we discuss the effects of varying the ratio of DMNP-EDTA to Tcb2 in the experimental solution. We expect that increasing the concentration of free DMNP, which are available to take up the Ca^2+^ ions released by Ca^2+^-bound DMNP-EDTA upon photolysis, reduces the available supply of Ca^2+^ to bound to Tcb2 and thereby hinders growth of the network. Indeed, we find from repeated trials that networks fail to nucleate and grow even with the large illumination diameter (100 *µ*m) when we double the concentration of free DMNP-EDTA in solution. One such example is shown in Fig. S25, which we compare to the default concentrations used elsewhere. Simulations qualitatively support this result, showing a reduced growth rate with greater DMNP-EDTA concentrations than at normal conditions. However, in simulations the nucleation is not prevented from occurring at high DMNP-EDTA concentrations, indicating that further refinement of the nucleation model is needed in future work. These results suggests that a somewhat precise balance of chemical concentrations is needed to prevent the system from being overwhelmed with chelators that soak up the available Ca^2+^.

**FIG. S24.**
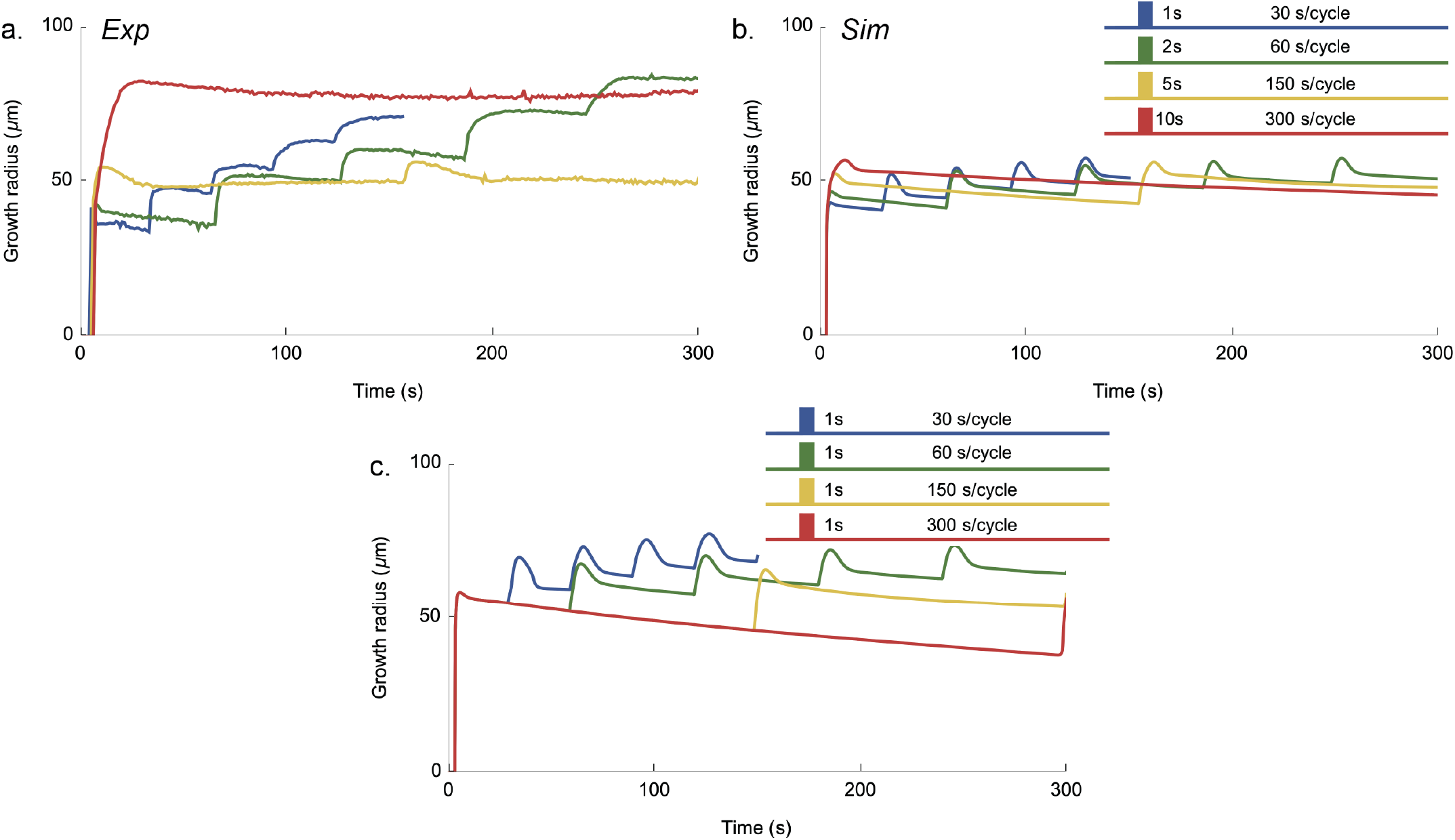
Network growth dynamics with varying light pulse frequency. (a) Experimental growth radius as a function of time for various pulse protocols. See panel b for the color legend. (b) Simulated growth radius as a function of time for various pulse protocols. The color legend indicates the pulse length and cycle length for each protocol. (c) Simulated growth radius as a function of time for various pulse protocols in which the pulse length is fixed at 1 s.

### K. Boundary accumulation depends on diffusion of Tcb2

Here we study the accumulation of bound Tcb2 near the periphery of the network. We used 1.01 *µ*M Fluoresbrite Polychromatic Red Microspheres (PolyScience 18660-5) for boundary accumulation experiment. In Fig. S26 we show experimental images of embedded fluorescent particles which report on the local density of the system. After light activation, we observe that that the formed network has radially structured density profile, with signatures of accumulation and depletion near the network boundary.

**FIG. S25.**
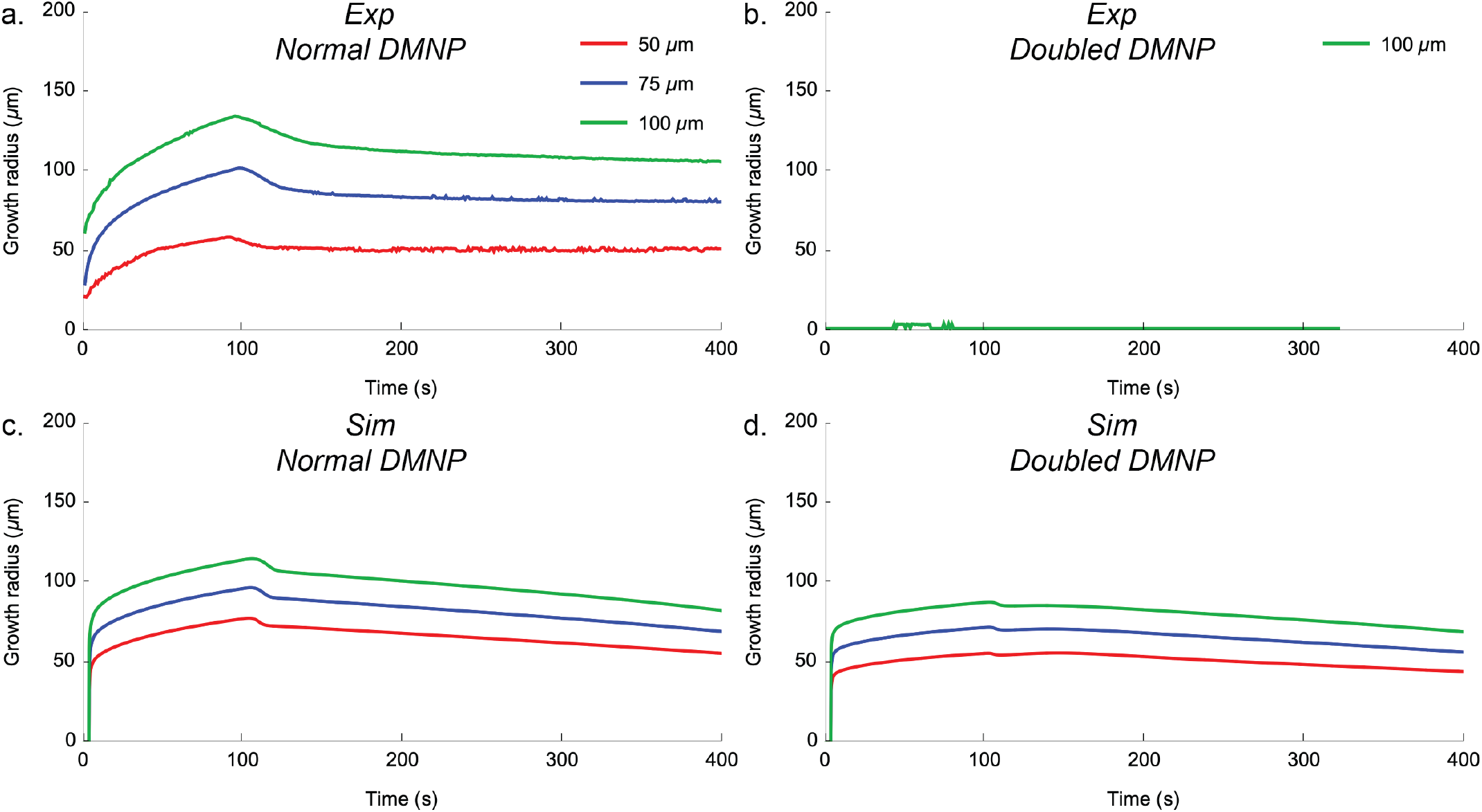
Network growth dynamics with varying concentrations of Tcb2 and DMNP. (a) and (b) Experimental growth radius as a function of time for various illumination diameters, at a normal (used elsewhere in the paper) and doubled DMNP-EDTA concentrations. (c) and (d) Same as panels a and b, for simulations.

**FIG. S26.**
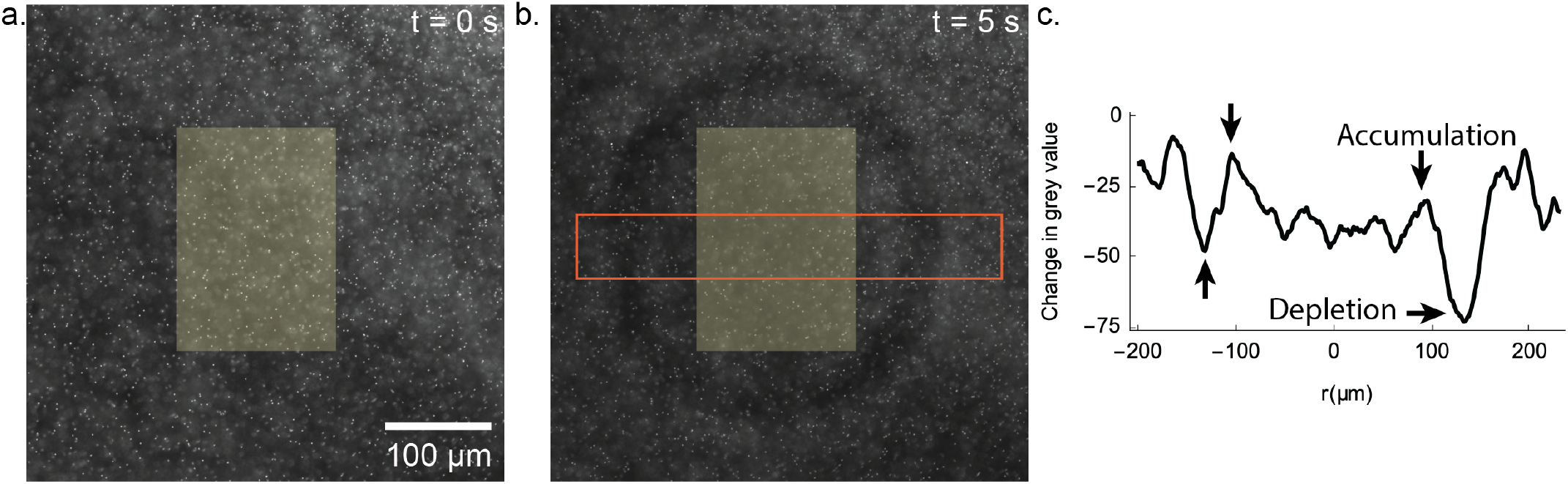
Boundary effects. (a) Fluorescent image of a Tcb2 network before light activation with embedded 1 *µ*m beads. Same as panel a, after 5 seconds of light activation in the labeled yellow region. (c) Average change after 5 s in grey values within the the horizontal orange strip in the middle of panel b, with a 70 *µ*m height and 400 *µ*m width.

In Fig. S27 we demonstrate in simulation that the radial density structure of bound Tcb2 near the periphery of the network is largely controlled by the diffusion of Tcb2. In the network interior, there is little diffusing Tcb2, setting up a concentration gradient compared with the exterior. Tcb2 then diffuses down the gradient and binds to the network. As it encounters the peripheral region of the network, it starts to bind first there before it diffuses to further inside. This leading to a bump of density of bound Tcb2 at the periphery. Additionally, there is a trough in density just exterior to the network because of local depletion via diffusion towards the network. As we turn down the diffusion constant of Tcb2, we see that the profile becomes more homogeneous as expected.

**FIG. S27.**
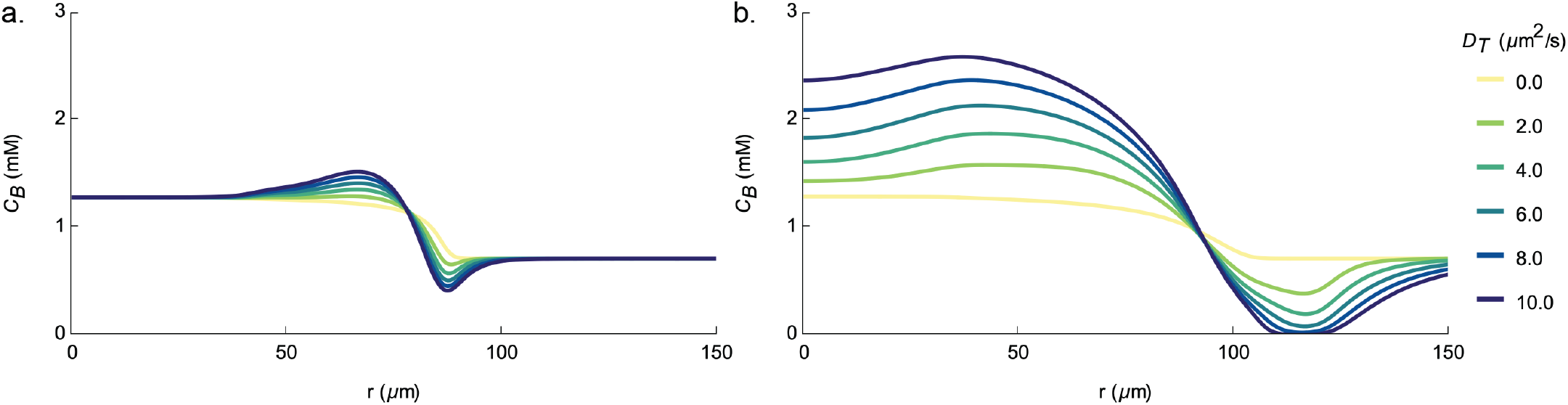
Effect of Tcb2 diffusion on boundary accumulation. (a) Simulated profile of the bound Tcb2 concentration after 30 s of the continuous light protocol for varying values of the Tcb2 diffusion constant *D*_*T*_. (b) Same as panel a, for the 30^th^ cycle of the pulsed light protocol.

### L. Spatially dependent particle transport

In Fig. 5a and b of the main text, we demonstrate that lipid particles are displaced in the presence of a growing Tcb2 network, exhibiting complex dynamics: some particles are pulled inward, others are pushed outward, and some experience sequential push-then-pull motion. This behavior primarily depends on each particle’s initial radial distance from the illumination zone. Particles close to the zone are predominantly pulled inward, while those at greater distances are predominantly pushed outward. We illustrate this radial dependence in Fig. S28a, where we plot the radial motion of each particle including its initial offset. We successfully transported three types of particles: polystyrene beads (diameter: 10–20 *µ*m), liposomes, and lipid particles, with their displacement profiles shown in Fig. S28. The pulling effect arises from mechanical contraction of the Tcb2 network, while the pushing effect likely results from steric interactions between particles and the expanding network periphery. Particles closer to the illumination zone become trapped within the forming network and experience primarily pulling forces, whereas those at greater distances encounter the slowly expanding network front and experience primarily pushing forces. Note that current simulations do not incorporate steric interactions between the Tcb2 network and particles, thus modeling only the pulling effect via network contraction.

### M. Formation of a peripheral network

Although the Tcb2 network grows at high density primarily within the illumination region, the rapid diffusion of Ca^2+^ outward from the illumination region also sets up a lower-density peripheral network. This rapidly forming network can then sterically interact with embedded particles, such as the liposome in Fig. 5c of the main text, on the time scale of seconds. We highlight the growth of this peripheral network in Fig. S29.

**FIG. S28.**
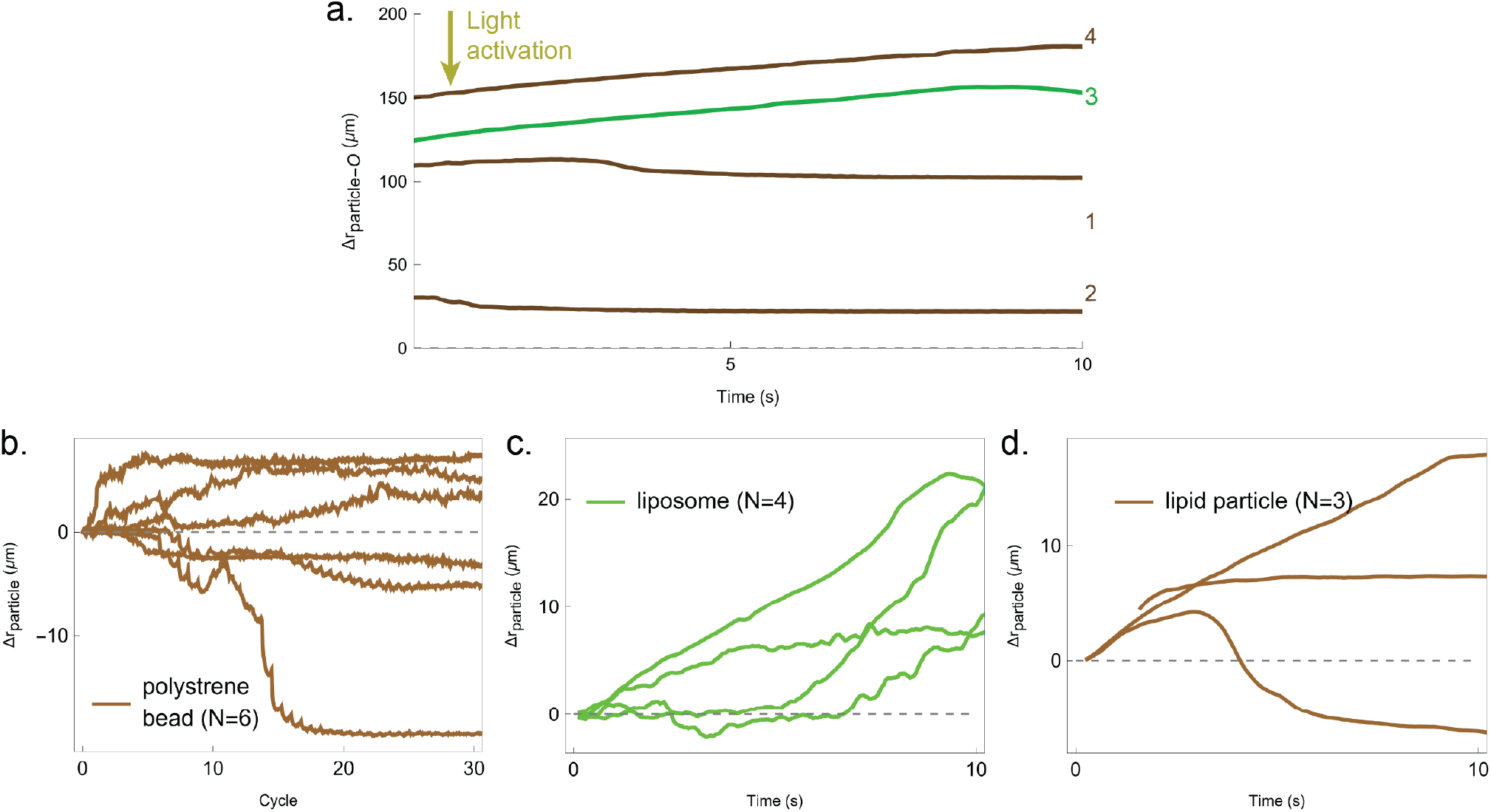
Dependence of pushing and pulling on initial radial distance. (a) This plot corresponds to Fig. 5c of the main text, replotted so that the initial radial distance of each object from the origin is shown. Displacements of (b) polystyrene beads (N=6) under pulsed light (1s on, 29s off), (c) liposomes (N=4), and (d) lipid particles (N=3) under continuous light.

**FIG. S29.**
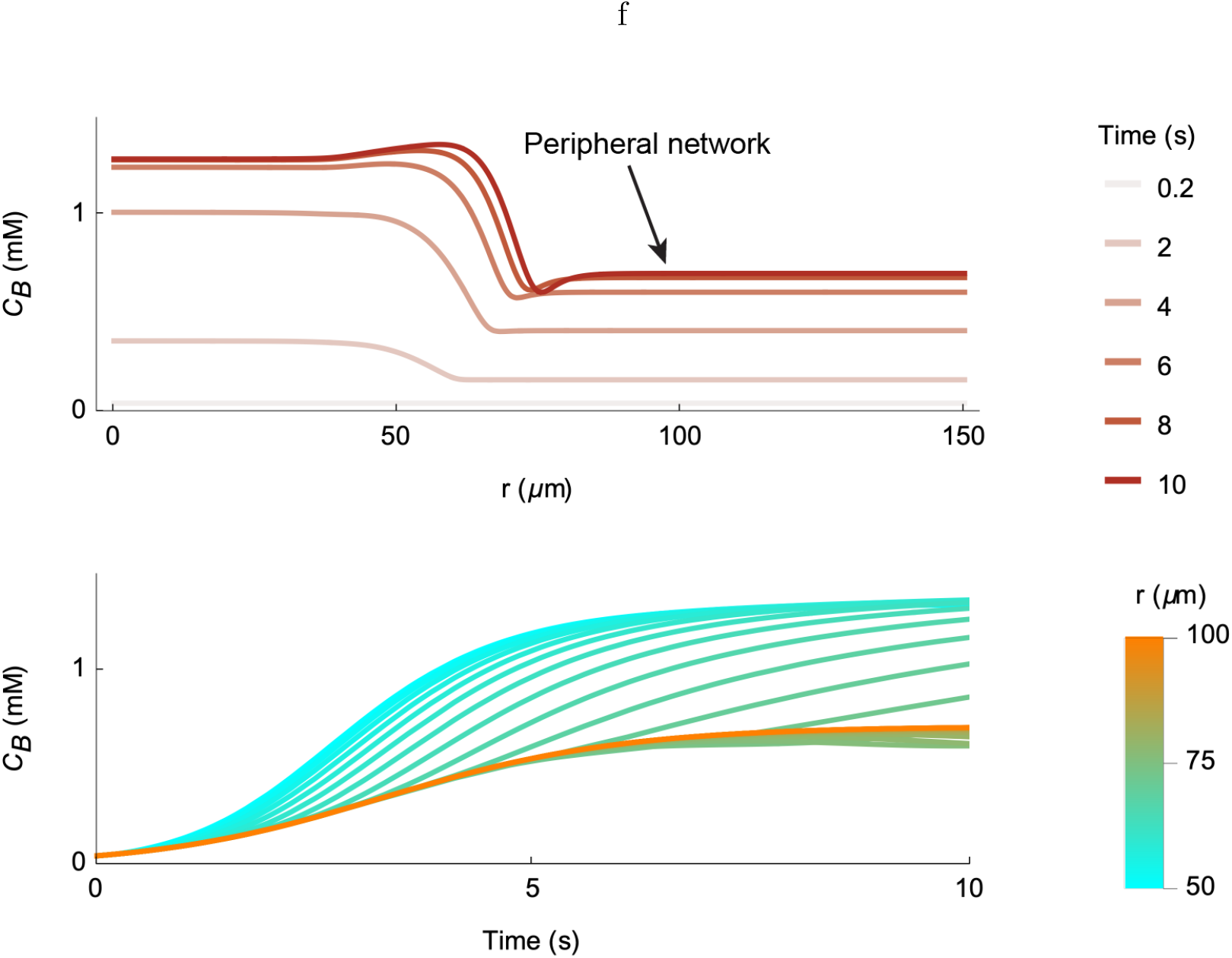
Growth of the peripheral network. *Top*: The radial profile of bound Tcb2 concentration at different times during the early part of a continuous light protocol with illumination diameter of 75 *µm*. The peripheral network, at lower density and further from central illuminated zone, is indicated. *Bottom*: An alternate view showing time profiles at different radial slices, highlighting that the main and peripheral networks grow on similar timescales, though to different densities.

## III. SUPPLEMENTARY VIDEO CAPTIONS

- Part 1 Section I video: Ca^2+^ diffuses into a sandwich-structured chamber from left to right, inducing a gel-like transition in the Tcb2 solution and contraction is happening along the boundary.
- Part 1 Section II video: Tcb2 network formation, imaged using DIC, for a star-shaped DMD pattern.
- Part 1 Section II video: Dynamic Tcb2 network formation by a 0.125 Hz clockwise cyclic light pattern.
- Part 1 Section III video: Tcb2 network formation under illumination with a 75 *µ*m diameter for 100 seconds. Right: Corresponding experimental analysi PIV image.
- Part 2 Section III video: Corresponding Tcb2 simulation showing light intensity, Tcb particle radial velocity*v*_*r*_, bound inactivated Tcb2 (*C*_*BI*_), bound activated Tcb2 (*C*_*BA*_), diffusing Ca^2+^ (*C*_*C*_), DMNP with Ca^2+^ (*C*_*D*_), DMNP-EDTA without Ca^2+^ (*C*_*D*∗_), diffusing inactivated Tcb2 (*C*_*DI*_) and diffusing activated Tcb2 (*C*_*DA*_) under illumination with a 75 *µ*m diameter for 100 seconds.
- Part 2 Section III video: Concentration profiles of all compounds as a function of radial position with a 75 *µ*m diameter for 100 seconds.
- Part 2 Section IV Video: Tcb2 network formation under illumination with a 75 *µ*m diameter, using a 1-second light-on and 29-second light-off cycle. Right: Corresponding experimental PIV analysis image.
- Part 2 Section IV Video: Simulation of Tcb2 dynamics under a 1-second light-on and 29-second light-off cycle.
- Part 2 Section IV Video: Concentration profiles of all compounds as a function of radial position under a 1-second light-on and 29-second light-off cycle.
- Part 2 Section V video: Embedded fluorescent 1 *µ*m beads within the Tcb2 network under light activation in the labeled yellow region.
- Part 2 Section VI Video: 150 cycles repeatability test with 50 *µ*m diameter under a 1-second light-on and 29-second light-off cycle.
- Part 3 Section VII Video: Tcb2 network facilitates the movement of liposomes and lipid particles with a designed light pattern.
- Part 3 Section VII Video: Tcb2 network facilitates the movement of 10–20 *µ*m polystyrene beads with multi-pattern light activation.
- Part 3 Section VIII Video: Rhodamine-2 Ca^2+^ indicator showing the Ca^2+^ profile during Tcb2 network formation.

